# A dolabralexin-deficient mutant provides insight into specialized diterpenoid metabolism in maize (*Zea mays*)

**DOI:** 10.1101/2022.07.21.500061

**Authors:** Katherine M. Murphy, Tyler Dowd, Ahmed Khalil, Si Nian Char, Bing Yang, Benjamin J. Endelman, Patrick M. Shih, Christopher Topp, Eric A. Schmelz, Philipp Zerbe

**Author notes:** To whom correspondence should be addressed: Philipp Zerbe. Current address: Donald Danforth Plant Science Center, St. Louis, MO, USA.

## Abstract

Two major groups of maize (*Zea mays*) specialized metabolites, termed kauralexins and dolabralexins, serve as known or predicted diterpenoid defenses against pathogens, herbivores, and other environmental stressors. To consider physiological roles of the recently discovered dolabralexin pathway, we examined dolabralexin structural diversity, tissue-specificity, and stress-elicited production in a defined biosynthetic pathway mutant. Metabolomics analyses support a larger number of dolabralexin pathway products than previously known. We identified dolabradienol as a previously undetected pathway metabolite and characterized its enzymatic production. Transcript and metabolite profiling showed that dolabralexin biosynthesis and accumulation predominantly occurs in primary roots and shows quantitative variation across genetically diverse inbred lines. Generation and analysis of CRISPR-Cas9-derived loss-of-function Kaurene Synthase-Like 4 (*Zmksl4*) mutants demonstrated dolabralexin production deficiency, thus supporting ZmKSL4 as the diterpene synthase responsible for the conversion of geranylgeranyl pyrophosphate precursors into dolabradiene and downstream pathway products. *Zmksl4* mutants further display altered root-to-shoot ratios and root architecture in response to water deficit, consistent with an interactive role of maize dolabralexins in plant vigor during abiotic stress.

## Introduction

Plant terpenoids are a structurally diverse metabolite class that serve critical functions in growth, defense, and environmental adaptation (Gershenzon and Dudareva, 2007). Beyond widely conserved gibberellin (GA) phytohormones with key roles in plant growth, the vast majority of diterpenoids are often species-specific and serve specialized functions in stress responses during microbial attack, herbivory, as well as abiotic stress (Schmelz et al., 2014; Tholl, 2015; Block et al., 2019; Murphy and Zerbe, 2020; Li et al., 2021; Zhang et al., 2021).

In maize (*Zea mays*), two groups of bioactive diterpenoids, termed kauralexins (KX) and dolabralexins (DX), have been identified as serving defense-related functions (Murphy and Zerbe, 2020). KX are acidic diterpenoids that feature an *ent*-isokaurene-derived backbone with distinct levels of oxidation and reduction (Ding et al., 2019). KX metabolites function as local defenses produced at nearly any site of challenge in response to pathogen infection by, for example, *Fusarium verticillioides, F. graminearum, Rhizopus microsporus, Cochliobolus heterostrophus,* and *Colleototrichum graminicoloca*, as well as in response to water deficit and below-ground oxidative stress (Schmelz et al., 2011; Schmelz et al., 2014; Christensen et al., 2018; Ding et al., 2019). Aligned with their stress-elicited formation, KX display potent antifungal activity *in vitro* and *in planta*, as well as anti-feedant activity against the European corn borer (*Ostrinia nubilalis*) (Schmelz et al., 2011; Vaughan et al., 2015; Ding et al., 2019). Increased pathogen susceptibility of both the KX-deficient *ent*-copalyl pyrophosphate synthase (*ent*-CPS) mutant termed *Anther Ear 2* (*Zman2*) and the *Kaurene Synthase-Like 2* (*Zmksl2*) mutant demonstrated significant impacts of diterpenoids on maize stress resilience (Harris et al., 2005; Vaughan et al., 2015; Christensen et al., 2018; Ding et al., 2019).

Recently, DX metabolites were discovered as an additional group of maize diterpenoids that are structurally distinct from KX, featuring a dolabradiene scaffold with oxygenations, often including a C-15,16 epoxide (Fig. 1) (Mafu et al., 2018). While the physiological functions of DX remain unknown, metabolite and transcript profiling of the DX pathway showed inducible DX formation in maize roots following infection by *F. graminearum* and *F. verticillioides*, as well as below-ground oxidative stress (Mafu et al., 2018). DX metabolites have been found to accumulate up to 225 µg g^-1^ fresh weight (FW) in mature maize roots infected with *F. graminearum*, compared to 9 µg g^-1^ FW KX in the same tissues (Mafu et al., 2018). Moreover, select DX display potent antifungal efficacy *in vitro* against highly evolved *Fusarium* pathogens in maize (Mafu et al., 2018). Collectively, these findings are consistent with an important role of DX in below-ground stress responses.

**Figure 1.**
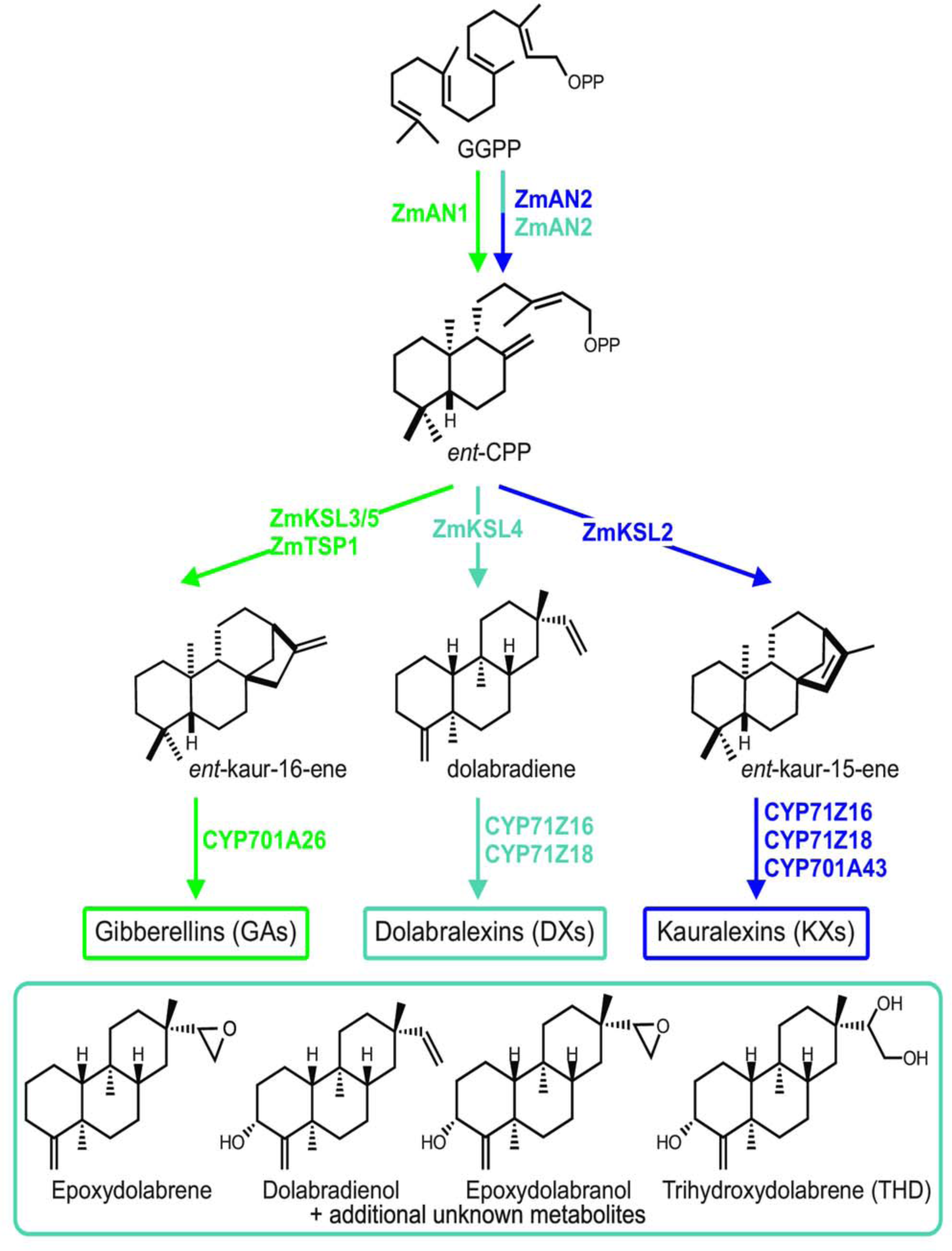
Maize diterpenoid biosynthetic network. Gibberellin (GA) biosynthesis (green) and specialized metabolite pathways for dolabralexins (DX) (blue) and kauralexins (KX) (purple) utilize the same central precursor. Known DX structures are shown in the bottom blue box. GGPP, geranyl geranyl diphosphate; KSL, Kaurene Synthase-Like; CPP, copalyl diphosphate; An, Anther Ear.

Despite structural and functional distinctions, GA, KX, and DX metabolites derive from a common precursor, *ent*-copalyl pyrophosphate (*ent*-CPP) that is formed by two catalytically redundant class II diterpene synthase (diTPS) enzymes, Anther ear 1 and 2 (ZmAn1 and ZmAn2), in the *ent*-CPS family (Fig. 1) (Bensen et al., 1995; Harris et al., 2005; Schmelz et al., 2011; Christensen et al., 2018; Mafu et al., 2018; Murphy et al., 2018). Gene expression, along with genetic and phenotypic studies of *Zman1* mutants, demonstrated that ZmAn1 functions in GA biosynthesis (Bensen et al., 1995). In contrast, the *Zman2* mutant displays no GA-deficiency phenotype (Harris et al., 2005). Multi-functional roles of ZmAn2 and derived KX and DX metabolites in biotic and abiotic stress responses are supported by *Zman2* mutant phenotypes such as deficiencies in both KX and DX, increased susceptibility to fungal infection and water deficit, and altered rhizosphere microbiomes (Harris et al., 2005; Vaughan et al., 2014; Vaughan et al., 2015; Christensen et al., 2018; Mafu et al., 2018; Ding et al., 2019; Murphy et al., 2021).

Downstream of *ent*-CPP, the KX and DX pathways bifurcate through the activity of two class I diTPSs, ZmKSL2 and Kaurene Synthase-Like 4 (ZmKSL4), respectively. ZmKSL2 converts *ent*-CPP into *ent*-iso-kaurene en route to KX, whereas ZmKSL4 transforms *ent*-CPP into the DX precursor dolabradiene (Mafu et al., 2018; Ding et al., 2019) (Fig. 1). Functional modification of these hydrocarbon scaffolds through cytochrome P450 monooxygenases (P450s) of the CYP701 (ZmCYP701A43) and CYP71 (ZmCYP71Z16, ZmCYP71Z18) families, along with a steroid 5α- reductase termed Kauralexin Reductase 2 (ZmKR2), then largely defines the structural diversity and bioactivity of KX and DX metabolites (Fig. 1, gene IDs listed in Supplemental Table 1). In addition, CYP71Z16 and CYP71Z18 exhibit expansive substrate promiscuity and function in the endogenous biosynthesis of diverse acidic sesquiterpenoid phytoalexins termed zealexins, highlighting an interconnected and modular pathway network for producing distinct defense metabolite families (Mao et al., 2016; Mafu et al., 2018; Ding et al., 2019; Ding et al., 2020). Two additional substrate-promiscuous P450s, Kaurene Oxidase (KO) 1 (ZmCYP701A26) and ZmKO2 (ZmCYP701A43), are tandemly arrayed on the same chromosome. While ZmKO1 exhibits activity in the GA pathway, ZmKO2 biochemically responsible for highly oxidized KX (Mafu et al., 2016; Mao et al., 2017; Ding et al., 2019).

In this study, we expand the known diversity of DX metabolites and define the first dedicated DX pathway branch point. Gene expression studies and metabolite profiling show DX accumulation primarily in roots with context-specific constitutive production when plants are grown in the field or field-collected soils. Genetic studies clarify that ZmKSL4 is the sole diTPS responsible for DX biosynthesis. Analysis of a DX-deficient *Zmksl4* maize mutant demonstrates reduced plant biomass and an altered root system architecture, particularly under water deficit conditions, supporting non-redundant roles with the parallel KX pathway branch.

## Results

### *Zmksl4* mutants display deficient DX production *in planta*

To investigate the physiological relevance of DX in maize that is independent from KX, we generated a CRISPR-Cas9-enabled *Zmksl4* mutant through *Agrobacterium*-mediated transformation of the Hi-II maize hybrid line using a guide RNA targeting specifically *ZmKsl4* (*kaurene synthase-like 4*) (Char et al., 2016). Seed of *Zmksl4*, homozygous for a one base pair insertion (Fig. 2A), and its isogenic wild type (WT) sibling, both having the *Cas9* removed through backcrossing to B73 and bulked prior to use in this study. Earlier studies showing abiotic stress inducibility of diterpenoid metabolism and increased drought susceptibility of the KX- and DX-deficient *Zman2* mutant suggested interactive roles in the maize water deficit response (Vaughan et al., 2014; Vaughan et al., 2015). To specifically consider this hypothesis in the context of DX, WT and *Zmksl4* plants were grown in fabric pots outdoors in maize field soil with controlled drip irrigation. Water deficit was imposed on half of the plants by limiting water for six consecutive days at either one-month-old (vegetative stage 5, V5) or two-month-old (vegetative stage 12, V12) at which time plants were assessed for root and shoot fresh weights, and leaf and root samples were collected for downstream analysis (Fig. 2B). No large-scale visual phenotypic differences were observed between WT and *Zmksl4* under control or water deficit conditions and all plants exhibited characteristic leaf curling upon water deficit (Fig. 2B).

**Figure 2.**
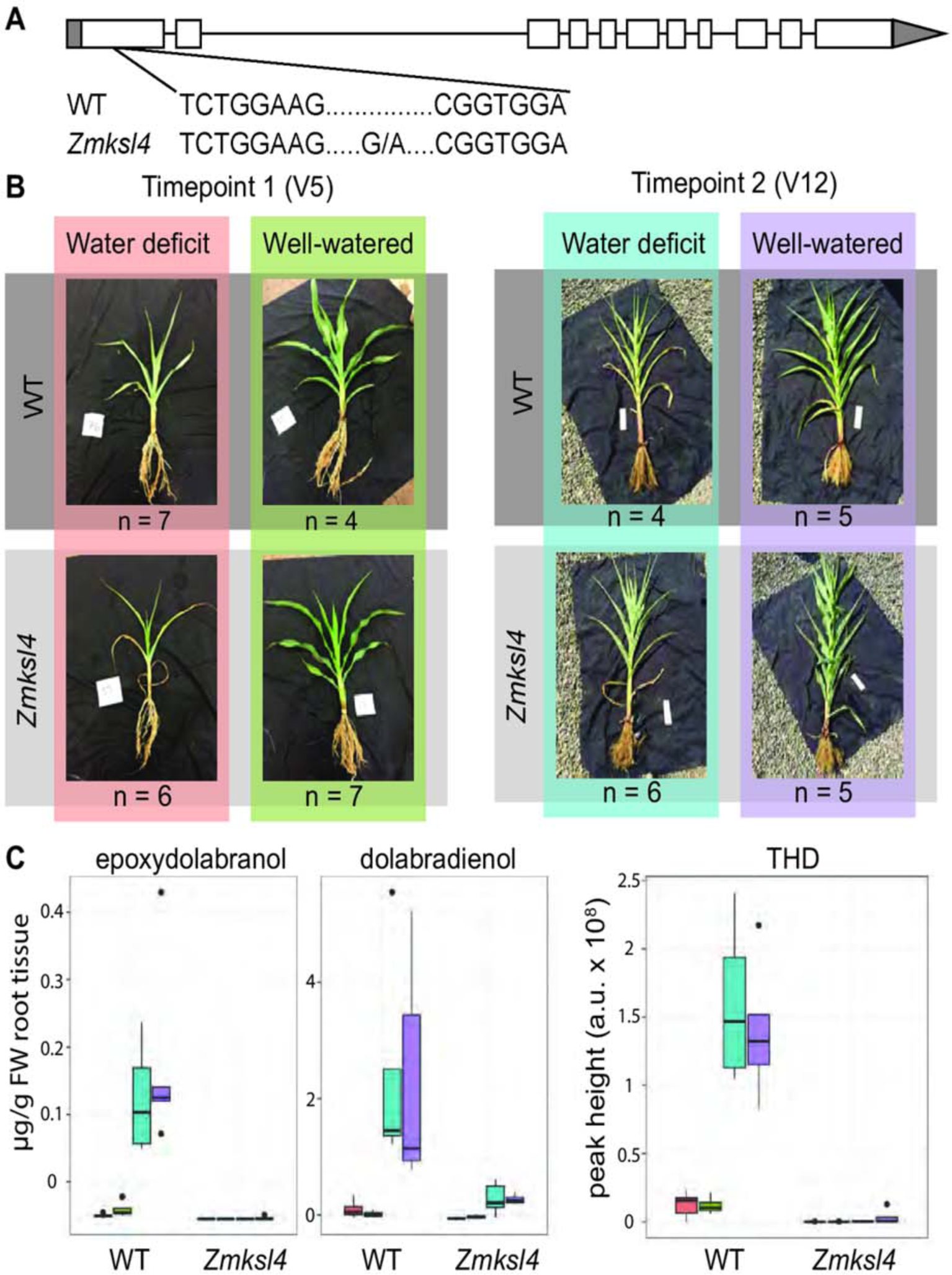
*Zmksl4* maize mutant lacks DX metabolites. (A) *Zmksl4* contains a 1 bp insertion in the ZmKsl4 gene, causing a frame-shift mutation. (B) Experimental design used in this study. WT (dark grey) and *Zmksl4* (light grey) plants were assessed for metabolite and transcript abundances at vegetative stage (V) V5 and V12. Water-deficit and well-watered were analyzed for each respective developmental stage and genotype. Photos are representative plants from each sample type. Biological replicates (n) listed for each sample type. Sample types (particular timepoint and water status) correspond to plot colors throughout this study. (C) LC-MS/MS abundances of DÅs for which standards were available. Letters represent significantly different abundance. P < 0.01 (dolabradienol), p < 0.001 (epoxydolabranol or THD). Bar colors in (C) correspond to sample types colored in (B).

Targeted positive and negative mode Electrospray Ionization (ESI) liquid chromatography mass spectrometry (LC-MS/MS) analysis using enzymatically produced DX or tissue-purified standards, including the major DX metabolites epoxydolabranol and trihydroxydolabrene (THD), respectively, showed no detectable levels in roots or leaves of *Zmksl4* mutant plants (Fig. 2C; metabolite atlas in Supplemental Fig. 1). Both epoxydolabranol and THD were found at low levels in WT V5 roots, and significantly greater in WT V12 roots;here was no significant difference in response to water status (Fig. 2C). Epoxydolabrene was not detectable in any samples, including the purified standard, due to insufficient ionization for ESI LC-MS/MS analysis. THD could not be absolutely quantified because the enriched natural product standard was not of sufficient quantity or purity to determine an accurate mass, but it was the most abundant DX by LC peak area of its most dominant peak. Parallel analysis of available KX standards (kauralexin B1 and A3) showed overall low abundance in WT and *Zmksl4* samples and no enrichment due to genotype. KA3 was found to be slightly enriched (1.7- old) in V5 *Zmksl4* roots compared to WT under well-watered conditions, but was not significantly enriched in V5 water deficit roots, V12 roots, or any leaf tissue (Supplemental Table 2B and 2D for root and leaf, respectively). The lack of key DX metabolites without significant variation in KX abundance in the *Zmksl4* mutant verifies that DX biosynthesis proceeds solely via ZmKSL4 without functional redundancies of known or yet uncharacterized maize diTPS.

### Expansion of the DX metabolite family

Next, LC-MS/MS untargeted metabolomics of WT and DX-deficient *Zmksl4* roots was used to investigate the chemical diversity of maize DX metabolites. To identify possible new DXs, a generalized linear model was used to identify features enriched in WT and comparatively depleted in *Zmksl4* roots (p < 0.1, fold change > 2.5) at developmental stage V12, given that this was the stage when THD and epoxydolabranol were found to be most abundant in WT (Fig. 2C). Given that there was no significant difference in epoxydolabranol or THD due to water status, samples were not separated by water status for this analysis. A total of 124 specific MS2 parent mass fragments at a particular retention time (RT), called “features”, were found to be enriched in WT but absent or substantially lower abundant in *Zmksl4* (Supplemental Table 2A), representing 0.3% of all detected features. Of these features, 91 had available MS2 spectra in at least one WT root sample at V12 (mass spectra and abundance plots of the most abundant feature for each predicted DX are in Supplemental Figs. 2 and 3, respectively).

**Figure 3.**
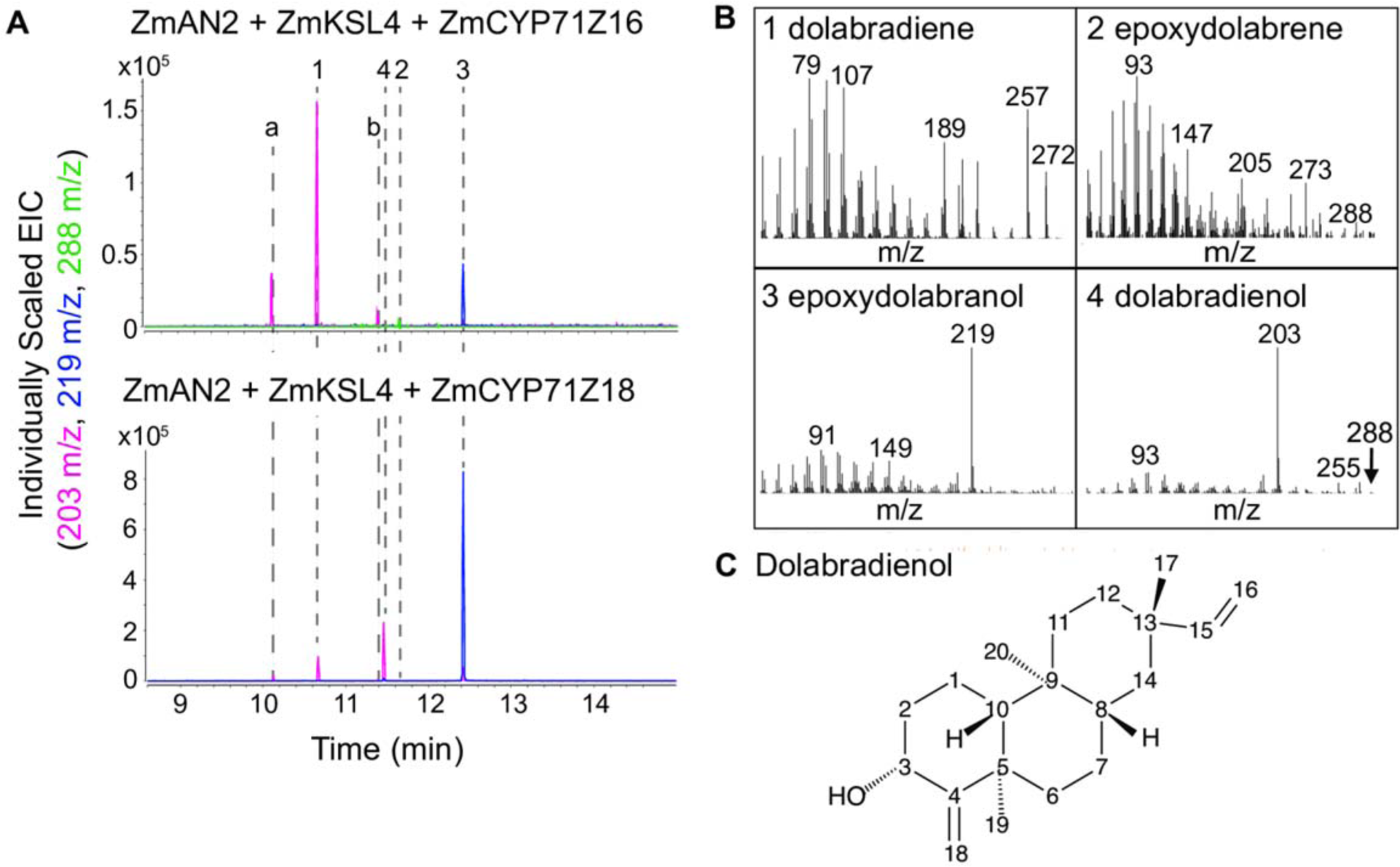
ZmCYP71Z18 makes a distinct dolabralexin (DX) product, dolabradienol. (A) Venn diagram of metabolite features enriched in WT roots compared to *Zmksl4*. Overlapping areas are features enriched at multiple developmental stages and water treatments. A core 34 features are known or predicted DX. (B) GC-MS extracted ion chromatograms of *E. coli* co-expression of maize diterpenoid biosynthesis genes. (C) Dolabradienol structure determined by NMR. (D) Mass spectra of ZmCYP71Z16/18 products, peak numbers correspond to peaks numbered in (B).

As expected, both THD and epoxydolabranol were found among the enriched features; the major THD feature represented the most abundant in this selected list, with a log2 fold change of 8.9 fold greater in WT than *Zmksl4*. As individual metabolites may be represented by multiple features, all focal features were manually inspected for RT and possible presence in the available standards, for a total of 48 predicted, uncharacterized DX metabolites. The list includes 32 metabolites in positive mode and 15 in negative mode. Several of these metabolites had peaks with the same parent ion and same MS2 at different RT (Supplemental Table 2A) – these represent either isomers or fragments of larger molecules that have the same structure. For features with parent masses less than 200 *m*/*z*, identical fragments are likely given that DX have the same backbone. Interestingly, DA-26 appeared to be an isomer of the parent mass of THD, not a matching fragment (Supplemental Fig. 2). The metabolites identified in positive mode range in their parent mass from *m*/*z* 95 to 391 and many contained characteristic diterpenoid mass ions such as *m*/*z* 81, 95, 119, and 145 that are also found in THD. All but eight of the positive mode, predicted DXs are within one minute of the THD retention time. While these metabolites were identified by their enrichment in all V12 WT roots compared to *Zmksl4*, 42 of the 48 predicted DXs were also enriched in V5 WT roots compared to *Zmksl4* under either just well-watered or water deficit conditions, or both, when samples were analyzed separately by water status.

To further investigate possible alterations in *Zmksl4* metabolite profiles beyond DX, statistical analysis using PERMANOVA was used to determine the factors influencing the root and leaf metabolomes. Genotype was not a significant factor in V12 root feature abundance, nor in leaves at any stage (statistics available in Supplemental Table 3A). Genotype was a significant factor for feature abundance in V5 roots, when DX are in low abundance in WT (Supplemental Table 3A). Outside of the 48 putative DX candidates, *Zmkls4* mutants did not cause major shifts in other metabolite groups of V12 maize roots, making *Zmksl4* plants suitable for investigating the diversity and biological relevance of DX metabolism in maize. No features were enriched in *Zmksl4* roots compared to WT at V12 when samples are not separated by water status (p < 0.10, fold change > 2.5); however, when separating by water status, some features were enriched, although not to the degree of DX enrichment in WT (Supplemental Table 2B). Mass spectra and RTs for all features are available online (see Methods for details), and lists of features enriched in each sample type, as well as their abundance and statistics related to significance, are available in Supplemental Table 2.

The complex co-occurrence of structurally related primary and specialized metabolites in maize tissue creates a challenge for DX purification for NMR structural elucidation. To circumvent this limitation, we used large-scale *E. coli* co-expression of ZmAn2, ZmKSL4, the maize cytochrome P450 reductase ZmCPR2, and ZmCYP71Z16 or ZmCYP71Z18 to produce previously unidentified pathway products to levels enabling purification (Fig. 3A and B). Of the resulting metabolites, one ZmCYP71Z18 product featured a RT and mass fragmentation pattern similar to, but distinct from, the known DX metabolites epoxydolabranol and epoxydolabrene (Fig. 3B and C). Purification of this compound using an optimized silica chromatography and semipreparative HPLC method (Murphy et al., 2019) enabled sufficient metabolite quantities to perform 1D and 2D NMR analysis, resulting in identification of the product as dolabradienol (Fig. 3D, Supplemental Fig. 4). Dolabradienol contains a single hydroxyl group in the C3 position and lacks the epoxy group of epoxydolabranol. In addition, comprehensive NMR analysis of DX products and select pathway intermediates refined previous structural assignments (Mafu et al., 2018), specifically defining a 5β, 10α rather than the previously reported 5α, 10β configuration of the DX skeleton (Fig. 1, Supplemental Fig. 4). Notably, co-expression of ZmCYP71Z16 did not result in the formation of detectable amounts of dolabradienol, suggesting catalytic differences between ZmCYP71Z16 and ZmCYP71Z18 (Fig. 3B). Dolabradienol and epoxydolabrene – the two minor products of ZmCYP71Z18 and Z16, respectively – represent opposite single oxygenation positions, but a combination of both oxygenations results in the epoxydolabranol structure seen as the dominant product for both ZmCYP71Z enzymes (Fig. 1; Fig. 3B). Additional LC-MS/MS analysis against the enzymatically produced dolabradienol standard revealed dolabradienol to be one of the 48 putative DX candidates, with 2.7-fold enrichment in WT V12 roots (Fig. 2C).

**Figure 4.**
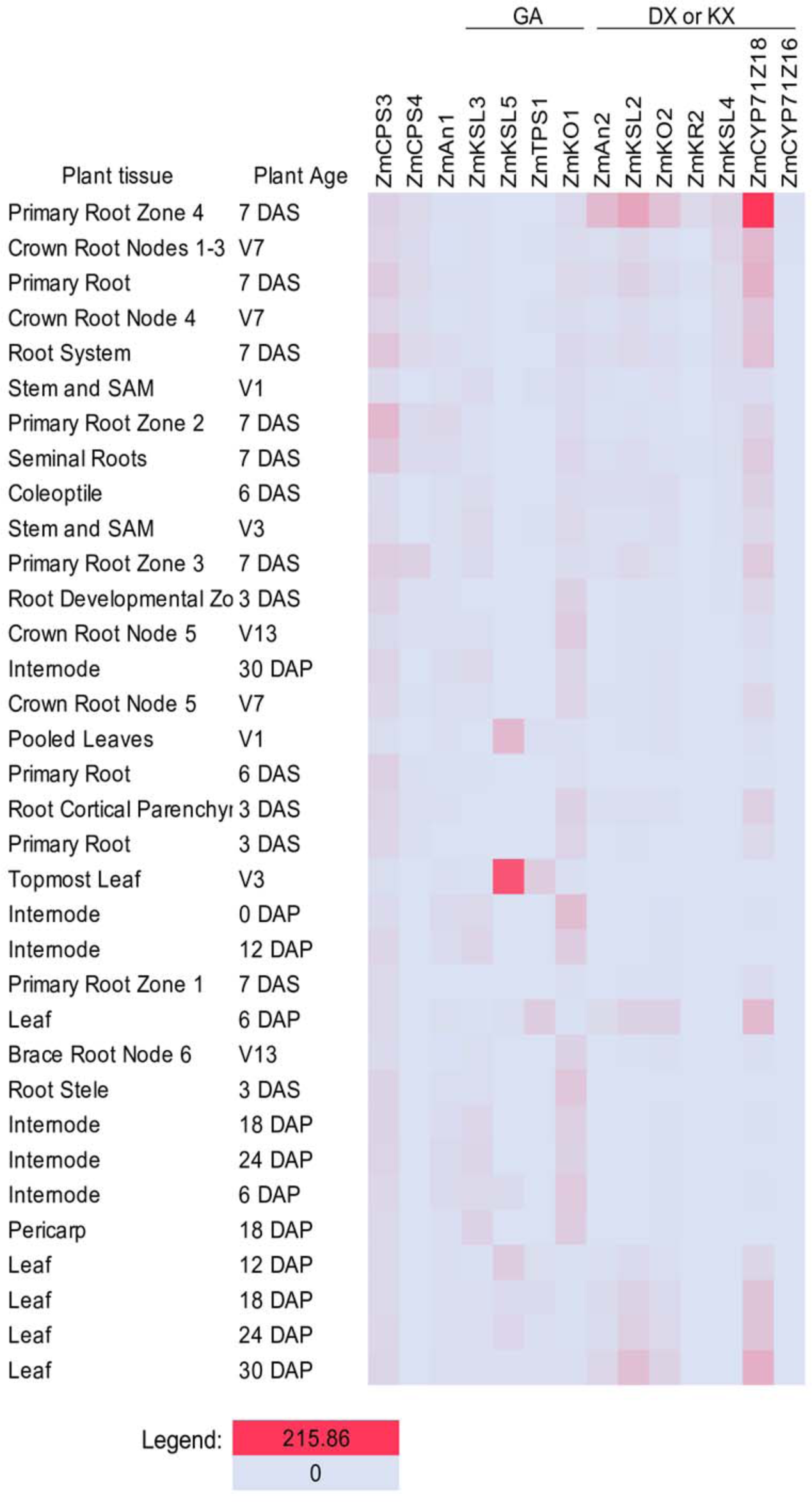
Spatiotemporal transcriptomes reveal tissue- and developmental-specificity of DX biosynthesis, as well as co-expression of KX and DX pathways. Data from Stepflug et al., sorted in descending order of *ZmKSL4* expression.

### Transcriptome analyses places DX biosynthesis predominantly in maize roots

GA, KX, and DX biosynthetic pathways all utilize the common precursor, *ent*-CPP, suggesting a tight control of the downstream pathway branches to drive the production of diterpenoids with distinct functions. To investigate the spatiotemporal differences in the biosynthesis of GA, KX, and DX metabolites, four publicly available RNAseq datasets (Chen et al., 2014; Stelpflug et al., 2016; Kremling et al., 2018; Yi et al., 2019) were analyzed for key genes of the respective GA, KX, and DX pathway branches (gene list in Supplemental Table 1). FPKM (fragments per kilobase of exon per million mapped fragments) values for the genes of interest from all datasets used here are available in Supplemental Table 4. Across all four developmental atlases (Chen et al., 2014; Stelpflug et al., 2016; Kremling et al., 2018; Yi et al., 2019), GA pathway genes, including *ZmAn1*, *ZmKo1*, *ZmKsl3* and to lesser extent *ZmKsl5* showed patterns of co-expression across various primarily above-ground tissues and timepoints (Bensen et al., 1995; Fu et al., 2016; Mao et al., 2017), as expected (Fig. 4). By contrast, *ZmAn2*, alongside *ZmKsl2*, *ZmKo2*, *ZmCYP71Z16*, *ZmKr2* and *ZmKsl4* of DX metabolism showed highest expression in roots (Fig. 4). Of the root tissues, *ZmKsl2* was most abundant in primary root tissues, whereas *ZmKsl4* expression was lower and more broadly distributed in different root tissue types. Notably, *ZmKsl4* showed highest expression in primary roots seven days postgermination (Fig. 4). Analysis of an additional, dissected root transcriptome dataset showed highest expression of *ZmAn2* and *ZmKsl4* in seminal and crown roots, with only low expression in primary roots (Tai et al., 2016), whereas another atlas (Kremling et al., 2018) showed highest *ZmKsl4* expression in root tips of germinating seedlings. Together, these transcriptome data support the predominant expression of the *ZmKsl4-*mediated DX pathway in select root tissues, whereas the *ZmKsl2*-derived KX pathway was expressed in both above- and below-ground tissues.

Given that *ZmKSL4* expression predominates in roots, we sought to confirm the developmental and tissue-specificity of DX metabolites. Using the LC-MS/MS metabolite data for WT maize plants generated in this study, we found DX were most abundant in roots at the V12 stage and were of low abundance in stage V5 (Fig. 2C). DX were not detectable or lowly abundant (less than 1 x 10^5^ peak height) in leaf tissue at all stages (Supplemental Fig. 3B), consistent with the gene expression patterns from the transcriptome tissue atlases. Previous work demonstrated stress-inducible transcript and metabolite accumulation in the KX pathway under drought stress (Vaughan et al., 2015), and both KX and DX under other stress conditions (Christensen et al., 2018; Mafu et al., 2018; Ding et al., 2019). Additional analysis of public datasets of above-ground tissue under heat and cold stress (Makarevitch et al., 2015) and *Cercospora zeina* leaf infection (Swart et al., 2017) showed no induction of *ZmKsl4*. In an analysis of a public dataset of four-day-old B73 seedlings treated with PEG8000 as a water deficit proxy, *Zman2*, *ZmKsl2*, *ZmCYP71Z16*, and to a lesser extent *ZmKsl4* were expressed in maize root cortex cells, but not the meristem, elongation zone, or stele, under both water deficit and well-watered conditions (Opitz et al., 2016). Here, our LC-MS/MS of WT roots showed characterized and newly predicted DX to be abundant in both water deficit and well-watered V5 and V12 roots (Fig. 2C), with no significant difference due to water status at either V5 or V12 (Supplemental Table 3A).

### Maize inbred lines vary significantly in their levels of DX pathway transcripts

To establish common patterns of DX abundance in maize roots, we next examined select genetically diverse maize lines. Prior studies demonstrated differential DX accumulation in roots of three field-grown maize varieties – B73, Mo17 and hybrid sweetcorn (variety Golden Queen) with B73 showing the lowest DX levels (Mafu et al., 2018). Using publicly available transcriptome data (Kremling et al., 2018), we assessed the transcript abundance in roots of germinated seedlings across a larger set of maize inbred lines; all public transcriptome data used in this study is available in Supplemental Table 4. Expression of the DX and KX pathway genes, including *ZmAn2*, *ZmKsl2*, and *ZmKsl4,* varied significantly across maize inbred lines, with B73 and Mo17 showing comparatively low transcript abundance for KX- and DX-metabolic genes, whereas high gene expression was observed for B79 and B75 (Kremling et al., 2018) (Fig. 5A). Gene expression also varied across multiple replicates of B73 included in this study, and were averaged across 17 replicates (Fig. 5A, Supplemental Table 4). Of the 274 lines tested, all but 3 had greater *ZmKsl2* expression than *ZmKsl4* in roots, and 8 had equal expression (Fig. 5A, Supplemental Table 4), consistent with the observed tissue-specific gene expression levels (Stelpflug et al., 2016; Kremling et al., 2018) (Fig. 4). Notably, DX and KX pathway genes showed patterns of co-expression with all major specialized diTPS genes in the network of most tested inbred lines (Kremling et al., 2018). Only in select lines, such as W22, *ZmKsl4* expression was not detected, whereas *ZmAn2* and *ZmKsl2* were expressed, suggesting that different lines may show distinct mixtures of the different metabolite groups (Kremling et al., 2018) (Fig. 5A).

**Figure 5.**
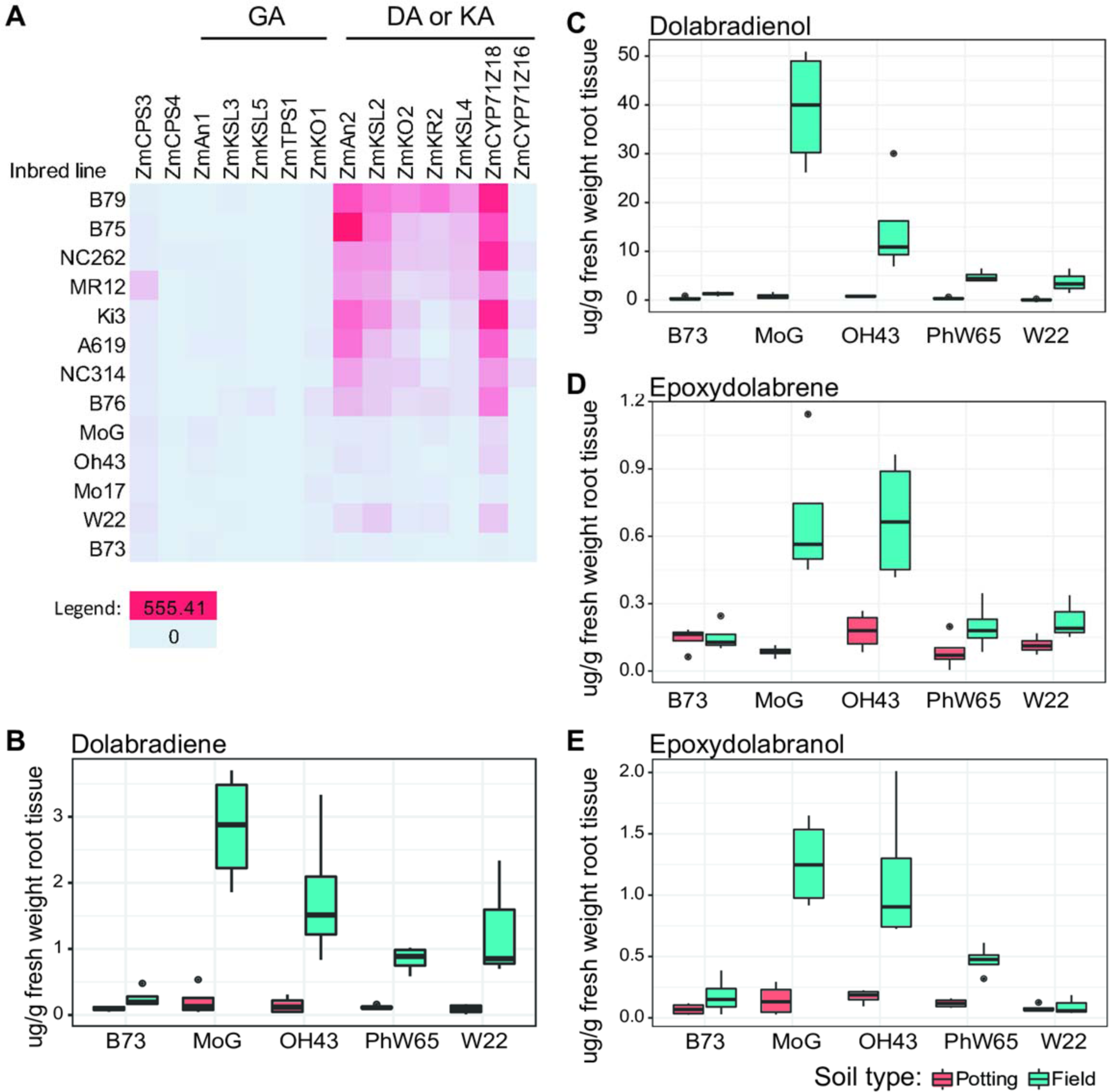
DX root transcript and metabolite abundances vary across maize inbred lines. (A) Transcript data is from Kremling et al. 2018. Data is sorted in descending order by *ZmKSL4* expression. B73 is averaged over 17 separate replicates. (B-E) Metabolite levels using GC-MS and authentic standards for major DX in a subset of available inbred roots grown in a greenhouse in field soil or potting soil.

To further investigate the influence of environmental and genetic variation, we grew four distinct maize inbred lines namely W22, PHW65, Oh43, and MoG, in the greenhouse using either commercial potting media comprised of peat moss or a 1:1 mixture of peat moss and soil from a maize field. In all cases root DX levels measured using GC-MS were nearly undetectable in peat moss, whereas DX levels in roots from plants grown in field soil were always significantly higher (Fig. 5B-E). Moreover, total DX levels varied across inbred lines by more than 7-fold, with MoG being the highest-producing line among those tested.

### *Zmksl4* shows reduced fitness under well-watered conditions

Given the stress-inducibility of DX and their abundance in roots under field conditions (Mafu et al., 2018), we performed physiological studies on *Zmksl4* mutant plants to understand the impact of DX-deficiency on *Zmksl4* vigor, under well-watered and water deficit conditions, as described above. To determine the effect of DX pathway products on the plant responses to water deficit, plants were analyzed for their fresh root and shoot weights at tissue harvest, using the same plants subsequently sampled for metabolite analysis described above. At V5, in which DX are only lowly abundant, genotype did not significantly impact root weight, shoot weight, or the root/shoot ratio, analyzed using Analysis of Variance (ANOVA), p < 0.05 (Fig. 6A). As expected, water stress significantly decreased root and shoot weights of both genotypes but did not affect the root/shoot ratio, as would have been expected (Fig. 6A). At V12, when DX were most abundant in WT samples, the root/shoot ratio significantly increased due to water deficiency in WT but did not increase in *Zmksl4*; *Zmksl4* showed 68% of the WT root/shoot ratio under water deficit conditions (Fig. 6B). Root and shoot weights were not impacted by either genotype or water status in V12 (Fig. 6B). Furthermore, while the WT root/shoot weight ratio average increased from 0.35 to 0.50 under water deficit conditions, the *Zmksl4* root/shoot ratio average ranged from 0.33 to 0.34 irrespective of the water condition (Fig. 6B).

**Figure 6.**
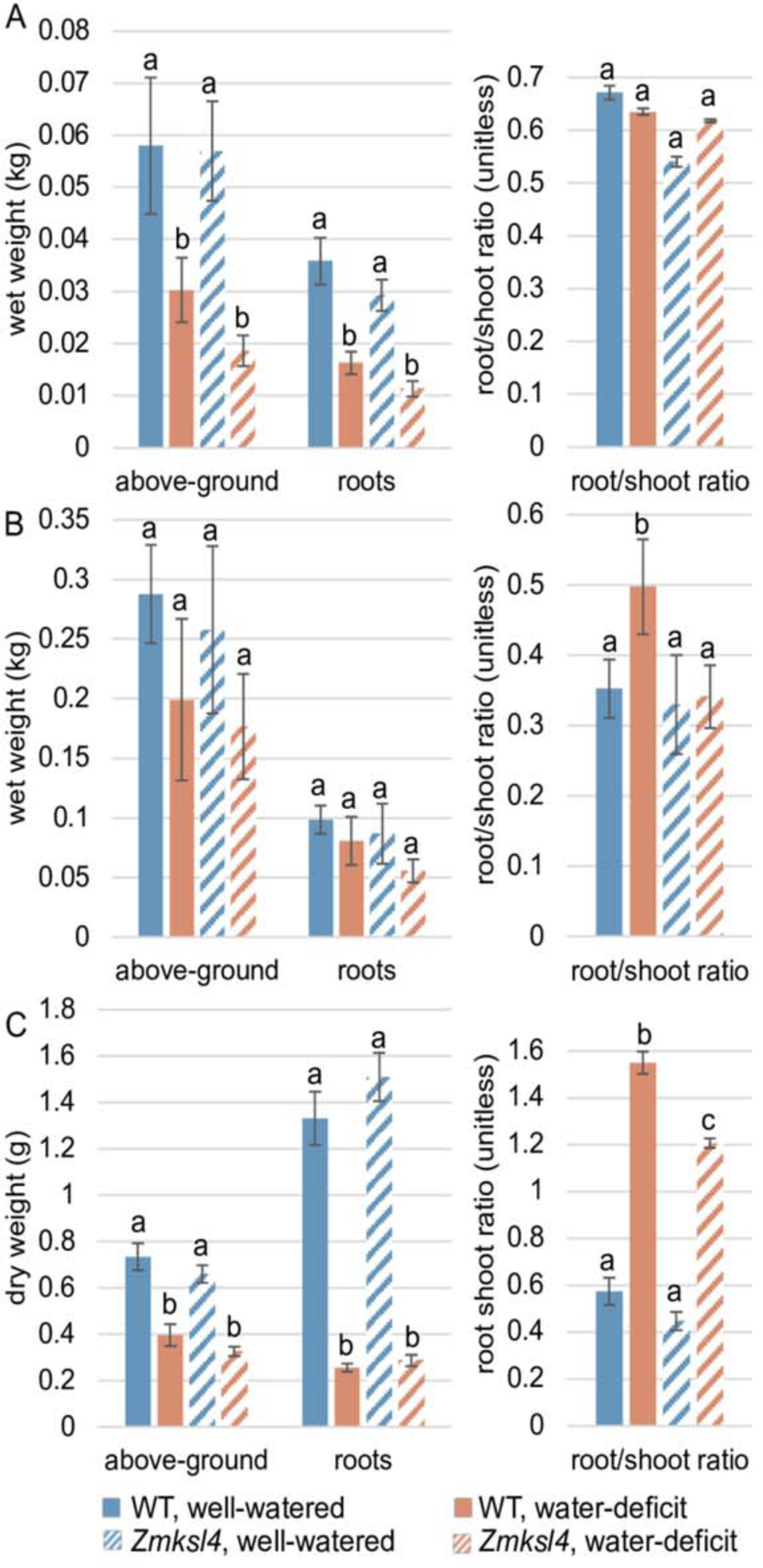
*Zmksl4* exhibits an altered root/shoot ratio. Letters represent statistical significance based on Tukey HSD t-test, with right-tailed values for water status and left-tailed values for genotype for root and shoot weight, and left-tailed values for both genotype and water status for root/shoot ratio. (A) Measurements based on fresh weight roots of all above-ground tissue of V5 roots used for metabolomics analysis. (B) Measurements based on fresh weight roots of all above-ground tissue of V12 roots used for metabolomics and transcriptomics analysis. (C) Measurements based on dry weight roots of all above-ground tissue of V5 roots used for root architecture analysis. Statistics and raw data available in Supplemental Table 3.

Given the altered *Zmksl4* response to drought stress, in that its root/shoot ratio did not change under water deficit at either stage (Fig. 6), we sought to understand broader differential *Zmksl4* response differences. To do so, LC-MS/MS untargeted metabolite profiles were used to comparatively analyze the metabolic response of WT and *Zmksl4* to water deficit. At V12, 3% of WT metabolomic features were significantly altered in abundance due to water deficit, as determined with a linear model (p < 0.1) (statistics in Supplemental Table 3A, metabolites and abundance in Supplemental Table 2C). By comparison, 8% percent of *Zmksl4* metabolites were changed. These features with altered abundance were dependent on genotype; only 9% of total features affected in all genotypes were commonly affected in both WT and *Zmksl4*, suggesting the genotypes have different metabolic responses to water deficit (Supplemental Fig. 5). Mass spectra and RTs for all features is available (see Methods for details), and features enriched in each sample type is available in Supplemental Table 2.

### *Zmksl4* has distinct root system architecture

To further investigate water deficit susceptibility of the *Zmksl4* mutant, the root system architecture of *Zmksl4* compared to its WT sibling was measured using root imaging and quantification of root traits using the DiRT software package. Here, WT and *Zmksl4* were grown in large pots of turface in a greenhouse until V5 (1 month after sowing), such that roots would not grow large enough to touch the insides of the pot. Turface was used, rather than field soil, because of its ability to easily wash away without damaging the root architecture. Water deficit was imposed by hydrating the media to a specific weight by volume prior to filling the pots; this level of hydration was then maintained by watering back to weight daily. Through this process the plants experienced a diurnal cycle of soil matrix potentials between approximately -0.7 MPa to -1.3 MPa, which was monitored hourly by matric potential sensors.

Measurements of dry tissue root and shoot weights recapitulated the previous experiment – genotype did not significantly impact root or shoot weight, but water had a significant impact (Fig. 6C). The difference between well-watered and water deficit weights for each genotype was greater than the previous experiment (Fig. 6A and C). Unlike the previous experiment at V5, the root/shoot ratio was significantly impacted by both genotype and water status (Fig. 6C). Agreeing with the V12 experiment, WT and *Zmksl4* plants have indistinguishable root/shoot ratios under well-watered conditions, but are significantly different under water deficit conditions (Fig. 6C). While both genotypes do show increased root/shoot ratios under water deficit, WT has a significantly greater increase than *Zmksl4* (Fig. 6C).

PCA analysis of root architecture traits showed separation of sample types by water status, but not by genotype (Fig. 7B). Separation was largely driven by root system surface area and width, leaf and root tip number, nodal root length and diameter, root density, and minimum lateral root branching angle, among others (Fig. 7C). Several specific root traits were affected under water deficit in both genotypes: Root system surface area, median and maximum root system width, and root tip number are among some of the most heavily affected traits under water deficit (Fig. 7D). While *Zmksl4* roots do have lower values for all these traits, it needs to be considered that *Zmksl4* plants are also smaller under optimal water conditions (Fig. 7D). Similarly, the total rooting depth remained unchanged by genotype, as well as the average lateral root length, and average lateral root diameter.

**Figure 7.**
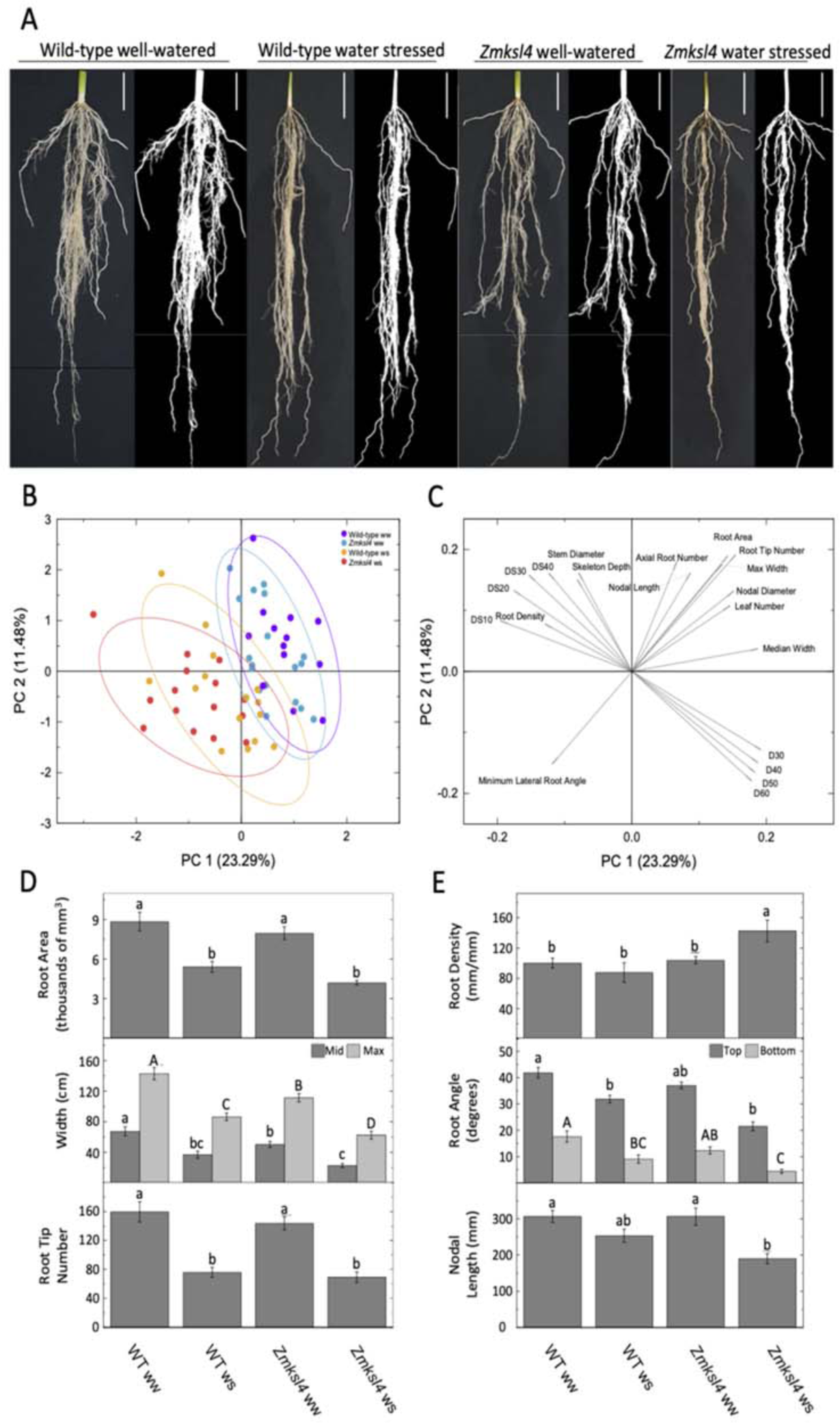
*Zmksl4* has altered root system architecture in V5 roots grown in turface. (A) Representative images, and their corresponding binary DIRT inputs, of wild-type and *Zmksl4* root systems under well-watered and water-stressed conditions, scale bars are 5 cm. (B) PCA analysis on DIRT traits show that groups separate based on watering treatment but largely overlap between genotypes. (C) The 20 of the most impactful DIRT trait vectors for PCA separation. (D and E) Specific root system traits impacted by genotype and water status; (D) many traits were affected to a similar degree relative to each well-watered control, (E) while others were affected differently in *Zmksl4* under water stress. Data are means ± SE (n=17-21). Different lettering represents significant differences at the 95% level determined by one-way ANOVA analysis. ww = well-watered, ws = water-stressed.

Specific traits were, however, significantly changed in the mutant (Fig. 7E). Under water deficit conditions, *Zmksl4* exhibited a significant increase in root system density compared to well-watered conditions, and compared to WT of either water status, up to 140% of well-watered levels. WT did not show increased root density upon water deficit. The root top angle, defining the angle of axial root trajectory from the horizontal soil line, was significantly decreased in *Zmksl4* plants, showing the roots grow more steeply downwards under both water conditions, resulting in a narrower root system. A similar observation was made for the root bottom angle, describing the angle of root branching from the horizontal soil line at depth. *Zmksl4* plants also showed a larger reduction in nodal root length in response to water deficit compared to WT.

To next investigate the root architecture phenotype in fully grown plants under field conditions, WT and *Zmksl4* plants were grown in the field until full maturity (i.e., seed harvest) and assessed by the DiRT software (Fig. 8A). Root crowns of 17 WT and 22 *Zmksl4* plants grown in Stanford, CA to full maturity were removed by uniform pulling of the plants until they were released from the soil. *Zmksl4* showed a severe reduction in root size (Fig. 8A) and analysis of root architecture traits significantly separated samples by genotype in PCA (Fig. 8B). Compared to WT, *Zmksl4* shows a reduced stem diameter and smaller root area, a lower width of the entire root system, both median and maximum, and a lower number of total roots (Fig. 8D). Additionally, *Zmksl4* showed a wider root diameter and a greater root density. Root top angle, however, was unaffected, and because these were crowns and not complete root systems, the root bottom angle could not accurately be quantified. The Drop 50, the measurement of the depth where 50 percent of the root tips have emerged, was larger for WT, depicting WT with deeper roots than *Zmksl4* (Fig. 8A and D).

**Figure 8.**
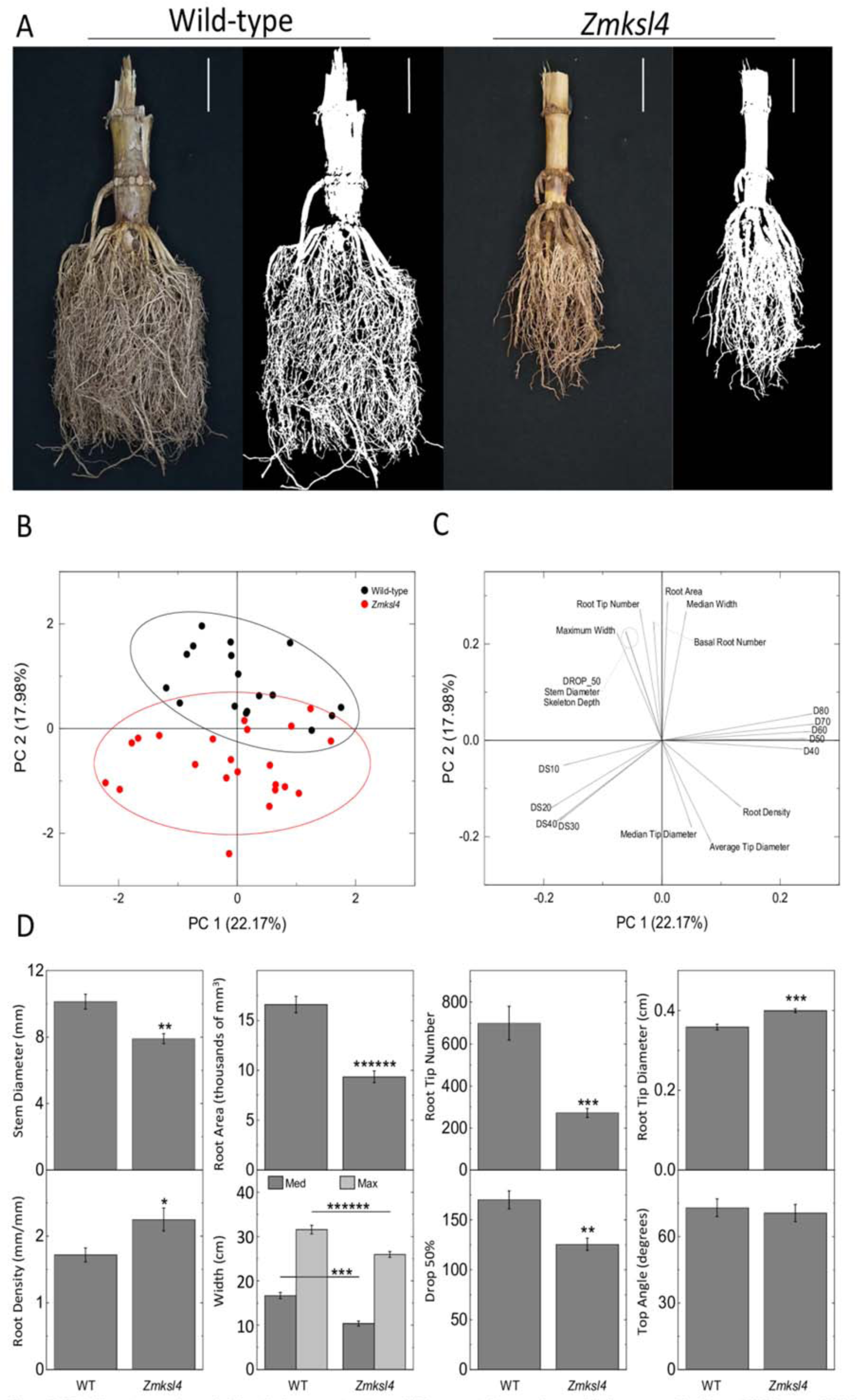
*Zmksl4* root crowns have significantly denser root systems. (A) Representative samples, and their corresponding binary DIRT inputs, of wild-type and *Zmksl4* mature root crowns, scale bars are 5 cm. (B) PCA analysis on DIRT traits show clear separation of genotypes. (C) The 20 of the most impactful DIRT trait vectors for PCA separation. (D) Several important root architectural traits were shown to be different between wild-type and *Zmksl4.* Data are averages ± SE (n = 17-21). Levels of significance were determined by student’s t-test; * = p ≤ .01, ** = p ≤ .001, *** = p ≤ .0001, ****** = p ≤ .0000001.

### *Zmksl4* transcriptome shows *Zmksl4* responds differently to water deficit

Transcriptome analysis of the *Zmksl4* mutant plants was performed using V12 roots, previously assayed for metabolite content and root/shoot weights (Fig. 2B), with a focus on the developmental stage with highest DX abundance to investigate connections between DX production and plant stress responses. Principal component analysis (PCA) demonstrated that while water deficit accounted for 73% of sample variance, genotype had a minimal effect of 6% (Supplemental Fig. 7A), separating the genotype sample clusters under water deficit conditions but not under well-watered conditions (Supplemental Fig. 7A). Furthermore, in a heat map of sample-to-sample distances, sample types predominantly cluster by water status, with limited clustering due to genotype (Supplemental Fig. 7B). Consistent with the genotype and environment effects on untargeted metabolite profiling described above, the mutation in *Zmksl4* had less of an effect on gene expression than did water status, and there are a small but significant number of DEGs affected by the genotype by water status interaction (Supplemental Table 3E).

Next, differentially expressed genes (DEGs) were determined using the DESeq2 package in R (Supplemental Table 5). Among the analyzed diTPS and P450 genes, only *ZmKsl4* was differentially expressed, as expected in the mutant (Fig. 9A, Supplemental Table 5A and B). However, one known diterpenoid pathway gene, *ZmKr2*, was differentially expressed between WT and *Zmksl4* under both well-watered and water deficit conditions. Of the currently known nine genes specifically impacting maize root architecture, *RTH1* (*ROOTHAIRLESS 1*)*, RTH3, RTH, RTH5, RTH6, RUM1* (*ROOTLESS WITH UNDETECTABLE MERISTEM 1*), *RTCS* (*ROOTLESS CONCERNING SEMINAL AND CROWN ROOTS*), *RTCL* (*RTCS-LIKE*), *RUL1* (*RUM1-LIKE*), and *LRP1* (*LATERAL ROOT PRIMORDIA 1*) (Bray and Topp, 2018), none showed differential expression between genotypes under water deficit or well-watered conditions. However, the gene *Zm00001d033169*, which is orthologous to the *Arabidopsis* gene *MORPHOGENESIS OF ROOT HAIR 6* (*MRH6*), was significantly down-regulated in *Zmksl4*. Of the 45,808 genes with non-zero expression, 136 genes were differentially expressed between *Zmksl4* and WT regardless of water status (Fig. 9B, Supplemental Table 5C). Of these DEGs due to the mutation (Supplemental Table 5A-C), there was no significant enrichment for any Gene Ontology (GO) terms. Genes down-regulated in *Zmksl4* roots under both stress conditions included transcription factors *C2C2-GATA-TRANSCRIPTION FACTOR 15* (*GATA15*), *OPAQUE2 HETERODIMERIZING PROTEIN1* (*OHP1*), AND *GNAT-TRANSCRIPTION FACTOR 7* (*HAGTF7*). *INDOLE-3-GLYCEROL PHOSPHATE LYASE1* (*IGL1*), a gene involved in benzoxazinoid biosynthesis, was also down-regulated, along with *NITRATE REDUCTASE 4* (*NNR4*), *GRANULE-BOUND STARCH SYNTHASE-I* which is more commonly known as *WAXY1* (*WX1*), and *PURPLE ACID PHOSPHATASE16* (*PAP16*) that is involved in plant phosphorous processes. Transcript levels of *Zm00001d003716,* orthologous to the *Arabidopsis* gene *USUALLY MULTIPLE ACIDS MOVE IN AND OUT TRANSPORTERS 25* (AT1G09380, UMAMIT25) were also strongly suppressed in *Zmksl4* roots, suggesting broader alteration of transport processes.

**Figure 9.**
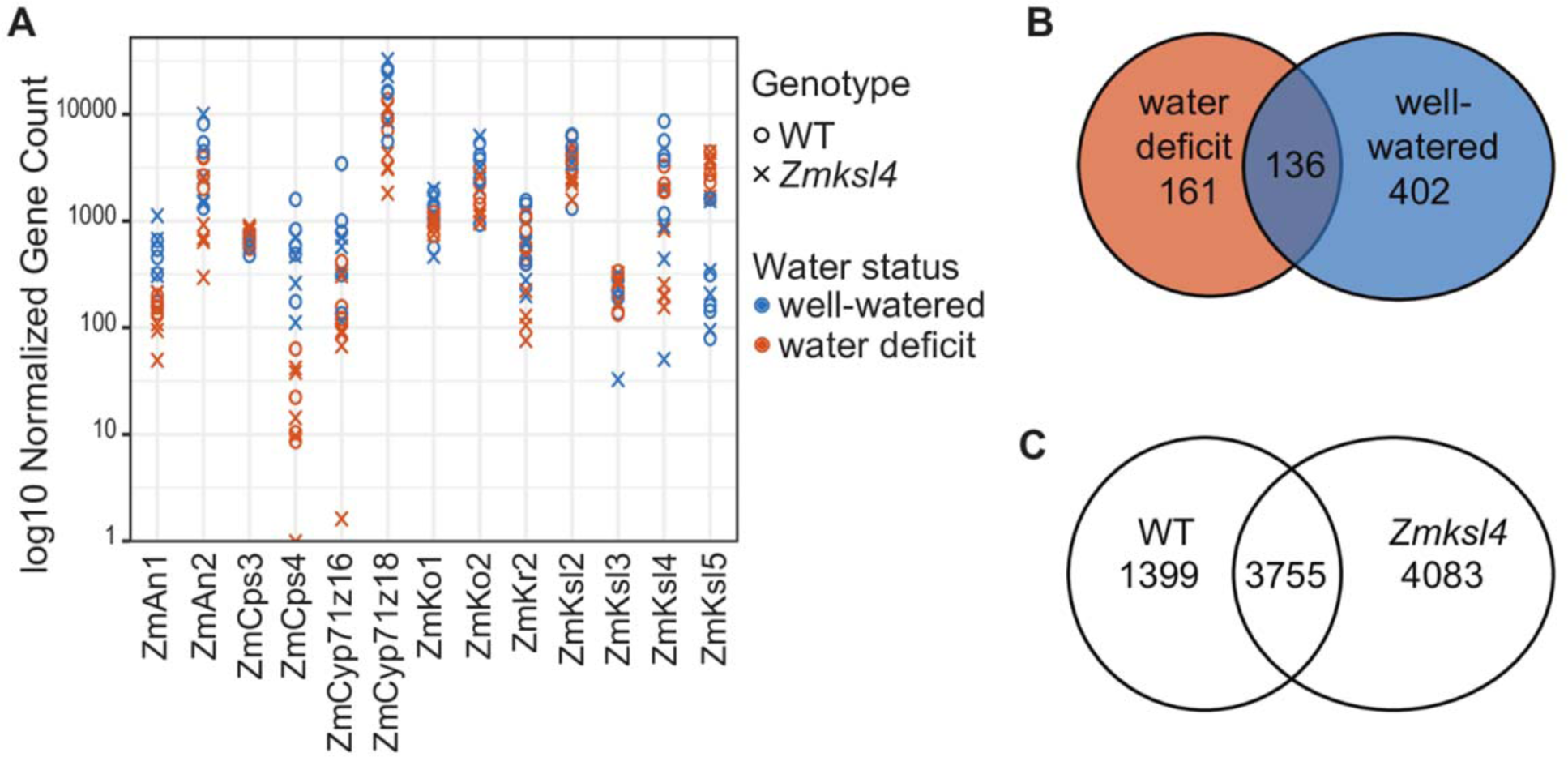
Transcriptome of *Zmksl4.* (A) Plot of normalized gene counts for diterpenoid pathways; only *ZmKsl4* is a differentially expressed gene (DEG). (B) Number of DEGs impacted by mutation for each water status; overlap are genes differentially expressed regardless of water status. (C) Number of genes differentially expressed due to water status for each genotype, respectively; overlap are genes differentially expressed due to water status in both genotypes. Genes in (B) and (C) are listed, along with fold change values, in Supplemental Table 5.

As demonstrated for significantly enriched metabolites, water status impacts each individual genotype differently: the number of genes commonly affected by water status in both genotypes was low, suggesting unique responses, and the number of genes affected in *Zmksl4* was greater (Fig. 9C, Supplemental Table 5E-G). Furthermore, GO terms enriched in the DEGs by water status in each genotype, independently, were largely different, supporting the different responses to water stress in the mutant (Supplemental Table 6A-D).

## Discussion

Diterpenoids have important functions in species-specific plant defense and environmental adaptation. In maize, KX and DX metabolites form two pathway branches with demonstrated or predicted roles in biotic and abiotic stress responses, as evidenced in previous work by the susceptibility of the *Zman2* and *Zmksl2* mutants to different stressors, the impact of KX and DX deficiency on the rhizosphere microbiome, the stress-inducibility of pathway transcripts and metabolites, and the *in vitro* anti-fungal activity of select metabolites (Schmelz et al., 2011; Christensen et al., 2018; Mafu et al., 2018; Ding et al., 2019; Murphy et al., 2021) (Fig. 1). In the current effort, we created a *Zmksl4* knock-out mutant with selective DX-deficiency, enabling us to isolate the roles of DX from interconnections with ZmAn2 and KX metabolism. Lack of DX metabolites in *Zmksl4* demonstrates that ZmKSL4 is the sole enzyme responsible for DX biosynthesis, and that without other significant impacts to the global metabolome or transcriptome, the mutant is suitable for analyzing the stress-protective relevance of DXs in maize (Fig. 2).

Comparative metabolite profiling of the *Zmksl4* mutant and its WT sibling demonstrate that DX encompass a larger metabolite family than previously identified. Here we predict up to 48 additional metabolites (Supplemental Table 2A, Supplemental Fig. 2 and 3) belonging to the DX family based on their absence in *Zmksl4* roots and their abundance in WT roots at V12. Because multiple features may belong to the same metabolite, or multiple metabolites may co-elute at the same RT, this number represents an estimate to guide further study. Many of these enriched features had the same parent mass and similar MS2 spectra, but different RTs, suggesting they are either isomers or that some of the fragment ions are identical, which is expected of metabolites in a pathway based on the same hydrocarbon scaffold.

NMR structural analysis of additional products of the combined activity of ZmAn2, ZmKLS4 and the P450 ZmCYP71Z18 (Supplemental Fig. 4) identified one of these metabolites as the dolabradiene-derivative, dolabradienol, that we subsequently confirmed as highly abundant in roots of maize grown with field soil (Fig. 2C, Fig. 3). Absence of dolabradienol in enzyme assays using the closely related ZmCYP71Z16 support catalytic differences between these otherwise highly promiscuous P450s, both of which generate epoxydolabranol as the dominant product. The broader range of DX metabolites in mature maize roots is well aligned with the largely root-specific expression of relevant DX pathway genes as shown here and in prior studies (Mafu et al., 2018). This further supports a below-ground function of DX metabolism, contrasting the more tissue-wide distribution of the KX pathway despite their shared use of ZmAN2 (Schmelz et al., 2011; Christensen et al., 2018; Mafu et al., 2018; Ding et al., 2019).

Our observation that DX metabolites are produced in mature roots of several maize inbred lines grown in field soil, yet significantly less abundant in plants grown in peat moss, suggests the presence of field soil factors play a role in DX production, which we hypothesize to include microorganisms (Fig. 5). While select DX metabolites have been shown to be antibiotic *in vitro*, the number of additional potential roles in complex biotic and environmental interaction in the soil are myriad (Mafu et al., 2018; Ding et al., 2021; Murphy et al., 2021). Prior analyses of *Zman2* mutants, now appreciated to be deficient in both KX and DX, demonstrated maize root diterpenoids collectively influence the rhizosphere microbiome, and support a collective role of root DX metabolites in plant-microbe interactions (Murphy et al., 2021). In contrast to earlier studies with young maize plants and root KX (Vaughan et al., 2015), our current study demonstrates that unlike KX, water deficit does not induce the accumulation of DX metabolites, but rather these metabolites are highly abundant in older plants grown in field soils under both well-watered and water deficit conditions tested here (Fig. 2C, Fig. 5). Maize DX display a high degree of root specificity and contrast KX metabolites, which are found both above and below ground at the local site of pathogen or pest attack in leaves, stems, and roots (Schmelz et al., 2011; Vaughan et al., 2015; Ding et al., 2019). Transcript and metabolite profiling of a range of maize inbred lines showed substantial diversity of DX formation across maize inbred lines (Fig. 5). For example, MoG roots produce >20-fold more DX than B73 roots at the same stage when grown in a greenhouse using field soils (Fig. 5). Select maize inbreds appear to produce and rely on different diterpenoid blends under different environmental conditions which provides an opportunity to investigate how distinct diterpenoids contribute to stress resilience and ultimately leverage such insights for understanding chemical defense traits and potential roles in breeding.

Although no increase in DX levels was detected in response to water deficit, phenotypic analysis of the DX-deficient *Zmksl4* mutant revealed a reduced root/shoot ratio and a reduced responsiveness to water deficit as compared to the WT sibling in both V5 and V12 roots, depending on the experiment (Fig. 6). Previous research at the seedling stage showed the *Zman2* mutant to have a reduced root/shoot ratio under drought stress, and increased KX levels (DX were unknown at the time of the study and not detected) (Vaughan et al., 2015). In our current effort, *Zmksl4* demonstrated a genotype by environment (GxE) interaction, with a different response to water deficit than WT in regard to its root/shoot ratio and changes in the metabolome and transcriptome. For example, the metabolites and transcripts that were differentially present under water deficit were largely different in *Zmksl4* and WT, and *Zmksl4* had a greater number of metabolites and unique transcripts changed (Supplemental Tables 2 and 5). These changes could be causing the differences in root architecture or could be a downstream effect of the altered architecture.

Given the root predominance of DX metabolism, and the altered root/shoot ratio, analysis of the root architecture of WT and *Zmksl4* plants in water-deficit and well-watered conditions was used to investigate how DX metabolites may impact plant vigor in *Zmksl4* mutant plants. Even under optimal conditions, *Zmksl4* in the early vegetative and fully mature stages demonstrated stunted growth in nearly all examined root traits, and mature root crowns of *Zmksl4* were significantly denser than WT (Fig. 7 and 8). While both WT and *Zmksl4* showed many classical water deficit responses, such as a reduction in stem diameter, root system surface area, and the total number of roots, only *Zmksl4* showed distinct stress symptoms, including most notably an up to 140% increase in root system density (Fig. 7 and 8). This response was not seen in WT roots, and is likely a result of the compounding effects of adopting a sharper rooting angle under the imposed stress while maintaining a similar level of root number to WT. However, it is important to note that many other root traits were affected in similar ways between genotypes when accounting for the change in the trait value between well-watered and water-deficit rather than absolute difference between WT and *Zmksl4* under stressed conditions. This suggests many of the differences observed in root architecture are due to overall environmental effects which may be more important than differential genotypic responses to water deficit specifically. The mutant phenotype suggests DX play a role in root growth and biomass, which in turn impacts plant vigor with regards to the root/shoot ratio under water deficit conditions (Fig. 6).

We hypothesize that observed differences in root architecture could result from altered layers of biochemical immunity, gene expression levels, and/or changes in the rhizosphere microbiome assembly, which includes plant-pathogen interactions. A modest level of significant changes in the *Zmksl4* transcriptome compared to WT, 136 genes independent of water status (Fig. 9), speak against large-scale changes in the regulation of root development. However particular DEGs could be directly impacting the root architecture, such as the transcription factors *GATA15*, *OHP1*, and *HAGTF7*, all of which were significantly down-regulated in *Zmksl4* in both water conditions. Another candidate of interest in the *Zmksl4* suppression of *Zm00001d033169*, orthologous to Arabidopsis MRH6, which has been implicated in root hair development. Changes in plant nutrient acquisition, processing, and transport could also be a cause or effect of the altered root architecture; both *NNR4* and *PAP16* were down-regulated and play roles in nitrate and phosphorous nutrition, respectively. Displaying opposite transporter responses, *Zmksl4* mutants demonstrated a suppression of *Zm00001d003716* (*Arabidopsis* ortholog, UMAMIT25) transcripts and increased accumulation of *Zm00001d019131* encoding an ATP-binding cassette (ABC) protein (Supplemental Table 5A and B). Given that most genes in the dataset remain unannotated, however, the precise causal effects of root architecture changes in *Zmksl4* remain to be investigated in future work. Complex interactions such as compositional microbiome changes in the DX- and KX-deficient *Zman2* mutant (Murphy et al., 2021) are supportive of DX roles in microbiome interactions which could further impact root architecture as an important component of water stress resilience, as plants navigate the soil for optimum water absorption. Whether changes in DX production impact maize gene expression directly, or indirectly via impacts on the rhizosphere microbiome, and whether the root architecture is a cause or product of either changes, remains to be investigated in detail.

The mechanisms underlying DX-mediated changes in root development and overall plant vigor can be further understood following identification of the larger and complex network of terminal DX pathway products made by roots and void from *Zmkls4* mutants (Supplemental

Table 2A). Future robust functional studies require defined mutants and sufficient purified pathway products. DX metabolites exist as significant metabolites in the roots of mature maize plants grown in field soils yet are uncommonly observed in standard greenhouse conditions. To date, pathogen induced regulation of the DX pathway only partially matches established patterns for the KX pathway activation (Mafu et al., 2018; Ding et al., 2019) which collectively implies biological roles for DX beyond antifungal defenses. Our collective findings create a foundation to elucidate the tissue- and pathway-specific functions of DXs that ultimately could inform for stress resilience breeding and engineering strategies in maize.

## Methods

### Generation of a maize *Zmksl4* loss-of-function mutant

*Zmksl4* mutant plants and isogenic WT sibling plants were generated as previously described (Char et al., 2016). In short, guide RNA uniquely targeting two genes, *ZmKsl4* and *ZmCYP71Z16*, was inserted into the pGW plasmid (based on the pMCG1005 backbone) containing the rice codon-optimized *Streptococcus pyogenes Cas9* under control of the maize *ubiquitin 1* promoter, and the *bar* gene under control of the 4xCaMV35S promoter as a selectable marker. The resulting construct was transformed into the Hi-II maize hybrid line using *Agrobacterium tumefaciens,* strain EHA101(Char et al., 2016). To generate the gene-specific *ZmKsl4* mutant, transgenic plants were then outcrossed to B73 and genotyped to select for plants that lacked the *Cas9* gene, were heterozygous for a one base pair insertion in *ZmKsl4*, and contained no mutation in *ZmCYP71Z16*. Genotyping was performed using PCR for *Cas9*, *ZmKsl4*, and *ZmCYP71Z16* using gene-specific primers and Phusion polymerase (New England BioLabs) according to manufacturer instructions. Presence or absence of *Cas9* was used for genotyping. For *ZmKsl4* and *ZmCYP71Z16*, PCR products were analyzed using PCR clean-up and sanger sequencing for the region of interest. An insertion of G or A at position 354 bp in the first exon of *ZmKsl4* was verified, resulting in a mutant featuring two different alleles that cause a frameshift rendering the protein truncated and predictively non-functional. Two plants, each without *Cas9* and heterozygous for a mutation, were crossed and genotyped, generating a 1:2:1 segregating population for homozygous mutant:heterozygous:homozygous WT (selfing was not possible because of developmental delays between ears and tassels). Two homozygous mutant plants were crossed (again due to developmental delays) to generate seed used in this study. One WT plant was selfed for use as a WT sibling. LC-MS/MS of the mutant and WT sibling described in this study confirm the absence of DX in *Zmksl4* and presence in its WT sibling.

### Plant growth and materials for drought elicitation

Maize plants were grown at UC Davis field site (Davis, CA, USA). To control water deficit conditions, field soil used for maize cultivation the prior year was sieved and placed into 3-gallon fabric pots on the field site. Seeds were sterilized by incubation with 15% bleach for 15 min, and rinsed thrice with sterile water. One seed per pot was planted by hand. Plants were irrigated using controllable irrigation drip tape. For timepoint 1, plants were grown for 30 days before half of the plants were subjected to 6 days of water deficit, consisting of half the water provided via irrigation. At the end of timepoint 1, well-watered plants were at the V5 stage. For timepoint 2, plants were grown for 61 days before half of the plants were subjected to 6 days of severe water deficit, in which they were provided half of the water via irrigation. Water deficit conditions for both timepoints was confirmed by a characteristic leaf rolling phenotype. Well-watered plants were at the V12 stage for timepoint 2. Plants were fertilized twice during the growing season using solubilized ammonium phosphate, potassium nitrate, and ammonium sulfate through the irrigation system so that each plant received, in total, 14.5 g of NaCl, 2.5 g of N, 0.5 g P_2_O_5_, and 2.5 g of K_2_O.

On the day of harvest (day 36 for V5; day 67 for V12), plants were removed from their pots and roots washed gently with water to minimize root damage and photographed. Plants were cut at the root-shoot junction and weighed for plant fresh weights. Washed roots and the uppermost leaf with a leaf collar were collected separately, frozen immediately in liquid N_2_, and stored at -80°C for downstream analysis. For each sample collection, the biological replicates were as follows: V5: n = 4 (WT, well-watered), n = 7 (WT, water deficit), n = 7 (*Zmksl4*, well-watered), n = 6 (*Zmksl4*, water deficit). V12: n = 5 (WT, well-watered), n = 4 (WT, water deficit), n = 5 (*Zmksl4*, well-watered), n = 6 (*Zmksl4*, water deficit).

To consider the interactions of maize genotypes and soil types on resulting DX production, all maize inbreds (PHW65, Oh43, MoG, B73, W22) were grown in the greenhouses at the UCSD Biology Field Station for 40 days. Plants were grown in either commercial peat moss based potting soil (BM2; Berger Corp) or a 1:1 mixture of local maize field soil and potting soil. Potted plants were removed from the soil, root systems were washed with H_2_O, frozen with liquid N_2_, ground in liquid N_2_ using a mortar and pestle to a fine powder and stored at -80°C. Working with liquid N_2_ and -80°C is essential to halt all enzyme activities and maintain stability of potentially labile analytes.

### P450 functional analysis

Functional characterization of ZmCYP71Z18 was performed using *E. coli* co-expression assays as previously described (Murphy et al., 2019). In brief, the following plasmids were transformed into BL21DE3-C41 cells (Lucigen): codon optimized pET- DUET1:ZmCPR2/ZmCYP71Z18 or ZmCYP71Z16; pET28b:ZmKSL4Δ106; pCOLA-DUET:GGPP-synthase:ZmAN2; pIRS plasmid of upstream pathway as described previously (Morrone et al., 2010; Mafu et al., 2018). Transformed *E. coli* was grown in 50 mL of Terrific Broth at 37°C until an OD_600_ of ∼0.6 was reached. Cultures were cooled to 16°C and protein production was induced with 1 mM isopropyl-thiogalactoside and supplemented with 25 mM sodium pyruvate, 4 mg/L riboflavin, and 75 mg/L δ-aminolevulinic acid. Enzyme products were extracted with 50 mL of 1:1 hexanes:ethyl acetate and a separatory funnel, and concentrated under N_2_ stream. Enzyme products were analyzed using GC-MS and NMR.

### Root metabolite extraction

Collected leaf and root samples from the Davis field site were frozen in liquid N_2_ and homogenized twice in 30-sec increments with freezing in liquid N_2_ in between using a tissue homogenizer (Retsch). The frozen, pulverized tissue was weighed, and methanol with 0.01% formic acid was added at 10:1 volume:tissue weight for metabolite extraction. Samples were gently shaken at room temperature for 24 hrs, after which 2 mL of extract were removed to a glass vial and stored at -80°C. A total of 200 µL of extract was then filtered into a 96-well plate using a filter plate to remove any possible particulates. V5: n = 4 (WT, well-watered), n = 7 (WT, water deficit), n = 7 (*Zmksl4*, well-watered), n = 6 (*Zmksl4*, water deficit). V12: n = 5 (WT, well-watered), n = 4 (WT, water deficit), n = 5 (*Zmksl4*, well-watered), n = 6 (*Zmksl4*, water deficit).

### GC-MS analysis

GC-MS analysis of *E. coli* generated metabolites was performed on an Agilent 7890B gas chromatograph with a 5977 Extractor XL MS Detector at 70 eV and 1.2 mL/min He flow, using an Agilent HP5-MS column (30 m, 250 mm i.d., 0.25 mm film) as previously described (Mafu et al., 2018; Murphy et al., 2019). Samples were suspended in 1 mL of hexanes, with 1 µL injection volume, pulsed splitless injection at 250°C inlet temperature. Oven temperature was ramped from 50°C to 300°C at 20°C/min and held for 3 min. MS data from 90 to 600 *m*/*z* were collected after a 10-min solvent delay.

For analysis of maize plants grown in different soil types, a simple approach to sample analysis relied on Vapor Phase Extraction (VPE) to remove high molecular weight analytes otherwise un-compatible with gas chromatography (GC). In this modified procedure, finely ground 50 mg sample aliquots were weighed, extracted with 1ml 1-propanol:hexane (3:10) and the resulting organic phase was derivatized using trimethylsilyl diazomethane (Schmelz *et al*., 2004). Following 20 min for derivatization, all liquids were dried under a N_2_ stream with care taken to avoid over-drying which could result in a loss of diterpenoid analytes. The modified sample preparation avoids the use of plasticware at all steps prior to VPE thus minimizing all sources of plasticizer contamination. Final analytical samples were eluted from the VPE traps using 150 ml of 1:1 hexane:ethyl acetate. GC/MS analyses were conducted using an Agilent 6890 series gas chromatograph joined to an Agilent 5973 mass selective detector (mass temperature, 150°C; interface temperature, 250°C; electron energy, 70 eV; source temperature, 230°C). DB-35 MS column (Agilent; 30 m × 250 μm × 0.25 μm film) was used for gas chromatography. Samples were introduced with an initial oven temperature of 45°C, as a splitless injection. The temperature was held for 2.25 min, then increased to 300°C with a gradient of 20°C min^−1^ and held at 300°C for 5 min. A solvent delay of 4.5 min was selected to prevent ethyl acetate present in the sample from damaging the EI-filament. GC–MS-based quantification of DX utilized an external standard curve of dolabradienol that was spiked into similar root tissue samples lacking DX metabolites. Product identification of previously known analytes was conducted using authentic standards. Agilent Mass Hunter Qualitative and MS Quantitative Analysis software alongside Agilent ChemStation qualitative programs were used to generate and analyze the GC-MS generated chromatograms and spectra. Replicated experiments were summarized with peak areas captured in MassHunter Qualitative Navigator B.08.00, and MS Quantitative Analysis B.08.00, quantified in Excel, and statistically evaluated in JMP. MassHunter MS Quantitative program peak selection methods were manually produced taking into account RT shifts. Program automated peak selections/integrations were instead substituted for manual selections/integrations of every target compound in every sample.

For purifying dolabradienol (2R,4bS,7S,10aR)-4b,7,10a-trimethyl-1-methylene-7- vinyltetradecahydrophenanthren-2-ol for use as a standard to quantify metabolite levels in testing maize plants in different soil types, 300 grams of 35-day-post-pollination field-grown MoG root tissue was ground to a fine powder in liquid N_2_, extracted first with 500 mL of methanol then secondarily extracted with 500 mL of ethyl acetate (EA), filtered and dried using a rotary evaporator. The resulting oily residue was then allowed to partition and re-dissolve in 300 ml EA. The EA fraction was then dried down and the resulting oily residue was then separated by preparative flash chromatography (CombiFlashRf; Teledyne ISCO) on a 5g C18, reverse phase, (RediSepRf High Performance GOLD) column. The mobile phase consisted of solvent A (100% H_2_O) and solvent B (100% Acetonitrile), with a continuous gradient of 0-100% B from 1 to 81 min using a flow rate of 18 ml min^−1^. 100 x l aliquots of these fractions were then derivatized using trimethylsilyldiazomethane, dried, then resolubilized in 200 ml of 1:1 hexane:EA and analyzed by GC/MS. One fraction spanning 42-43 minutes contained an enrichment of dolabradienol. Select concentrated samples were further purified by high performance liquid chromatography HPLC using repeated methylated/derivatized (using trimethylsilyldiazomethane) and 1 mL injections (passed through silica to filter out precipitate following derivatization) on a Zorbax RX-silica (250 x 4.6 mm, 5 µm; Agilent) column and a mobile phase consisting of solvent A (100% hexanes) and solvent B (100% EA) with a continuous gradient of A–B from 2 to 37 min using a flow rate of 1 ml min^−1^. The recollected HPLC fractions spanning 11-12 min retention times (RT) yielded dolabradienol at ∼95% purity.

### LC-MS/MS analysis

Plant metabolites were analyzed by liquid chromatography tandem mass spectrometry (LC-MS/MS) with UHPLC reverse phase chromatography performed using an Agilent 1290 LC coupled with a QExactive Orbitrap MS (QE=139) (Thermo Scientific, San Jose, CA). Samples were kept at 4°C prior to injection. Chromatography was performed using a C18 column (Waters Acquity UPLC BEH, 1.7 um, with VanGuard Pre-Column, 2.1 x 5 mm) at a flow rate of 0.4 mL/min and injection volume of 5 µL, with a column temperature of 60°C. The column was equilibrated with 100% buffer A (100% LC-MS water w/ 0.1% formic acid) for 2 min, following by a linear dilution of buffer A down to 0% with buffer B (100% acetonitrile w/ 0.1% formic acid) over 7 min, and followed by isocratic elution in 100% buffer B for 1.5 min. Full MS spectra were collected ranging from *m*/*z* 80-2,000 at 60,000 to 70,000 resolution in both positive and negative mode, with MS/MS fragmentation data acquisition using an average of stepped 10- 20-40 and 20-50-60 eV collision energies at 17,500 resolution. For targeted analysis, product identification by comparison to standards was performed where authentic standards were available. When possible, standards were weighed and serially diluted to generate standard curves for quantification.

For untargeted analysis, exact mass and retention time coupled with MS/MS fragmentation spectra were used to identify compounds. Features - high intensity signals narrowly contained at a given retention time and *m*/*z* - were detected using the MZMine software version 2.24 (http://dx.doi.org/10.1093/bioinformatics/btk039). First, in MZMine, peaks were detected using a centroid method, removing background peaks below 1 x 10^5^ for MS1 and 1 x 10^3^ for MS2. Chromatograms were built using the ADAP Chromatogram builder, with a minimum 5 scans per peak, a 10 ppm m/z tolerance, and group intensity threshold of 3 x 10^5^. Peaks were deconvoluted, isotopes were grouped, and peaks were aligned between different samples. The resulting feature list was filtered such that a minimum of 2 samples contained the identified feature, and gaps were filled at 10 percent intensity tolerance. Features with a maximum peak height out of all samples less than 3 times the peak height of the corresponding feature in the blank control were discarded as background.

Features that showed a significantly different abundance (peak height) using generalized linear models, calculated using custom R scripts and the lm() function (Team, 2019), with statistical analysis results in Supplemental Table 3 and significantly enriched or depleted features listed in Supplemental Table 2. Generalized linear models are linear regression models used to determine if a particular feature is significantly different in abundance between two genotypes. All features were annotated using Global Natural Products Social Molecular Networking (GNPS) (Katajamaa et al., 2006; Pluskal et al., 2010; Ono et al., 2014; Wang et al., 2016; Nothias et al., 2019). In short, a Feature-Based Molecular Networking workflow was used to assign features to a molecular network with a cosine score above 0.7 and more than six matched peaks. The maximum size of a molecular family was 100, and low scoring edges were removed to meet this threshold. The spectra were then searched against GNPS spectral libraries and annotated with the top hit, if there was one. Complete annotations, features present, and Cytoscape visualization networks are available online for positive mode (https://gnps.ucsd.edu/ProteoSAFe/status.jsp?task=b55e139a9644496aa65aed179858ada7) and negative mode

(https://gnps.ucsd.edu/ProteoSAFe/status.jsp?task=417de81fa3e04daf8e72d3465fe208d6). All scripts are available on GitHub (https://github.com/kmurphy61/ksl4/tree/main). Lists of significantly different features enriched or depleted in each sample type are available in Supplemental Table 2.

### NMR analysis

For structural elucidation of dolabralexin metabolites, compounds were produced in *E. coli* as described above, but scaled to 1 L cultures and concentration of hexane extracts using a rotary evaporator. Approximately 12 cultures were used to produce and purify sufficient compound amounts, followed by purification using silica column chromatography with a hexane:ethyl acetate gradient as previously described (Murphy et al., 2019). Products were controlled for purity using GC-MS, and high-quality samples were pooled and again concentrated by rotary evaporation. Compounds were then further purified to ≥98% purity by HPLC (Agilent 1100 Series) equipped with a diode array UV detector, and an Agilent ZORBAX Eclipse C18 semi-preparative column. A 4 mL/min flow-rate of an acetonitrile:water gradient as mobile phase was used. Purified products were confirmed via GC-MS, concentrated under N_2_ stream, and dissolved in 0.5 mL of deuterated chloroform. NMR spectra were acquired on a Bruker Avance III 800 spectrometer with 5-mm triple resonance cry probe at 25°C. Acquired spectra, including 1D ^1^H, 2D HSQC, correlation spectroscopy, HMBC, and 1D ^13^C (201 MHz) were performed using TopSpin 3.2 software and default parameters. Deuterated chloroform was used as a reference for chemical shift calculations (^13^C 77.23 ppm, ^1^H 7.24 ppm).

### Root architecture analysis

*Zmksl4* and WT maize seeds were planted in potting mix filled germination trays and at four days old were transplanted into 18-inch tall 6-inch-wide tree pots. The pots were filled with turface MVP pre-calibrated to either well-watered status or to a matric water potential of -0.7 MPa. Calibration was accomplished by mixing the turface with water to a specific water content in a separate tub before filling the pots. Following this, each pot was weighed, and the hydration level was maintained throughout the experiment by watering each pot up to weight daily. Well-watered plants were watered to the drip point and could freely drain. Matric potential was monitored hourly using TEROS21 sensors (METER Group, Inc. USA, Pullman, WA) to assure acceptable levels of stress were being imposed. Four pots per treatment were equipped with TEROS21 sensors. At harvest, each root system was carefully removed from the pots and all the turface was gently washed off using water. Roots were then imaged using the DIRT analysis software (Das et al., 2015).

Mature root crowns were harvested from *Zmksl4* and WT maize plants grown to maturity (i.e. seed harvest) at a maize field site in Stanford, CA. Root crowns were harvested by manually pulling plants from the soil, and removing the plant 6 inches from the root/shoot junction and above. Roots were shaken manually to remove large soil clumps, washed clean in a bucket of water, and air dried at room temperature. As above, roots were then imaged using the DIRT analysis software (Das et al., 2015). PCA, ANOVA and t-test analyses for all root architecture were conducted in OriginLab 2020.

### RNAseq analysis

Total maize RNA was extracted from frozen, homogenized plant tissue prepared as outlined above using a Qiagen RNEasy Plant Mini Kit and associated instructions, including an on-column DNAse treatment (Qiagen RNAse-free DNAse set). These were the same plants used for LC-MS/MS V12: n = 5 (WT, well-watered), n = 4 (WT, water deficit), n = 4 (*Zmksl4*, well- watered), n = 5 (*Zmksl4*, water deficit). Library preparation and sequencing were performed by the Vincent J. Coates Genomics Sequencing Laboratory and Functional Genomics Laboratory (UC Berkeley, CA, USA) according to standard protocols. First, RNA was quantified and quality-tested on a Femto Pulse; RNA Quality Number (RQN) were between 4.6 and 8.2. Kapa Biosystems library preparation kits and custom Unique Dual Indexes were used for library preparation. Sequencing was performed on Illumina NovaSeq 6000 150PE Flow Cell S4 (1% PhiX control run) with paired-end sequencing. Raw sequencing data is available via National Institutes of Health Sequence Read Archive (SRA), SRA archive submission number BioProjectID: PRJNA810958. The percent quality score of 30 or higher (%QC30) were between 90.44 and 93.8 percent. The mean quality scores ranged from 35.15 to 35.91, and yield was between 7,299 and 62.282 Mbases. The clusters passing filter (%PF) was 100 percent for every sample, and the percent perfect barcode was between 87.57 and 98.21 percent. For RNAseq analysis, the raw reads were paired and adapters were trimmed using the trim_galore function and low-quality reads removed based on standard parameters. Specifically, reads were mapped to the maize v4 reference (B73 RefGen_v4) genome using a Burrows-Wheeler Alignment (BWA mem) alignment, with a mapping quality cutoff of 40 to reduce multimapping, with samples ranging between 15,941,503 and 145,800,405 high-quality mapped reads per sample. Reads were counted using htseq for a total of 47,289 aligned genes (Dobin et al., 2013; Li, 2013; Portwood et al., 2019). Downstream analysis was performed using the DESeq2 package in R (Anders et al., 2013), with all code available in GitHub (https://github.com/kmurphy61/ksl4/tree/main). Specifically, the DESeq2 package with “ashr” was used for calculating log fold change using shrinkage (Stephens, 2017). Resulting DEG lists were aligned with B73 v3 gene IDs and, where available, GO terms, classical gene names, and orthologs were annotated. All resulting lists of DEGs and associated statistics are listed in Supplemental Table 5.

For Gene Ontology (GO) term enrichment analysis, GO terms were analyzed using GO Ontology database (DOI: 10.5281/zenodo.4735677 Released 2021-05-01) and the PANTHER Overrepresentation Test (Released 20210224) for up- and down-regulated DEG v4 IDs, separately. The Zea mays (all genes in database) was used as a reference list and GO biological processes (complete) was used as the annotation data set. Fisher’s exact test with a Bonferroni correction for multiple testing was employed.

### Analysis of publicly available transcriptomes

RNAseq data was accessed from published datasets, detailed in the respective publications (Chen et al., 2014; Stelpflug et al., 2016; Tai et al., 2016; Walley et al., 2016; Kremling et al., 2018; Yi et al., 2019). Custom R scripts, available at https://github.com/kmurphy61/ksl4/tree/main with all R scripts for this study, were used to filter data for the pathways of interest, with Gene IDs available in Supplemental Table 1. FPKM or RPKM values were plotted using the ggplot2 package in R.

## Supplemental Data

Supplemental Table 1: *Zea mays* V4 gene IDs used in this study and their abbreviations. Abbreviations include: CPS = copalyl pyrophosphate synthase, KSL = kaurene synthase-like, CYP = cytochrome P450, ZmCPS1/Anther Ear 1 (ZmAn1), ZmCPS2/Anther Ear 2 (ZmAn2), ZmKSL2 (ent-isokaurene synthase), (ZmKSL4) dolabradiene synthase, kaurene oxidase 1 (ZmKO1 / ZmCYP701A26), kaurene oxidase 2 (ZmKO2 / ZmCYP701A43), cytochrome P450 reductase (ZmCPR2), kauralexin reductase 2 (ZmKR2). See (DOI: 10.1038/s41477-019-0509-6; kauralexins) and (DOI: 10.1104/pp.17.01351; dolabralexins) for further information on listed maize biosynthetic genes.

Supplemental Table 2A: Metabolite features for peaks enriched in WT, V12 root samples at both well-watered and water deficit conditions (i.e. analyzed separately for each water status, then enriched in both), p<0.1 using a linear model for peak height. Peaks were inspected manually for identity to determine if they had the same retention time, or were present in an authentic DX standard. Peaks without MS2 in WT V12 roots were removed. Features where the highest peak of all samples was still less than 3 times the blank were discarded. Code used for analysis available at https://github.com/kmurphy61/ksl4/tree/main. Sample sizes as follows: V5: n = 4 (WT, well-watered), n = 7 (WT, water deficit), n = 7 (*Zmksl4*, well-watered), n = 6 (*Zmksl4*, water deficit). V12: n = 5 (WT, well-watered), n = 4 (WT, water deficit), n = 5 (*Zmksl4*, well-watered), n = 6 (*Zmksl4*, water deficit). BH = Benjamini-Hochberg Procedure for p-value adjustment.

Supplemental Table 2B: Metabolite features enriched in V5 or V12 roots under well-watered or water-deficit conditions (analyzed separately) due to mutation, p<0.1 using a linear model for peak height. Features where the highest peak of all samples was still less than 3 times the blank were discarded. Code used for analysis available at https://github.com/kmurphy61/ksl4/tree/main. Sample sizes as follows: V5: n = 4 (WT, well- watered), n = 7 (WT, water deficit), n = 7 (*Zmksl4*, well-watered), n = 6 (*Zmksl4*, water deficit). V12: n = 5 (WT, well-watered), n = 4 (WT, water deficit), n = 5 (*Zmksl4*, well-watered), n = 6 (*Zmksl4*, water deficit). BH = Benjamini-Hochberg Procedure for p-value adjustment.

Supplemental Table 2C: Metabolite features enriched in V5 or V12 roots in *Zmksl4* or WT (analyzed separately) due to water status, p<0.1 using a linear model for peak height. Features where the highest peak of all samples was still less than 3 times the blank were discarded. Code used for analysis available at https://github.com/kmurphy61/ksl4/tree/main. Sample sizes as follows: V5: n = 4 (WT, well-watered), n = 7 (WT, water deficit), n = 7 (*Zmksl4*, well-watered), n = 6 (*Zmksl4*, water deficit). V12: n = 5 (WT, well-watered), n = 4 (WT, water deficit), n = 5 (*Zmksl4*, well-watered), n = 6 (*Zmksl4*, water deficit). BH = Benjamini-Hochberg Procedure for p-value adjustment.

Supplemental Table 2D: Metabolite features enriched in V5 or V12 leaves under well-watered or water-deficit conditions (analyzed separately) due to mutation, p<0.1 using a linear model for peak height. Features where the highest peak of all samples was still less than 3 times the blank were discarded. Code used for analysis available at https://github.com/kmurphy61/ksl4/tree/main. Sample sizes as follows: V5: n = 4 (WT, well-watered), n = 7 (WT, water deficit), n = 7 (*Zmksl4*, well-watered), n = 6 (*Zmksl4*, water deficit). V12: n = 5 (WT, well-watered), n = 4 (WT, water deficit), n = 5 (*Zmksl4*, well-watered), n = 6 (*Zmksl4*, water deficit). BH = Benjamini-Hochberg Procedure for p-value adjustment.

Supplemental Table 2E: Metabolite features enriched in V5 or V12 leaves in *Zmksl4* or WT (analyzed separately) due to water status, p<0.1 using a linear model for peak height. Features where the highest peak of all samples was still less than 3 times the blank were discarded. Code used for analysis available at https://github.com/kmurphy61/ksl4/tree/main. Sample sizes as follows: V5: n = 4 (WT, well-watered), n = 7 (WT, water deficit), n = 7 (*Zmksl4*, well-watered), n = 6 (*Zmksl4*, water deficit). V12: n = 5 (WT, well-watered), n = 4 (WT, water deficit), n = 5 (*Zmksl4*, well-watered), n = 6 (*Zmksl4*, water deficit). BH = Benjamini-Hochberg Procedure for p-value adjustment.

Supplemental Table 3A: Metabolomics statistics. PERMANOVA analysis of WT and *Zmksl4* leaf and root metabolite features (i.e. peak height), using the LSMEANS package in R. Features where the highest peak of all samples was still less than 3 times the blank were discarded. Code used for analysis available at https://github.com/kmurphy61/ksl4/tree/main. Sample sizes as follows: V5: n = 4 (WT, well-watered), n = 7 (WT, water deficit), n = 7 (*Zmksl4*, well-watered), n = 6 (*Zmksl4*, water deficit). V12: n = 5 (WT, well-watered), n = 4 (WT, water deficit), n = 5 (*Zmksl4*, well-watered), n = 6 (*Zmksl4*, water deficit).

Supplemental Table 3B: Plant weight statistics. PERMANOVA analysis and t-tests of WT and *Zmksl4* fresh root and shoot weights at V12, and their ratio, using the LSMEANS package in R. Code used for analysis available at https://github.com/kmurphy61/ksl4/tree/main. Sample sizes as follows: V12: n = 6 (WT, well-watered), n = 4 (WT, water deficit), n = 5 (*Zmksl4*, well- watered), n = 6 (*Zmksl4*, water deficit). Raw data in Supplemental Table 3C and D. **Supplemental Table 3C:** Plant weight raw data. WT and *Zmksl4* fresh root and shoot weights at V5 and V12, and their ratio, grown in Davis, CA. Code used for analysis available at https://github.com/kmurphy61/ksl4/tree/main. Statistical analysis in Supplemental Table 3B. **Supplemental Table 3D**: Plant weight raw data. WT and *Zmksl4* fresh root and shoot weights at V5, and their ratio, grown in St. Louis, MO greenhouse in turface. Code used for analysis available at https://github.com/kmurphy61/ksl4/tree/main. Statistical analysis in Supplemental Table 3B.

Supplemental Table 3E: Transcriptomics statistics. Statistical analysis of differentially expressed genes (DEGs) in WT and *Zmksl4* root tissue at V12, using the DESeq2 package in R. These were the same samples used for metabolomics and weight analysis in Supplemental Table 2A, B, and C. Code used for analysis available at https://github.com/kmurphy61/ksl4/tree/main. Sample sizes as follows: V12: n = 5 (WT, well-watered), n = 4 (WT, water deficit), n = 4 (*Zmksl4*, well-watered), n = 5 (*Zmksl4*, water deficit). Abbreviations include: DEG = differentially expressed genes; LFC = log fold change.

Supplemental Table 4: Publicly-available transcriptomic data of diterpenoid-related genes used in this study. Sheet labeled “Data Description“ identifies the sample names, descriptions, references, and data source. Subsequent sheets represent gene expression data (FPKM) of selected maize biosynthetic pathway genes. Abbreviations include: CPS = copalyl pyrophosphate synthase, KSL = kaurene synthase-like, CYP = cytochrome P450, ZmCPS1/Anther Ear 1 (ZmAn1), ZmCPS2/Anther Ear 2 (ZmAn2), ZmKSL2 (ent-isokaurene synthase), (ZmKSL4) dolabradiene synthase, kaurene oxidase 1 (ZmKO1/ZmCYP701A26), kaurene oxidase 2 (ZmKO2/ZmCYP701A43), cytochrome P450 reductase (ZmCPR2), kauralexin reductase 2 (ZmKR2). See (DOI: 10.1038/s41477-019-0509-6; kauralexins) and (DOI: 10.1104/pp.17.01351; dolabralexins) for further information on listed maize biosynthetic genes.

Supplemental Table 5A: DEGs (differentially expressed genes) enriched or depleted in water-deficit roots at V12 due to the mutation (i.e. impacted by mutation, up- or down-regulated in *Zmksl4*). DEGs were calculated using the DESeq2 package in R, see Methods for details. Annotations were added when available. These were the same samples used for metabolomics and weight analysis. Code used for analysis available at https://github.com/kmurphy61/ksl4/tree/main. Sample sizes as follows: V12: n = 5 (WT, well- watered), n = 4 (WT, water deficit), n = 4 (*Zmksl4*, well-watered), n = 5 (*Zmksl4*, water deficit).

Supplemental Table 5B: DEGs (differentially expressed genes) enriched or depleted in well- watered roots at V12 due to the mutation (i.e. impacted by mutation, up- or down-regulated in *Zmksl4*). DEGs were calculated using the DESeq2 package in R, see Methods for details. Annotations were added when available. These were the same samples used for metabolomics and weight analysis. Code used for analysis available at https://github.com/kmurphy61/ksl4/tree/main. Sample sizes as follows: V12: n = 5 (WT, well- watered), n = 4 (WT, water deficit), n = 4 (*Zmksl4*, well-watered), n = 5 (*Zmksl4*, water deficit).

Supplemental Table 5C: DEGs (differentially expressed genes) enriched or depleted in water- deficit AND well-watered roots at V12 due to the mutation (i.e. impacted by mutation, up- or down-regulated in *Zmksl4*). DEGs were calculated using the DESeq2 package in R, see Methods for details. Annotations were added when available. These were the same samples used for metabolomics and weight analysis. Code used for analysis available at https://github.com/kmurphy61/ksl4/tree/main. Sample sizes as follows: V12: n = 5 (WT, well- watered), n = 4 (WT, water deficit), n = 4 (*Zmksl4*, well-watered), n = 5 (*Zmksl4*, water deficit).

Supplemental Table 5D: DEGs (differentially expressed genes) enriched or depleted in roots at V12 due to the interaction factor between water status x genotype. DEGs were calculated using the DESeq2 package in R, see Methods for details. Annotations were added when available. These were the same samples used for metabolomics and weight analysis in Supplemental Table 2A, B, and C. Code used for analysis available at https://github.com/kmurphy61/ksl4/tree/main. Sample sizes as follows: V12: n = 5 (WT, well-watered), n = 4 (WT, water deficit), n = 4 (*Zmksl4*, well-watered), n = 5 (*Zmksl4*, water deficit).

Supplemental Table 5E: DEGs (differentially expressed genes) enriched or depleted in *Zmksl4* roots V12 due to the water status (i.e. impacted by water status, up- or down-regulated in water deficit roots). DEGs were calculated using the DESeq2 package in R, see Methods for details. Annotations were added when available. These were the same samples used for metabolomics and weight analysis. Code used for analysis available at https://github.com/kmurphy61/ksl4/tree/main. Sample sizes as follows: V12: n = 5 (WT, well- watered), n = 4 (WT, water deficit), n = 4 (*Zmksl4*, well-watered), n = 5 (*Zmksl4*, water deficit).

Supplemental Table 5F: DEGs (differentially expressed genes) enriched or depleted in WT roots V12 due to the water status (i.e. impacted by water status, up- or down-regulated in water deficit roots). DEGs were calculated using the DESeq2 package in R, see Methods for details. Annotations were added when available. These were the same samples used for metabolomics and weight analysis. Code used for analysis available at https://github.com/kmurphy61/ksl4/tree/main. Sample sizes as follows: V12: n = 5 (WT, well- watered), n = 4 (WT, water deficit), n = 4 (*Zmksl4*, well-watered), n = 5 (*Zmksl4*, water deficit).

Supplemental Table 5G: DEGs (differentially expressed genes) enriched or depleted in *Zmksl4* AND WT roots at V12 due to the water status (i.e. impacted by water status, up- or down- regulated in water-deficit conditions). DEGs were calculated using the DESeq2 package in R, see Methods for details. Annotations were added when available. These were the same samples used for metabolomics and weight analysis. Code used for analysis available at https://github.com/kmurphy61/ksl4/tree/main. Sample sizes as follows: V12: n = 5 (WT, well- watered), n = 4 (WT, water deficit), n = 4 (*Zmksl4*, well-watered), n = 5 (*Zmksl4*, water deficit).

Supplemental Table 6A: GO (gene ontology) terms enriched for DEGs (differentially expressed genes) that were up-regulated in WT roots at V12 due to water deficit (i.e. higher expression in water deficit roots). DEGs were calculated using the DESeq2 package in R, see Methods for details. These were the same samples used for metabolomics and weight analysis. Code used for analysis available at https://github.com/kmurphy61/ksl4/tree/main. Sample sizes as follows: V12: n = 5 (WT, well-watered), n = 4 (WT, water deficit).

Supplemental Table 6B: GO (gene ontology) terms enriched for DEGs (differentially expressed genes) that were down-regulated in WT roots at V12 due to water deficit (i.e. lower expression in water deficit roots). DEGs were calculated using the DESeq2 package in R, see Methods for details. These were the same samples used for metabolomics and weight analysis. Code used for analysis available at https://github.com/kmurphy61/ksl4/tree/main. Sample sizes as follows: V12: n = 5 (WT, well-watered), n = 4 (WT, water deficit).

Supplemental Table 6C: GO (gene ontology) terms enriched for DEGs (differentially expressed genes) that were up-regulated in *Zmksl4* roots at V12 due to water deficit (i.e. higher expression in water deficit roots). DEGs were calculated using the DESeq2 package in R, see Methods for details. These were the same samples used for metabolomics and weight analysis. Code used for analysis available at https://github.com/kmurphy61/ksl4/tree/main. Sample sizes as follows: V12: n = 4 (*Zmksl4*, well-watered), n = 5 (*Zmksl4*, water deficit).

Supplemental Table 6D: GO (gene ontology) terms enriched for DEGs (differentially expressed genes) that were down-regulated in *Zmksl4* roots at V12 due to water deficit (i.e. lower expression in water deficit roots). DEGs were calculated using the DESeq2 package in R, see Methods for details. These were the same samples used for metabolomics and weight analysis. Code used for analysis available at https://github.com/kmurphy61/ksl4/tree/main. Sample sizes as follows: V12: n = 4 (*Zmksl4*, well-watered), n = 5 (*Zmksl4*, water deficit).

Supplemental Figure 1: Metabolite atlas for LC-MS/MS with available standards. Upper spectra is peak in a representative root sample, lower is peak in standard. See Methods for details on LC-MS/MS methods and standard preparation.

Supplemental Figure 2: Supplemental Figure 2: LC-MS/MS spectra of new DA metabolites (i.e. not Supplemental Figure 1 for which standards were available) enriched in WT roots, but absent in *Zmksl4*, at V12 and for which MS/MS spectra was available in at least one WT V12 sample. Statistics and peak intensity available in Supplemental Table 1.

Supplemental Figure 3: Boxplots of feature abundance of DA candidates (i.e. the features commonly enriched in V12 WT root samples for which MS2 were available) in (A) root samples and (B) leaf samples.

Supplemental Figure 4: NMR analysis of dolabradienol.

Supplemental Figure 5: Root metabolic features (LC-MS/MS) affected by water status in *Zmksl4* and WT. Few features are commonly affected in both genotypes. Features enriched in (A) water deficit samples and (B) well-watered samples due to water status. Significant features determined by a generalized linear model, P <= 0.1.

Supplemental Figure 6: Root and shoot weights for plants used in root architecture experiment. WS = water stressed, WW = well-watered, R:S = rot/shoot ratio.

Supplemental Figure 7: Global transcriptome changes in WT and Zmksl4. RNA sequencing shows mutation has a minor effect on global transcriptome of Zmksl4. (A) Principal component analysis of all sample types. (B) Sample-to-sample distances, each row represents a biological replicate.

## Author Contributions

KMM, EAS, and PZ designed the experiments; KMM performed most experiments; TD and KMM performed root architecture experiments; AK and EAS performed select metabolite extraction experiments; BY generated mutant plants; BE and PS assisted with metabolomics analysis. KMM and PZ wrote the article with contributions of all other authors.

## Acknowledgements

The authors gratefully acknowledge and Dr. Reuben Peters (Iowa State University) for providing the pIRS construct.

