## Supplemental Figure 1 for "A dolabralexin-deficient mutant provides insight into specialized diterpenoid metabolism in maize (*Zea mays*)"

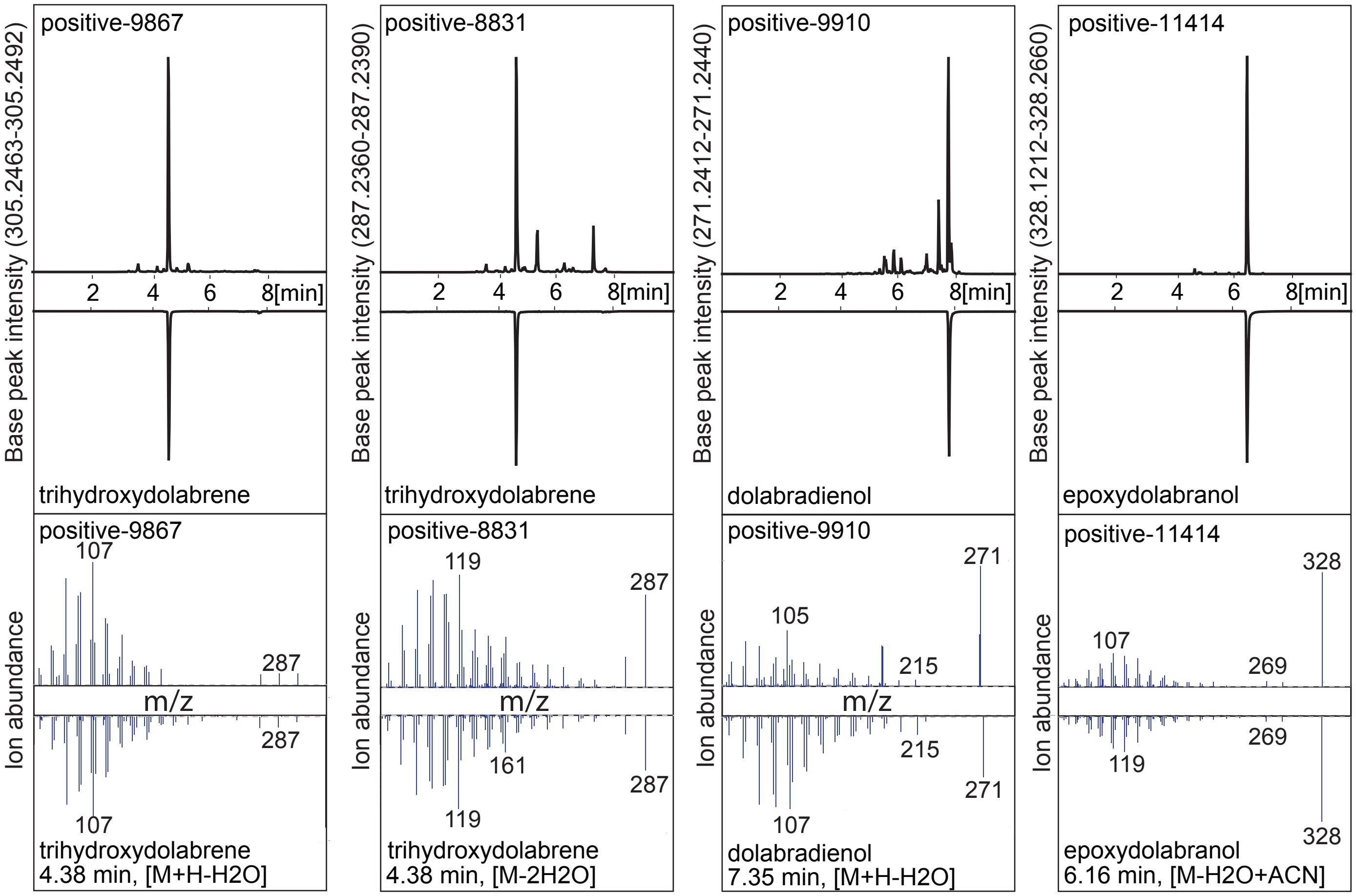


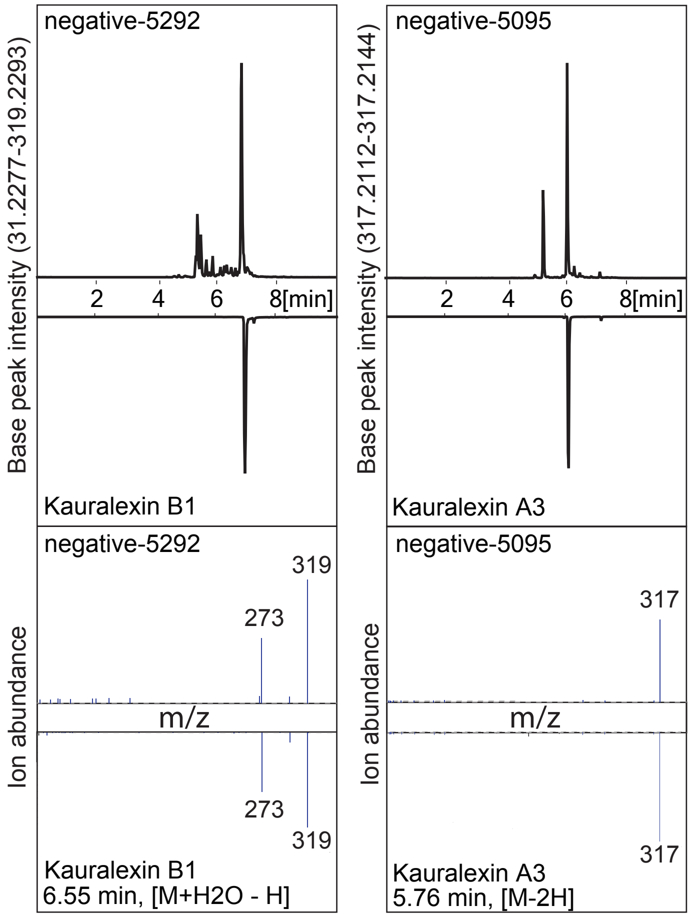


Supplemental Figure 1: Metabolite atlas for LC-MS/MS with available standards. Upper spectra is peak in a representative root sample, lower is peak in standard. See Methods for details on LC-MS/MS methods and standard preparation.
