## Supplemental Figure 2 for "A dolabralexin-deficient mutant provides insight into specialized diterpenoid metabolism in maize (*Zea mays*)"

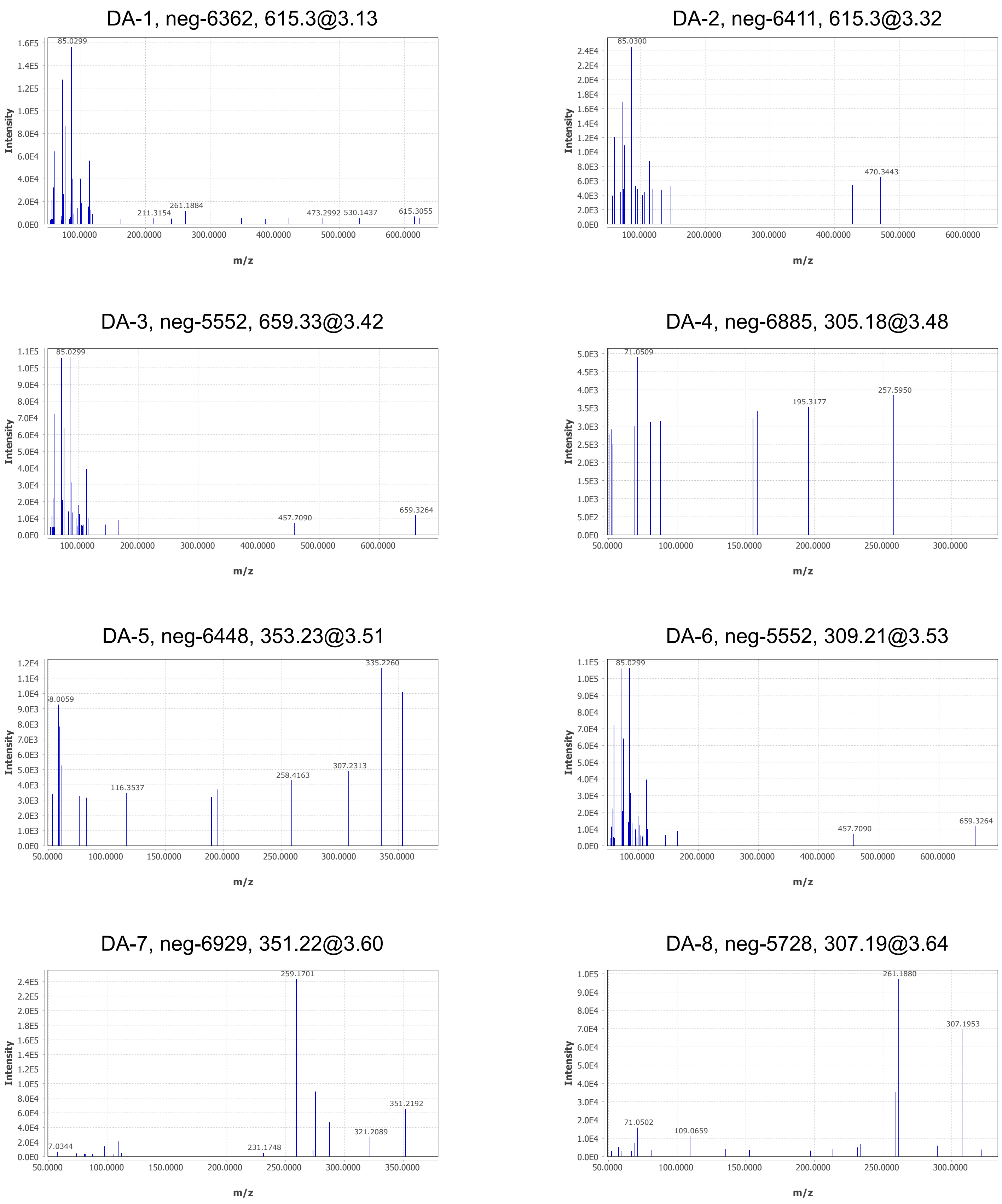


Supplemental Figure 2: LC-MS/MS spectra of new DA metabolites (i.e. not Supplemental Figure 1 for which standards were available) enriched in WT roots, but absent in *Zmksl4*, at V12 and for which MS/MS spectra was available in at least one WT V12 sample. Statistics and peak intensity available in Supplemental Table 2.

### DA-9, neg-5503, 393.23@3.68

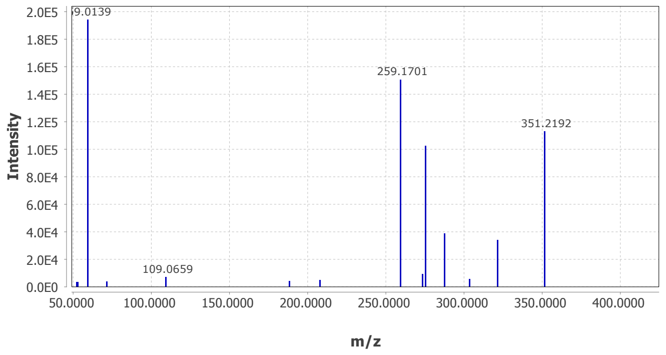


### DA-10, neg-6901, 321.21@3.89

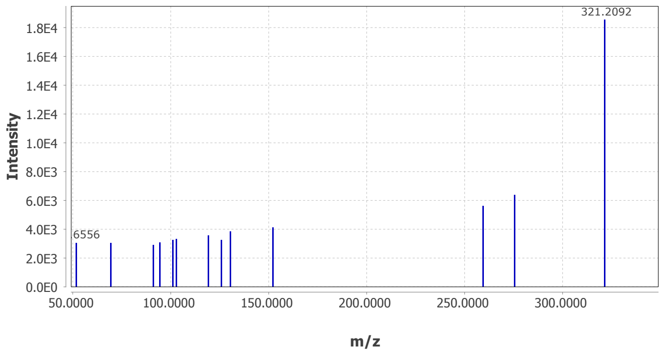


### DA-11, neg-7012, 307.19@3.98

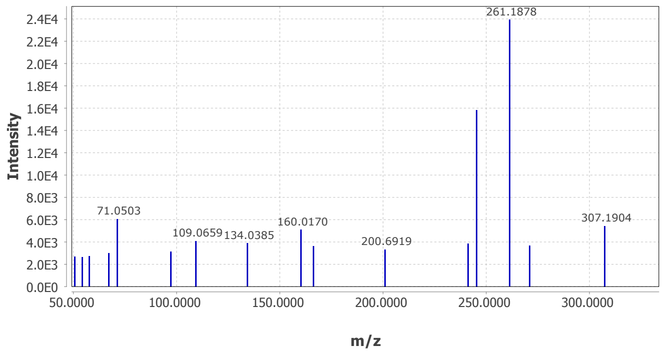


### DA-12, neg-7004, 277.22@4.05

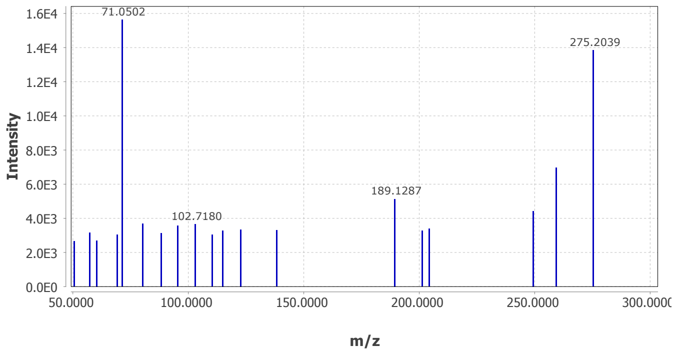


### DA-13, neg-6400, 277.27@4.27

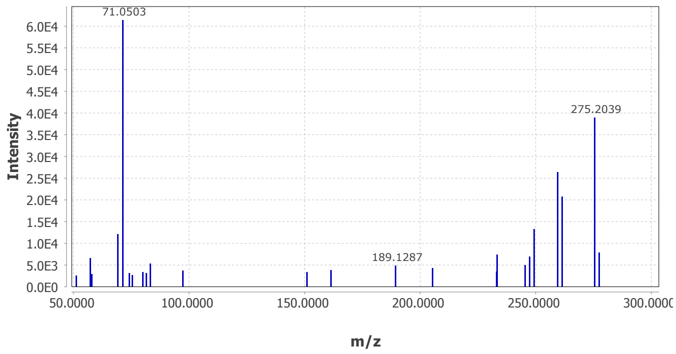


### DA-14, neg-5319, 333.21@3.43

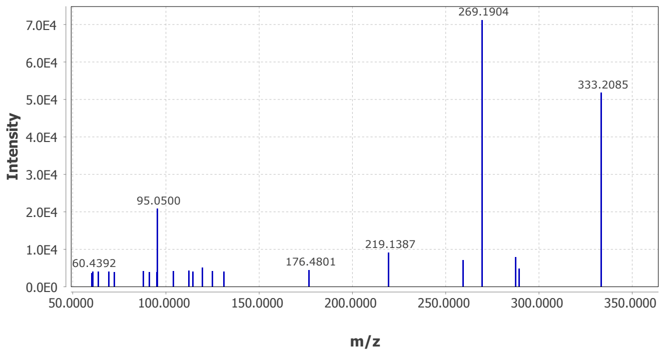


### DA-15, neg-6986, 317.21@6.94

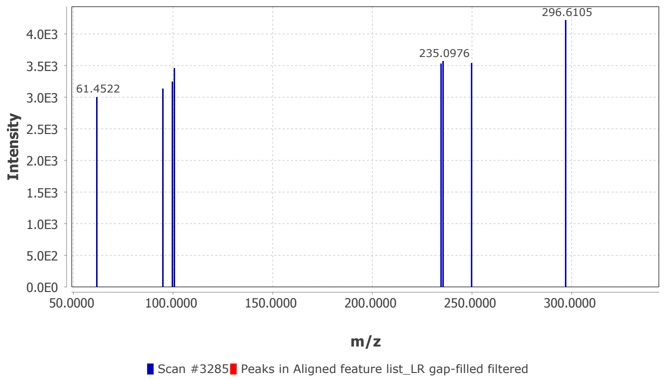

DA-16, pos-11627, 149.04@2.5

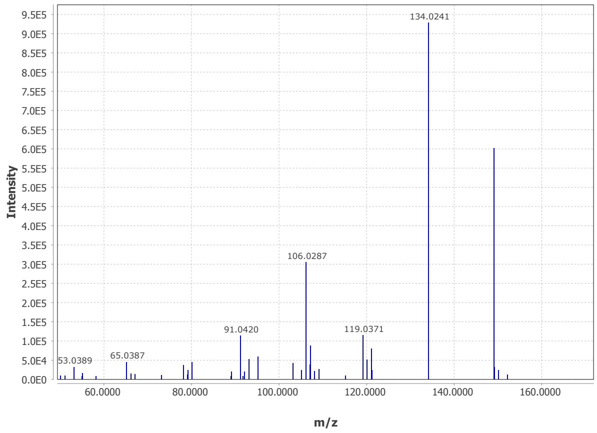


DA-17, pos-10240, 337.24@2.97

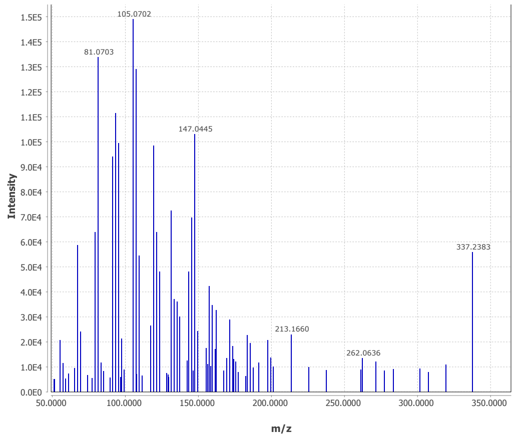


DA-18, pos-11218, 259.21@3.11

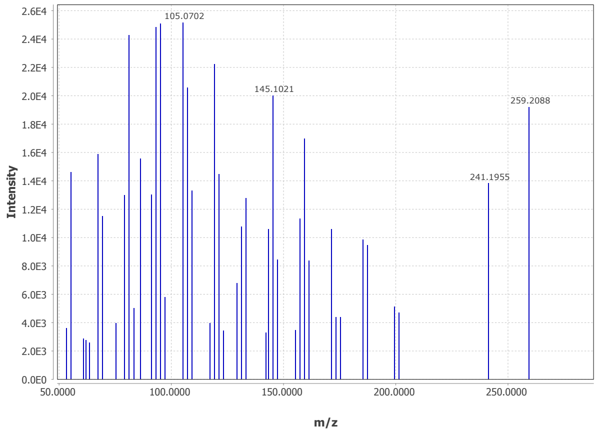


DA-19, pos-11181, 320.26@3.20

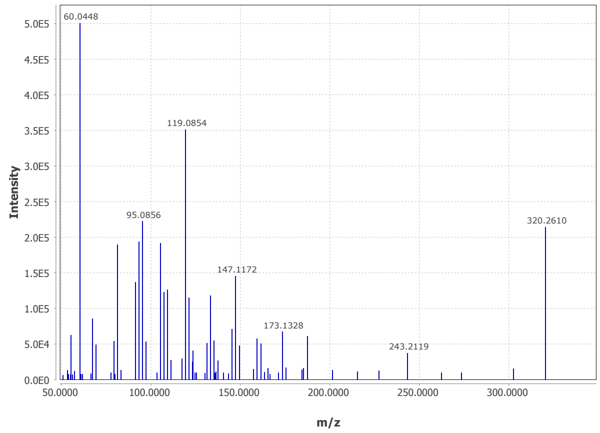


DA-20, pos-8849, 259.21@3.44

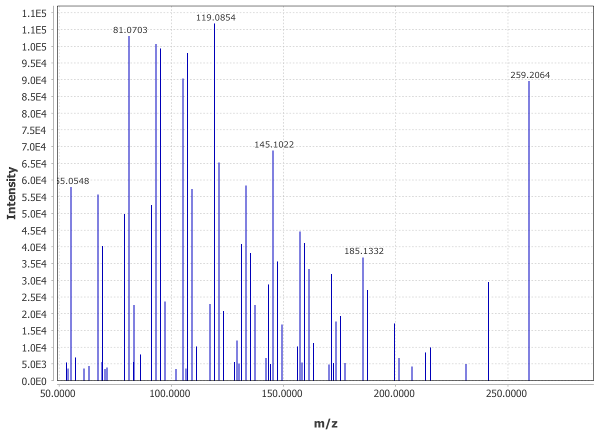


DA-21, pos-8851, 259.21@3.61

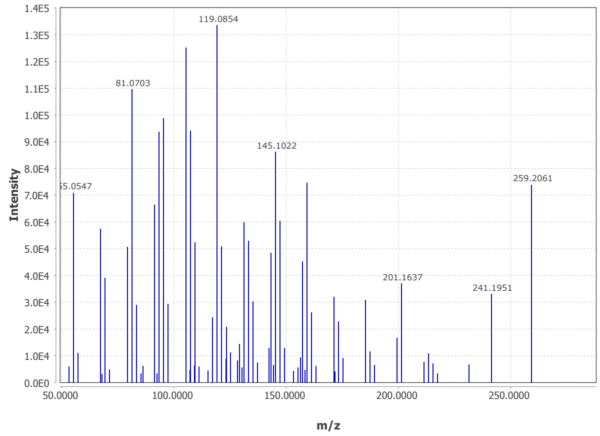


DA-22, pos-9634, 339.25@3.68

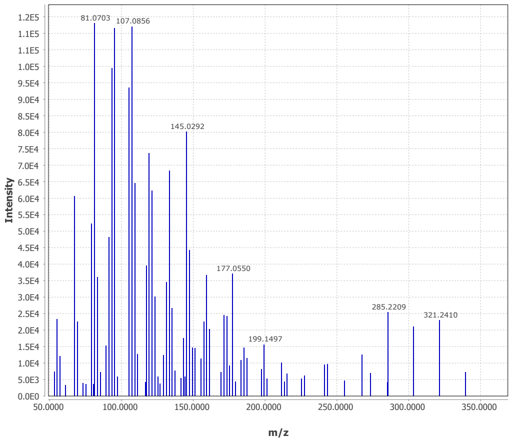


DA-23, pos-10079, 241.2@3.7

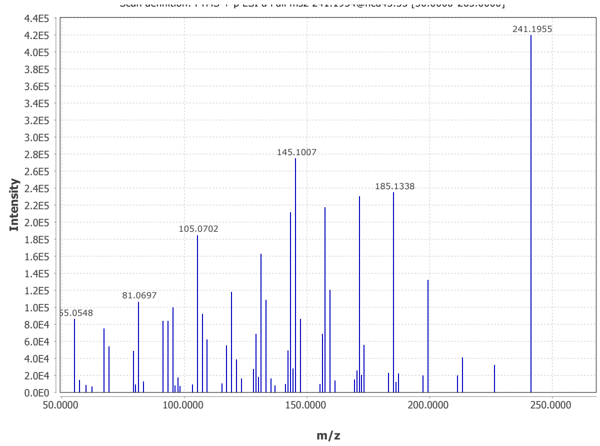

DA-24, pos-7359, 321.24@3.76

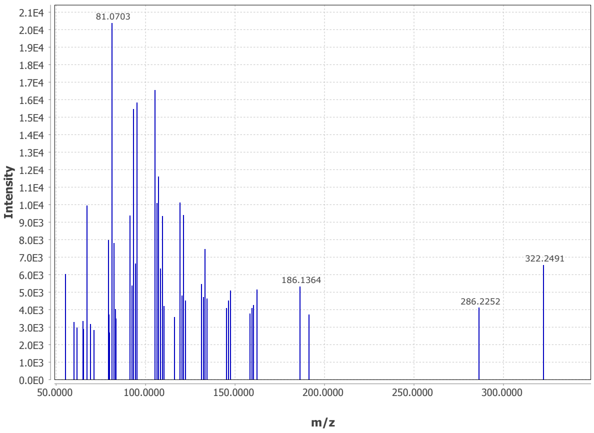


DA-25, pos-9793, 259.21@3.89

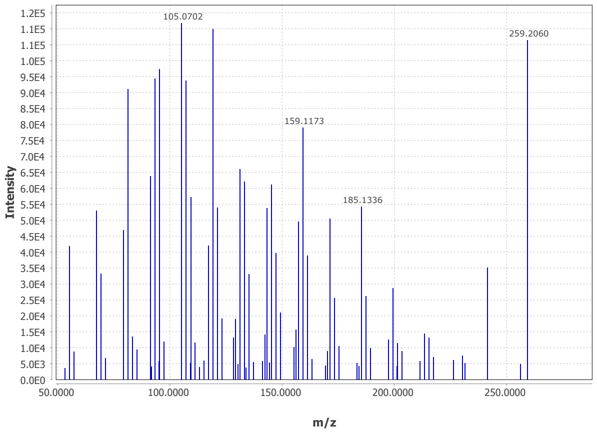


DA-26, pos-3224, 305.25@4.04

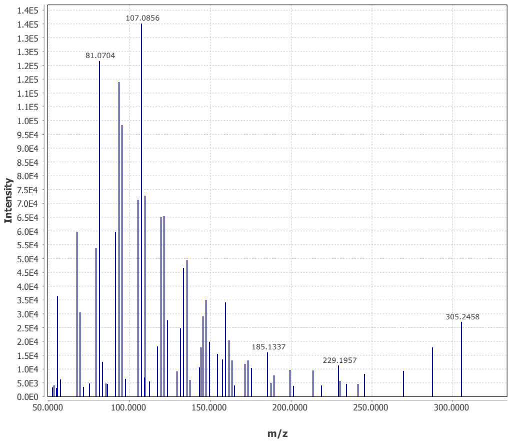


DA-27, pos-1013, 320.26@3.20

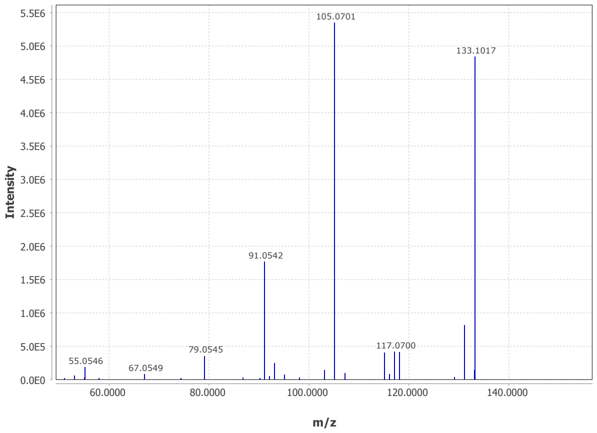


DA-28, pos-9959, 259.21@3.11

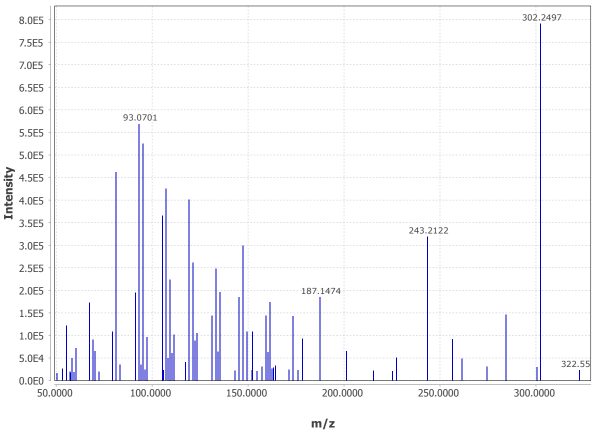


DA-29, pos-10149, 259.21@3.61

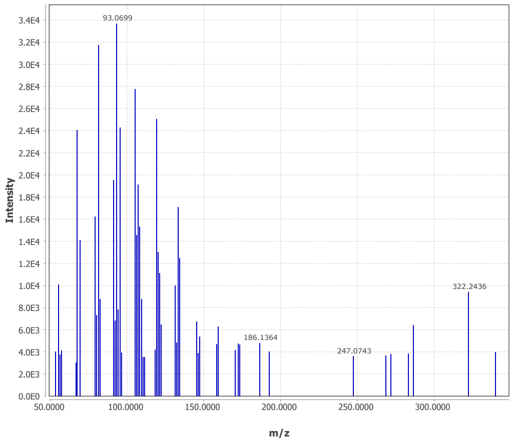


DA-30, pos-1917, 339.25@3.68

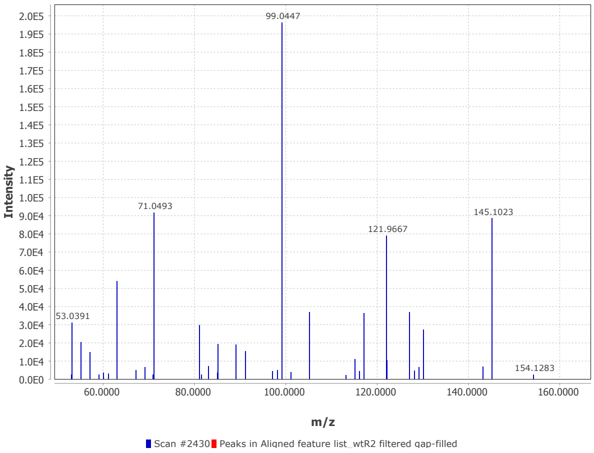


DA-31, pos-11174, 321.24@4.23

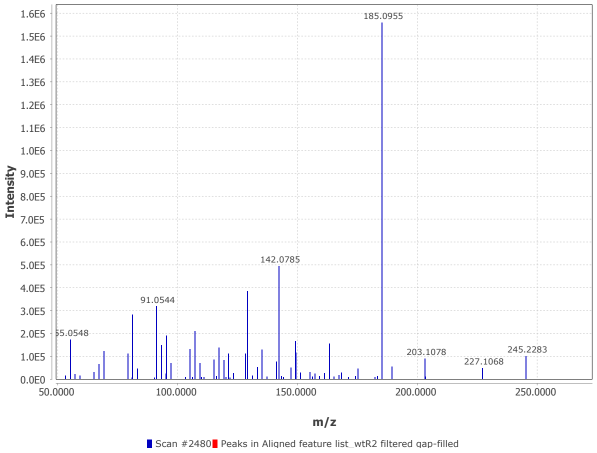

### DA-32, pos-8784, 187.15@4.47

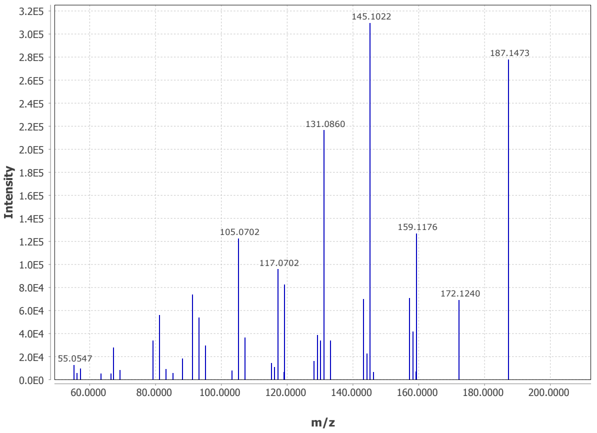


### DA-33, pos-9959, 302.25@4.49

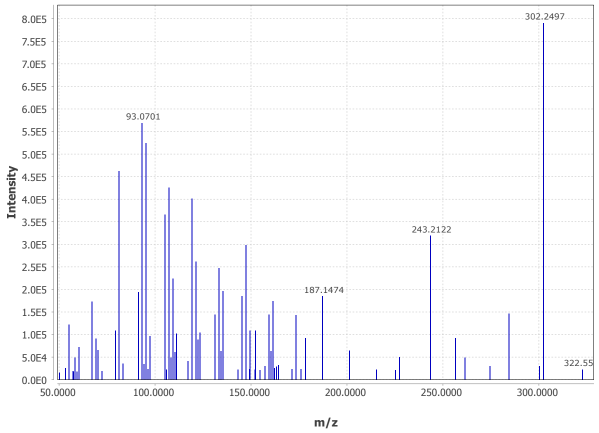


### DA-34, pos-10149, 321.24@4.53

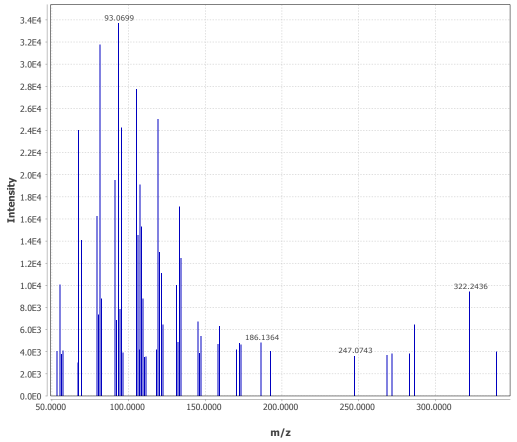


### DA-35, pos-11174, 245.23@4.57

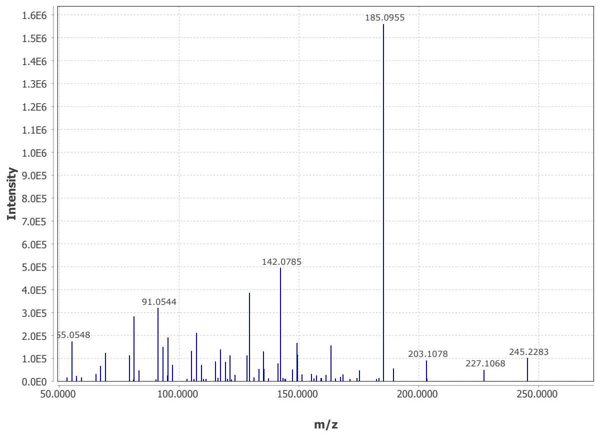


### DA-36, pos-1917, 141.1@4.59

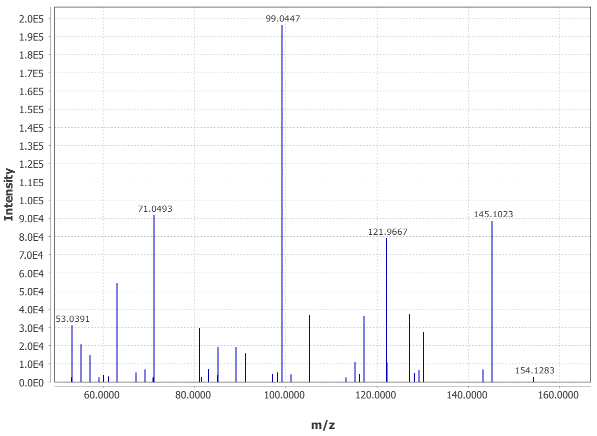


### DA-37, pos-10706, 391.25@4.63

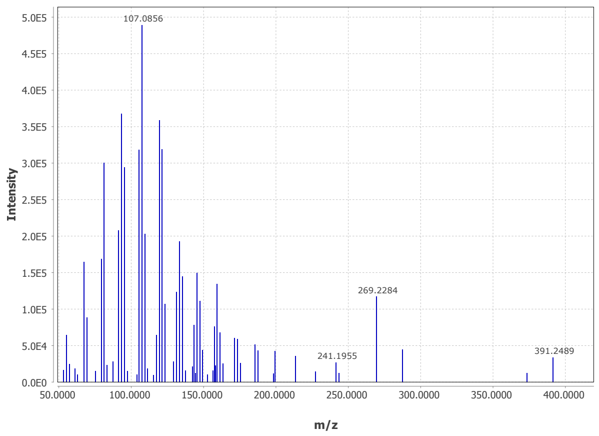


### DA-38, pos-10372, 323.26@4.68

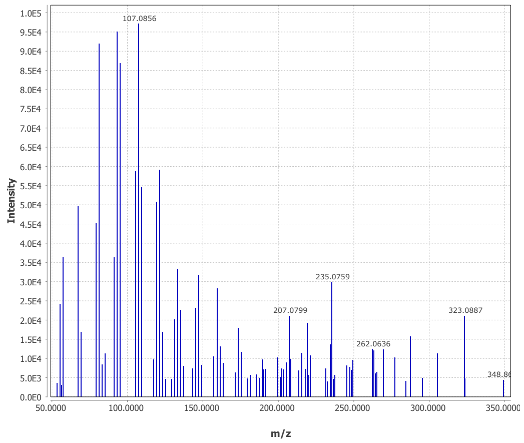


### DA-39, pos-10184, 348.29@4.75

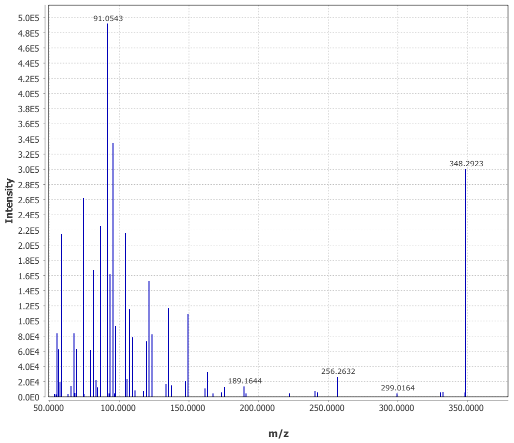

### DA-40, pos-11051, 261.22@4.78

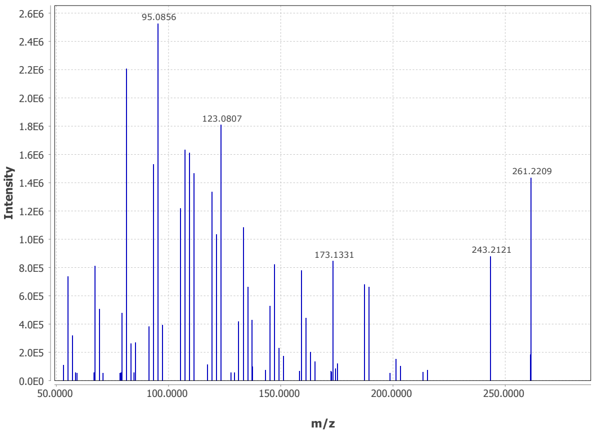


### DA-41, pos-9877, 303.23@4.83

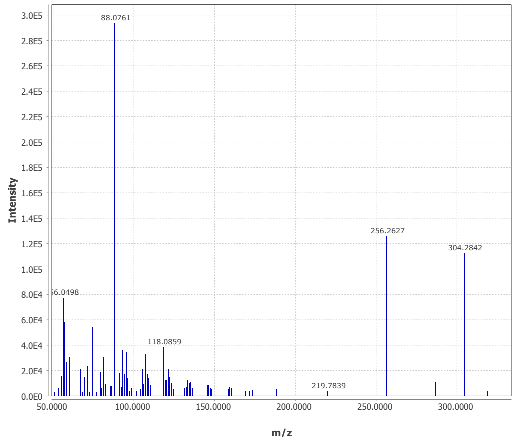


### DA-42, pos-9892, 289.22@4.84

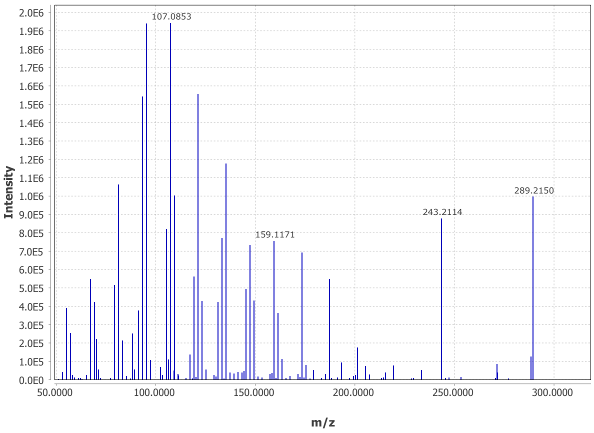


### DA-43, pos-9647, 319.23@4.96

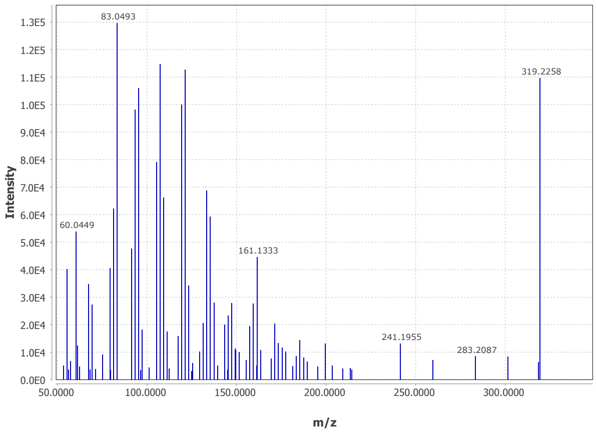


### DA-44, pos-9659, 303.23@5.13

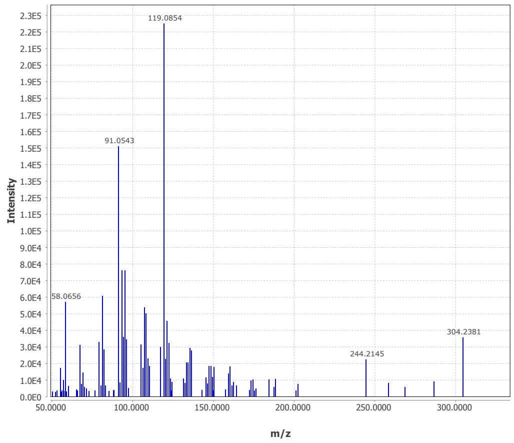


### DA-45, pos-10006, 289.25@5.3

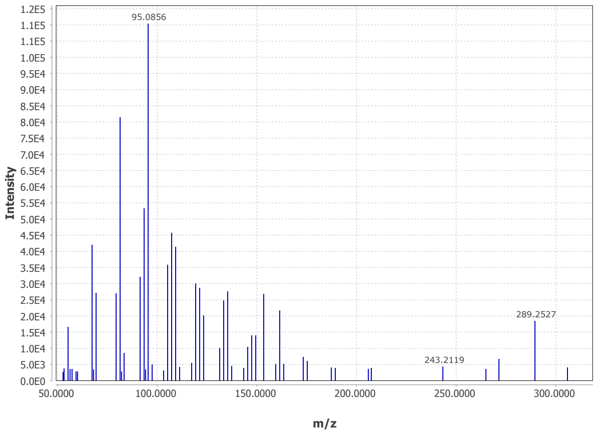


### DA-46, pos-10012, 289.25@6.63

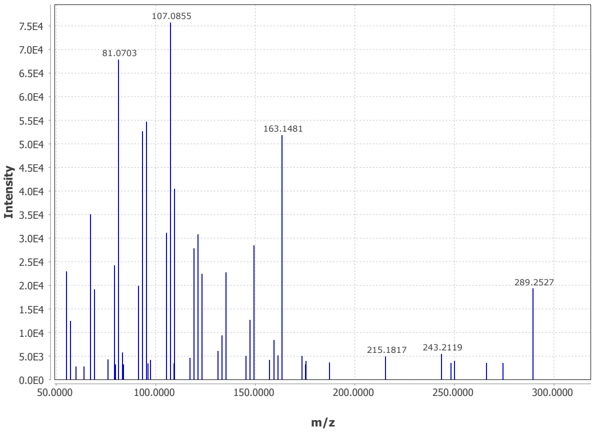


### DA-47, pos-11229, 259.24@7.45

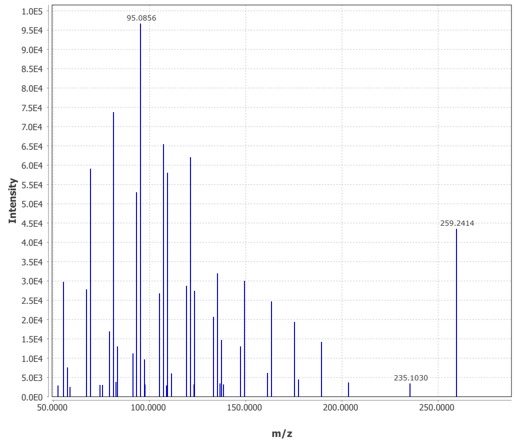
