## Supplemental Figure 3 for "A dolabralexin-deficient mutant provides insight into specialized diterpenoid metabolism in maize (*Zea mays*)"

A

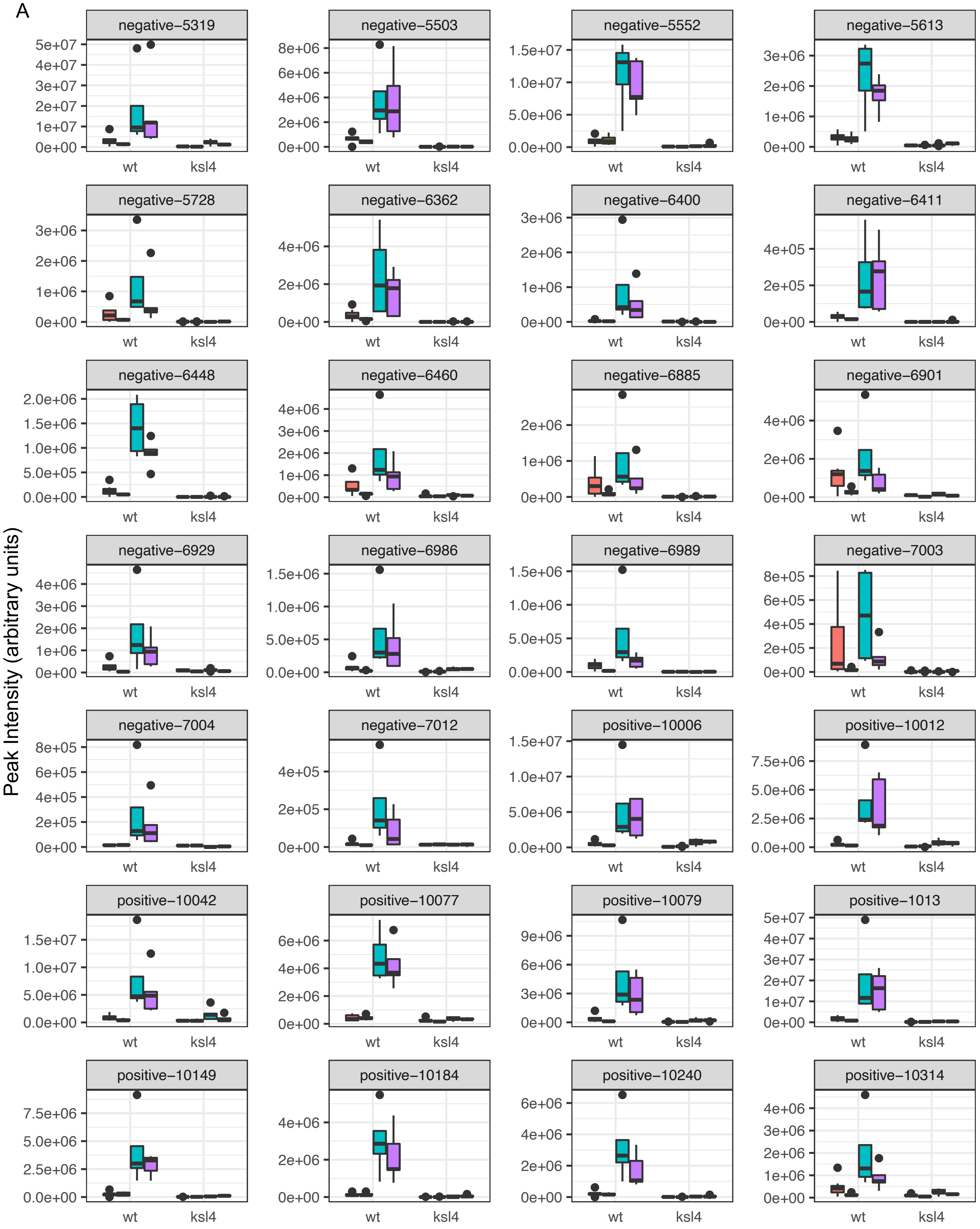


Supplemental Figure 3: Boxplots of feature abundance of DA candidates (i.e. the features commonly enriched in V12 WT root samples for which MS2 were available) in (A) root samples and (B) leaf samples.

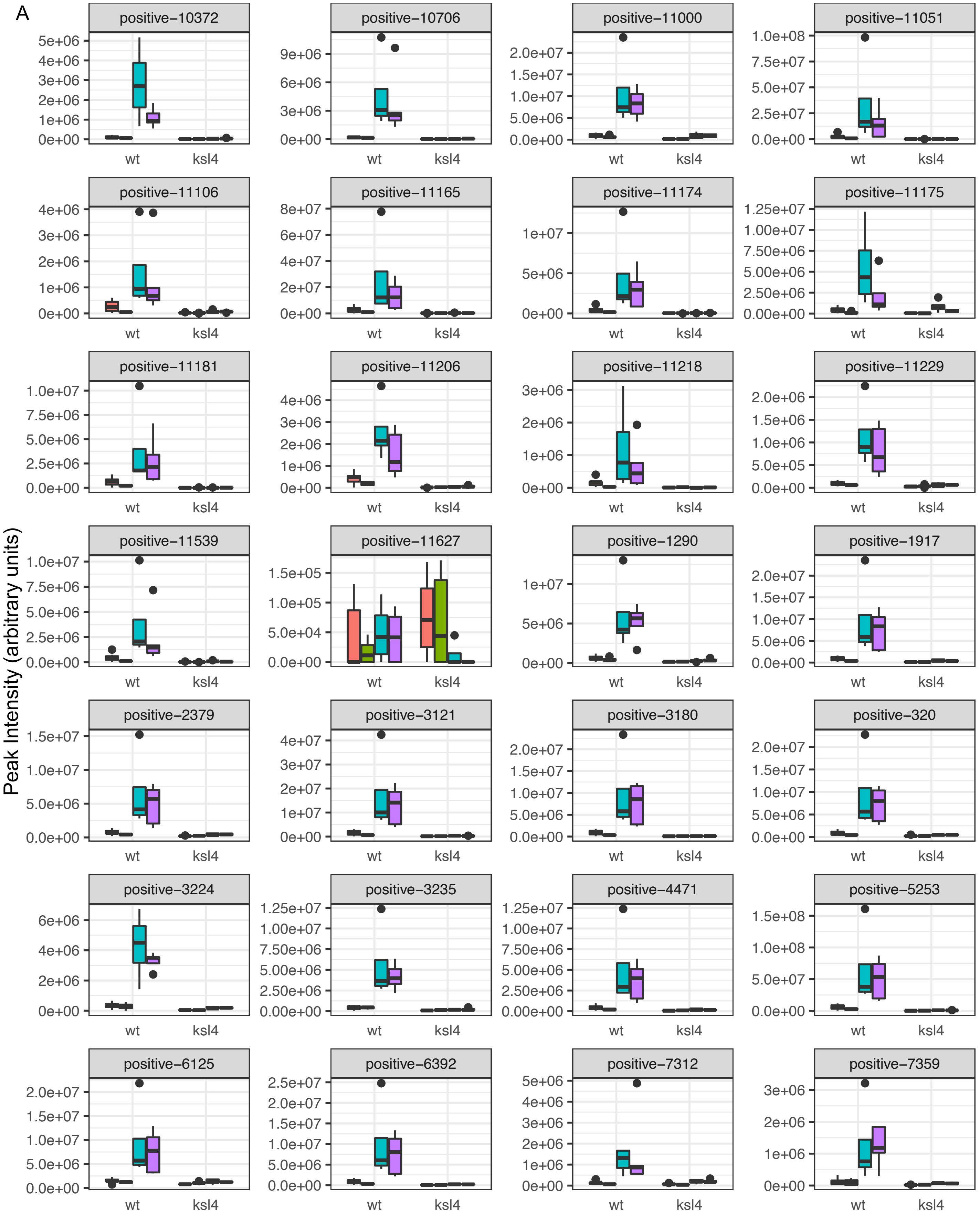


Supplemental Figure 3: Boxplots of feature abundance of DA candidates (i.e. the features commonly enriched in V12 WT root samples for which MS2 were available) in (A) root samples and (B) leaf samples.

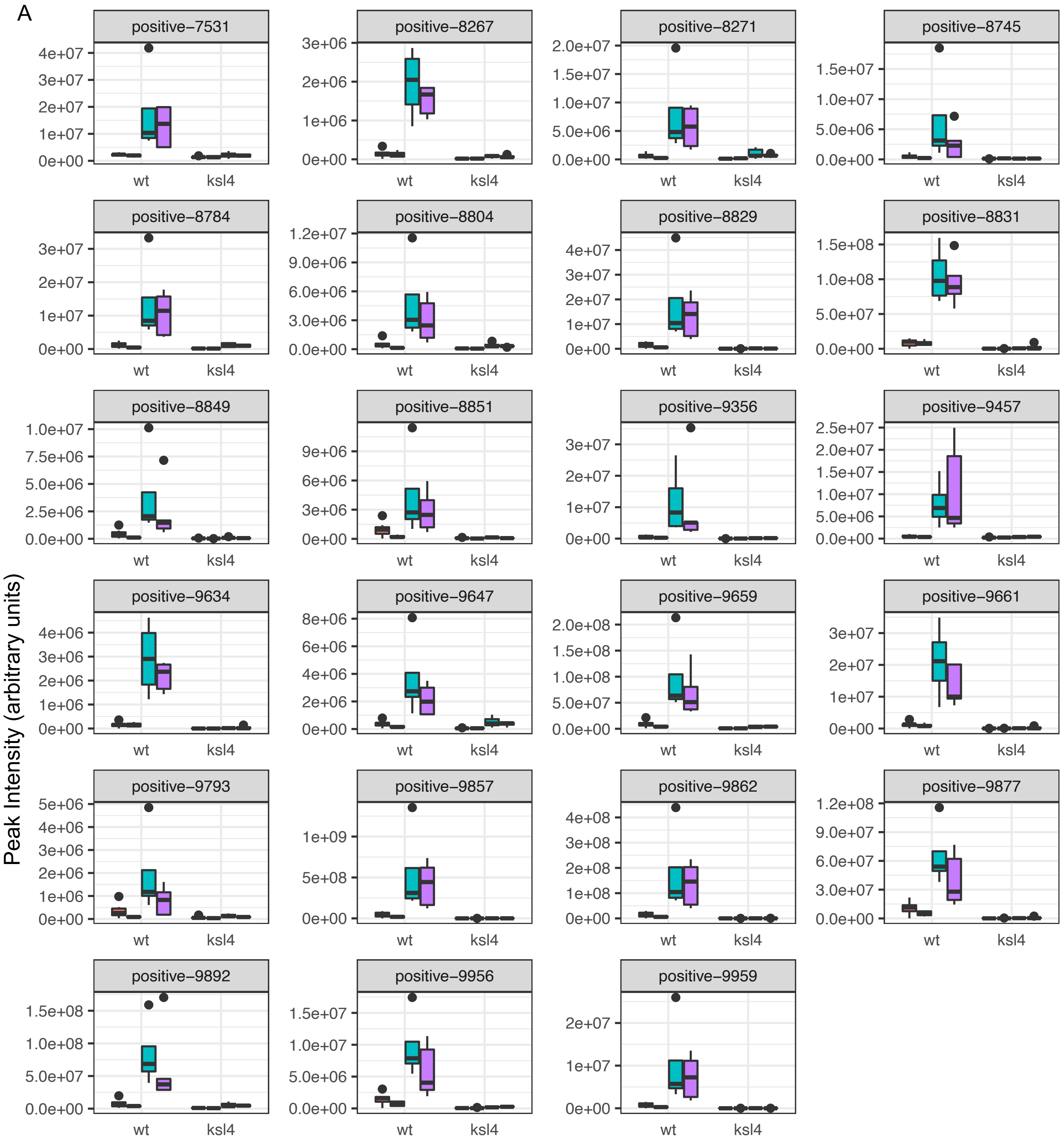


Supplemental Figure 3: Boxplots of feature abundance of DA candidates (i.e. the features commonly enriched in V12 WT root samples for which MS2 were available) in (A) root samples and (B) leaf samples.

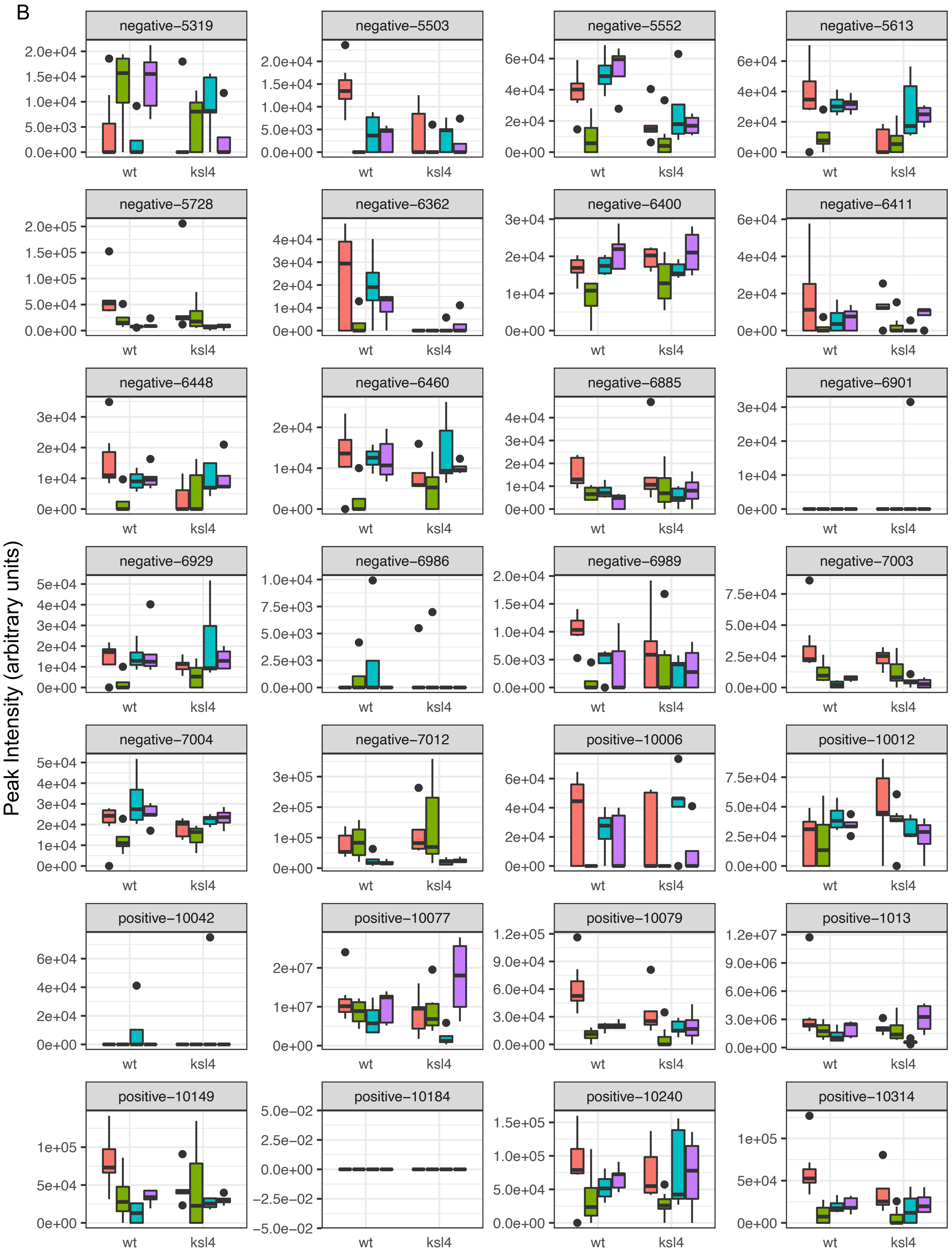


Supplemental Figure 3: Boxplots of feature abundance of DA candidates (i.e. the features commonly enriched in V12 WT root samples for which MS2 were available) in (A) root samples and (B) leaf samples.

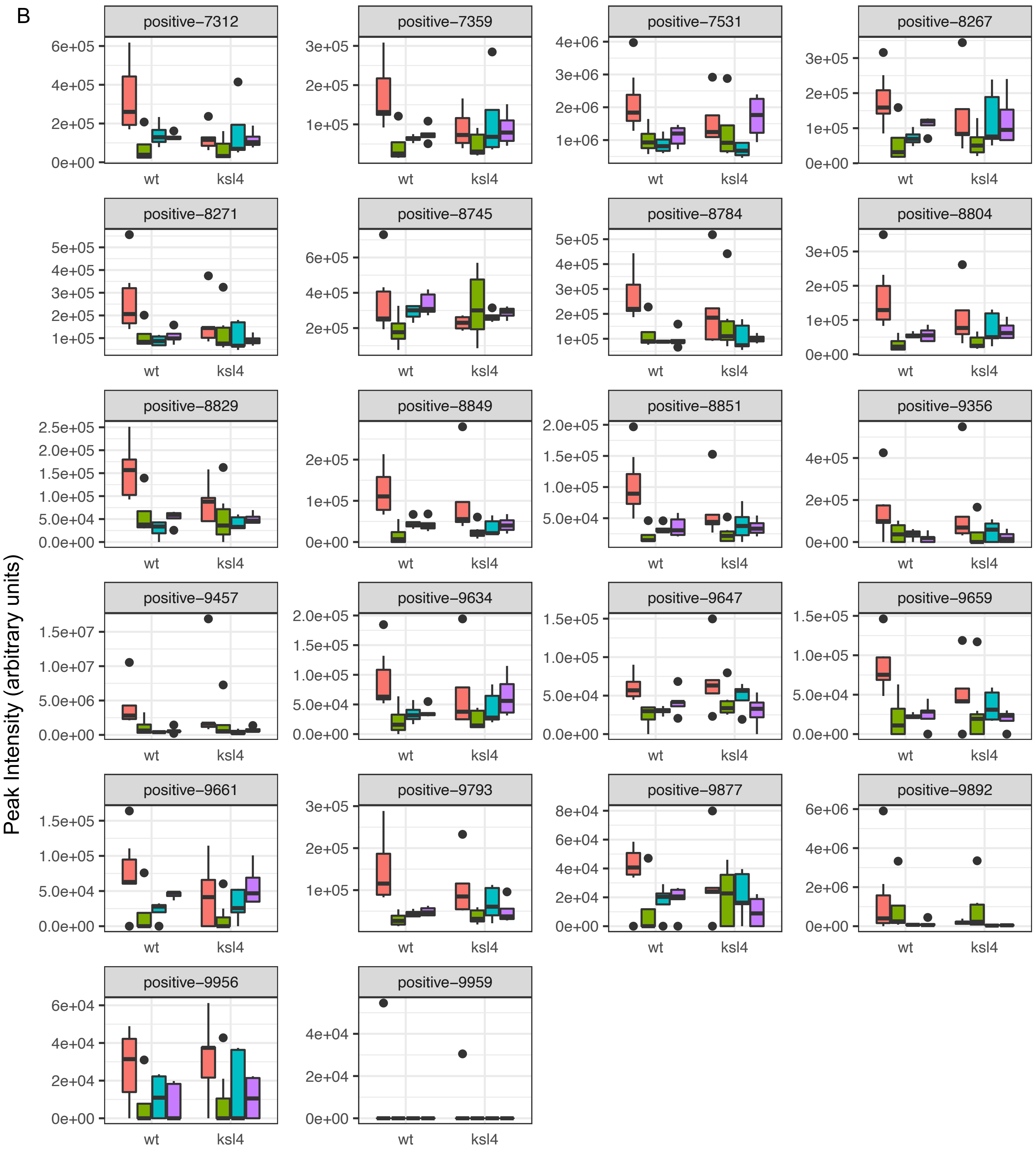


Supplemental Figure 3: Boxplots of feature abundance of DA candidates (i.e. the features commonly enriched in V12 WT root samples for which MS2 were available) in (A) root samples and (B) leaf samples.

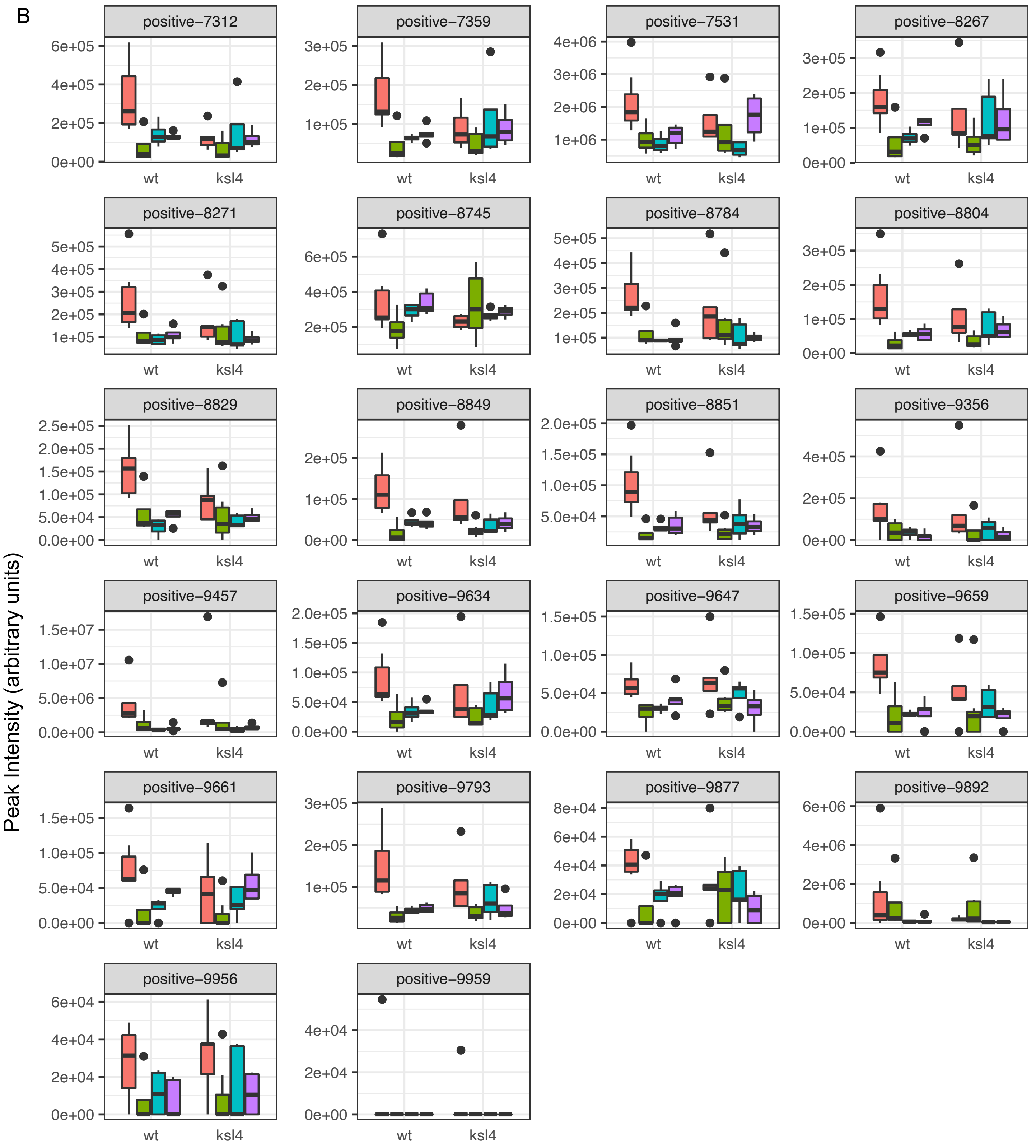


Supplemental Figure 3: Boxplots of feature abundance of DA candidates (i.e. the features commonly enriched in V12 WT root samples for which MS2 were available) in (A) root samples and (B) leaf samples.
