## Supplemental Figure 4 for "A dolabralexin-deficient mutant provides insight into specialized diterpenoid metabolism in maize (*Zea mays*)"

Supplemental Figure 1: NMR Analysis of Dolabradienol

(A) Table of $^{1}H$ chemical shifts dolabradienol (800 MHz chloroform-*d*, 303.3 K)

| | dolabradienol | | | |
|---|---|---|---|---|
| **Label** | **Shift δ** (ppm) | **split** | **# of H** | **J coupling (Hz)** |
| *H1a* | 1.54 | m | 1 | |
| *H1b* | 1.75 | m | 1 | |
| *H2a* | 2.02 | m | 1 | |
| *H2b* | 1.56 | m | 1 | |
| *H3a* | 4.33 | m | 1 | |
| *H3b* | - | - | - | |
| *H6 (a)* | 1.75 | dq | 1 | |
| *H6b* | 1.53 | m | 1 | |
| *H7a* | 1.46 | m | 1 | |
| *H7b* | 1.2 | m | 1 | |
| *H8* | 1.25 | m | 1 | |
| *H10* | 0.93 | ddt | 1 | 12.35 |
| *H11a* | 1.57 | m | 1 | |
| *H11b* | 1.02 | dd | 1 | |
| *H12a* | 1.56 | m | 1 | |
| *H12b* | 1.19 | m | 1 | |
| *H14a* | 1.34 | q | 1 | 17.02 |
| *H14b* | 0.96 | dt | 1 | 12.94, 1.58 |
| *H15* | 5.8 | dd | 1 | 17.5, 10.7 |
| *H16a* | 4.89 | ddt | 1 | 17.52 |
| *H16b* | 4.82 | ddt | 1 | 19.79 |
| *H17* | 1.04 | s | 3 | |
| *H18a* | 4.83 | m | 1 | |
| *H18b* | 4.78 | m | 1 | |
| *H19* | 1.25 | t | 3 | 6.81 |
| *H20* | 0.79 | s | 3 | |
| *3-OH* | 4.11 | s | 1 | |

(B) Table of corrected $^{1}H$ shifts for previously published dolabralexins (800 MHz chloroform-*d,* 303.3 K)

| | *dolabradiene* | | | | *epoxydolabranol* | | | | *epoxydolabrene* | | | | *trihydroxydolabrene* | | | |
|---|---|---|---|---|---|---|---|---|---|---|---|---|---|---|---|---|
| **Label** | **Shift δ** (ppm) | **split** | **# of H** | **J coupling (Hz)** | **Shift δ** (ppm) | **split** | **# of H** | **J coupling (Hz)** | **Shift δ** (ppm) | **split** | **# of H** | **J coupling (Hz)** | **Shift δ** (ppm) | **split** | **# of H** | **J coupling (Hz)** |
| *H1a* | 1.44* | m | 1 | | 1.55* | m | 1 | | 1.41* | m | 1 | | 1.42* | m | 1 | |
| *H1b* | 1.68* | m | 1 | | 1.76* | m | 1 | | 1.65* | m | 1 | | 1.66* | m | 1 | |
| *H2a* | 1.9 | m | 1 | | 2.02 | m | 1 | | 1.88 | m | 1 | | 1.83 | m | 1 | |
| *H2b* | 1.3 | m | 1 | | 1.56 | m | 1 | | 1.27 | m | 1 | | 1.4 | m | 1 | |
| *H3a* | 2.32 | tdt | 1 | 13.7, 5.2, 1.6 | 4.33 | t | 1 | 2.93 | 2.29 | m | 1 | | 4.13 | m | 1 | |
| *H3b* | 2.12 | ddt | 1 | 13.5, 4.2, 1.8 | - | - | - | | 2.1 | dt | 1 | 13.62, 2.15, 2.15 | | | | |
| *H6 (a)* | 1.66 | m | 2 | | 1.74 | dq | 1 | 9.35, 3.04, 3.04 3.04 | 1.63 | m | 2 | | 1.61 | m | 1 | |
| *H6b* | - | - | - | | 1.53 | m | 1 | | | | | | 1.45 | m | 1 | |
| *H7a* | 1.46 | m | 1 | | 1.45 | m | 1 | | 1.2 | m | 1 | | 1.41 | m | 1 | |
| *H7b* | 1.25 | dq | 1 | 13.4, 3.4 | 1.2 | m | 1 | | 1.4 | m | 1 | | 1.15 | m | 1 | |
| *H8* | 1.32 | m | 1 | | 1.25 | m | 1 | | 1.26 | m | 1 | | 1.17 | m | 1 | |
| *H10* | 0.95 | ddt | 1 | 12.5, 2.8 | 0.92 | ddt | 1 | | 0.91 | m | 1 | | 0.81 | m | 1 | |
| *H11a* | 1.56 | m | 1 | | 1.62 | m | 1 | | 1.56 | m | 1 | | 1.52 | m | 2 | |

| | | | | | | | | | | | | | | | | |
|---|---|---|---|---|---|---|---|---|---|---|---|---|---|---|---|---|
| *H11b* | 1.05 | tdt | 1 | 16.0, 15.2, 4.6 | 1.02 | dd | 1 | 10.12, 2.05 | 1.02 | td | 1 | 13.62, 13.62, 4.04 | | | | |
| *H12a* | 1.57 | m | 1 | | 1.65 | m | 1 | | 1.62 | m | 1 | | 1.51 | m | 1 | |
| *H12b* | 1.21 | m | 1 | | 1.18 | m | 1 | | 1.24 | m | 1 | | 0.94 | m | 1 | |
| *H14a* | 1.37 | q | 1 | 14.0, 13.5 | 1.26 | q | 1 | 10.16, 2.16 | 1.26 | m | 1 | | 1.15 | m | 1 | |
| *H14b* | 0.99 | dt | 1 | 12.8, 2.5 | 0.83 | dt | 1 | | 0.83 | dt | 1 | 7.78 | 0.96 | m | 1 | |
| *H15* | 5.83 | ddt | 1 | 17.5, 10.7 | 2.69 | ddt | 1 | 4.00, 3.4 | 2.69 | td | 1 | 3.48 | 3.92 | m | 1 | |
| *H16a* | 4.93 | ddt | 1 | 17.5, 1.4 | 2.66 | ddt | 1 | 4.72, 2.96 | 2.66 | dd | 1 | 4.48, 3.11 | 3.98 | t | 1 | 8 |
| *H16b* | 4.86 | ddt | 1 | 10.7, 1.4 | 2.64 | ddt | 1 | 4.4 | 2.64 | t | 1 | 4.4 | 3.87 | m | 1 | |
| *H17* | 1.02 | s | 3 | | 0.92 | s | 3 | | 0.93 | s | 3 | | 0.8 | s | 3 | |
| *H18a* | 4.487* | t | 1 | 1.5 | 4.81* | m | 1 | | 4.51* | m | 1 | | 4.7* | dd | 1 | 4.5 |
| *H18b* | 4.507* | t | 1 | 1.8 | 4.87* | m | 1 | | 4.52* | m | 1 | | 4.63* | dd | 1 | 5.5, 1.2 |
| *H19* | 1.09 | s | 3 | | 1.5 | s | 3 | | 1.06 | s | 3 | | 1.18 | s | 3 | |
| *H20* | 0.78 | s | 3 | | 0.79 | s | 3 | | 0.75 | s | 3 | | 0.73 | s | 3 | |
| *3-OH* | - | - | - | - | - | - | - | - | - | - | - | - | 4.53 | s | 1 | |

(C) Table of $^{13}$C chemical shifts for dolabralexins, including dolabradienol 800 MHz, chloroform-d for all except dolabradiene (methanol-d), 303.3 K

| Label | dolabradiene annotation | epoxydolabrenol annotation | dolbradienol annotation | epoxydolabrene annotation | THD annotation |
|---|---|---|---|---|---|
| *1* | 21.18 | 16.21 | 16.19 | 21.17 | 16.6 |
| *2* | 28.78 | 34.86 | 34.88 | 28.75 | 35.9 |
| *3* | 33.17 | 74.78 | 74.8 | 33.13 | 73 |
| *4* | 160.93 | 160.98 | 161.05 | 160.84 | 160.3 |
| *5* | 40.69 | 40.35 | 40.4 | 40.64 | 40.5 |
| *6* | 37.95 | 38.73 | 38.79 | 37.88 | 39 |
| *7* | 26.05 | 25.64 | 25.6 | 26.08 | 25.7 |
| *8* | 42.45 | 41.89 | 42.43 | 41.92 | 41.7 |
| *9* | 37.36 | 37.35 | 37.3 | 37.38 | 37.5 |
| *10* | 56.43 | 56.33 | 56.39 | 56.35 | 56.4 |
| *11* | 35.27 | 34.87 | 35.36 | 34.8 | 34.7 |
| *12* | 32.1 | 29.57 | 32.08 | 29.61 | 27.4 |
| *13* | 36.42 | 33.41 | 36.37 | 33.42 | 37.1 |
| *14* | 39.23 | 34.28 | 39.18 | 34.25 | 34.4 |
| *15* | 151.47 | 60.96 | 151.36 | 61.07 | 83.5 |
| *16* | 108.54 | 43.61 | 108.58 | 43.51 | 64.8 |
| *17* | 23.06 | 20.45 | 23.07 | 20.51 | 18.9 |
| *18* | 102.01 | 109.13 | 109.06 | 102.03 | 107.5 |
| *19* | 21.6 | 23.67 | 23.68 | 21.55 | 23.4 |
| *20* | 12.42 | 12.33 | 12.42 | 12.31 | 12.8 |

(D) Dolabradienol numbering and correlations

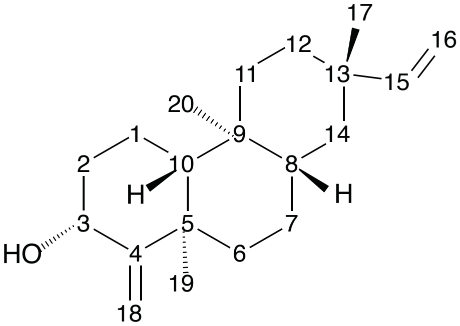


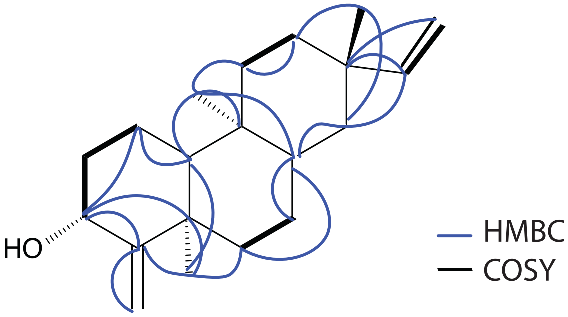


(E) Updated stereochemistries of dolabradiene, epoxydolabranol, epoxydolabrene, and trihydroxydolabrene based on analysis of dolabradienol.

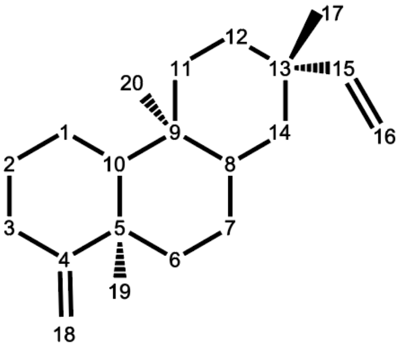

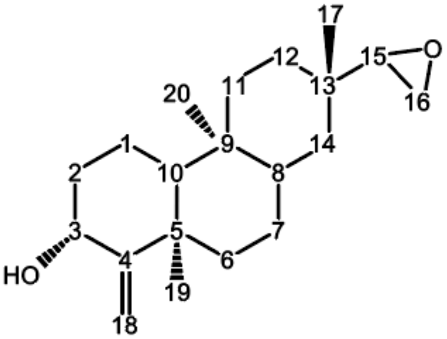


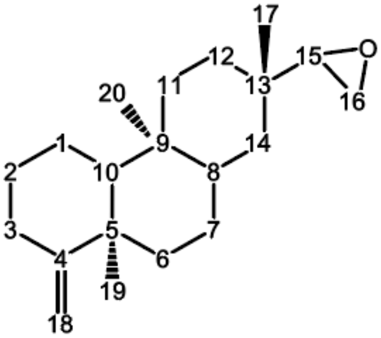

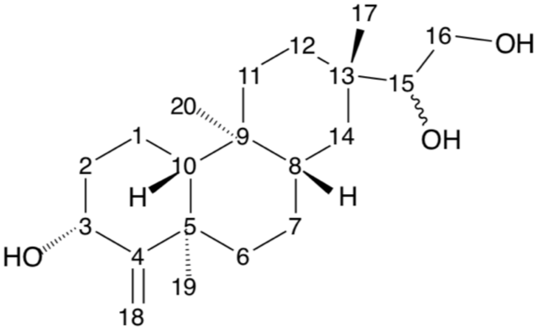

(F) $^1$H spectrum of dolabradienol

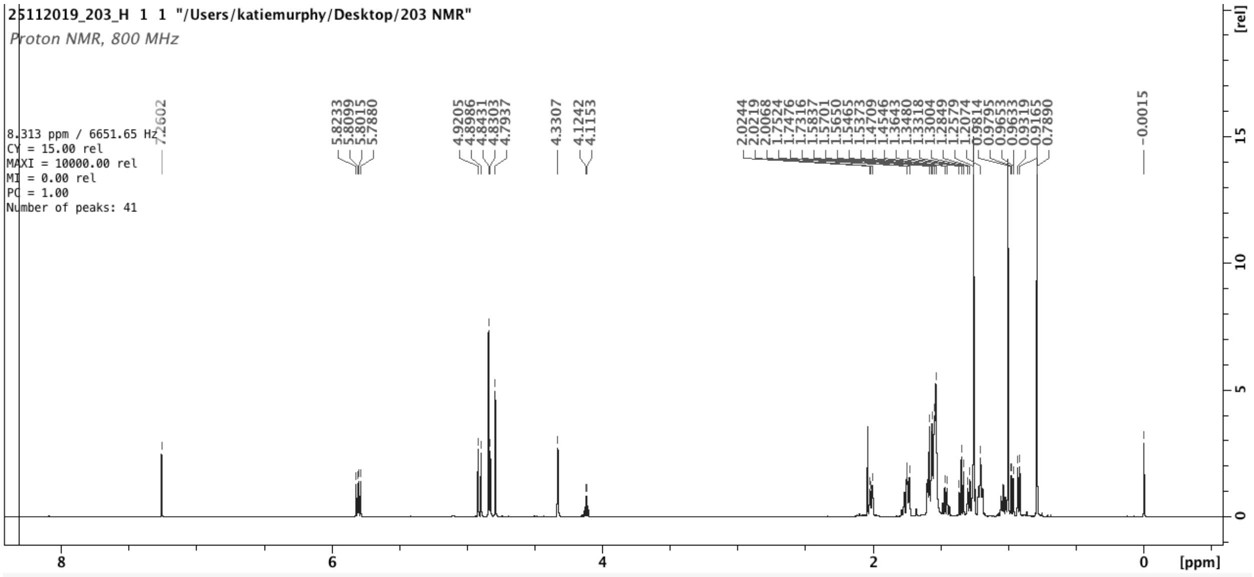


(G) $^{13}$C spectrum of dolabradienol

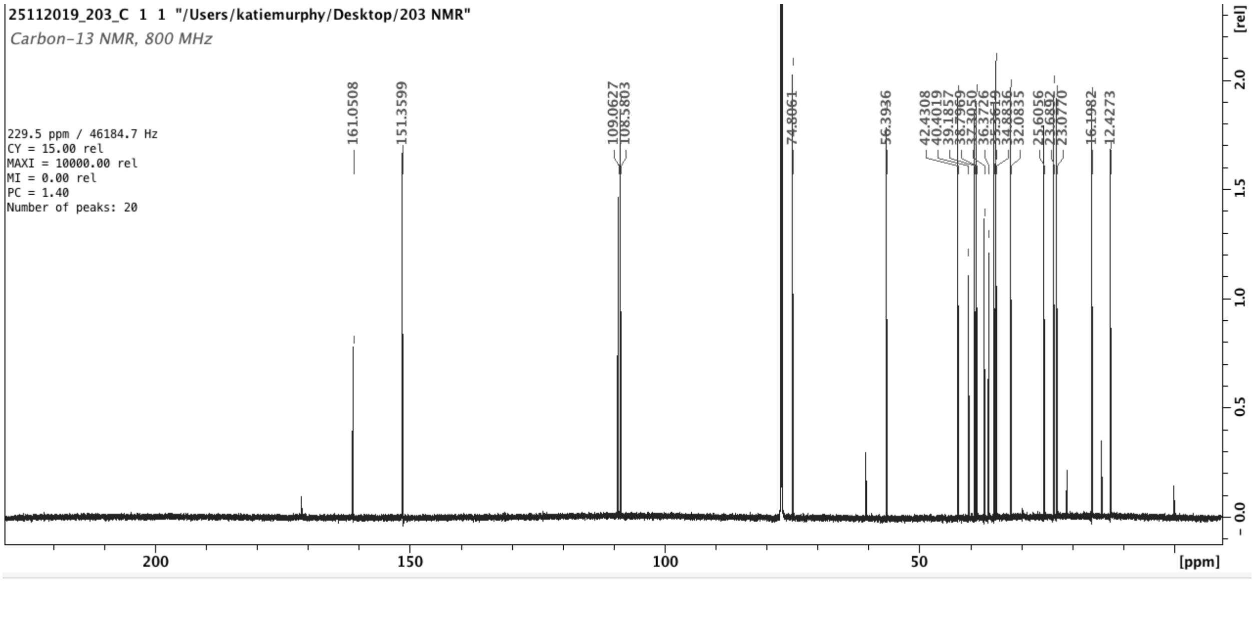

(H) $^{1}$H COSY spectrum of dolabradienol

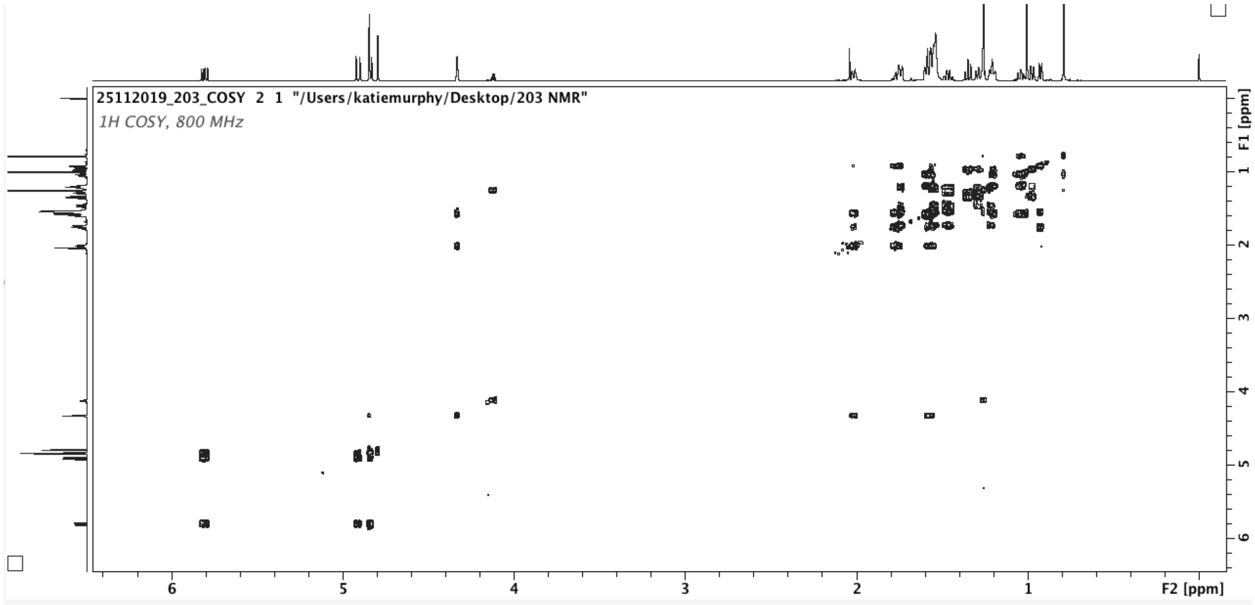


(I) $^{1}$H-$^{13}$C HMBC spectrum of dolabradienol

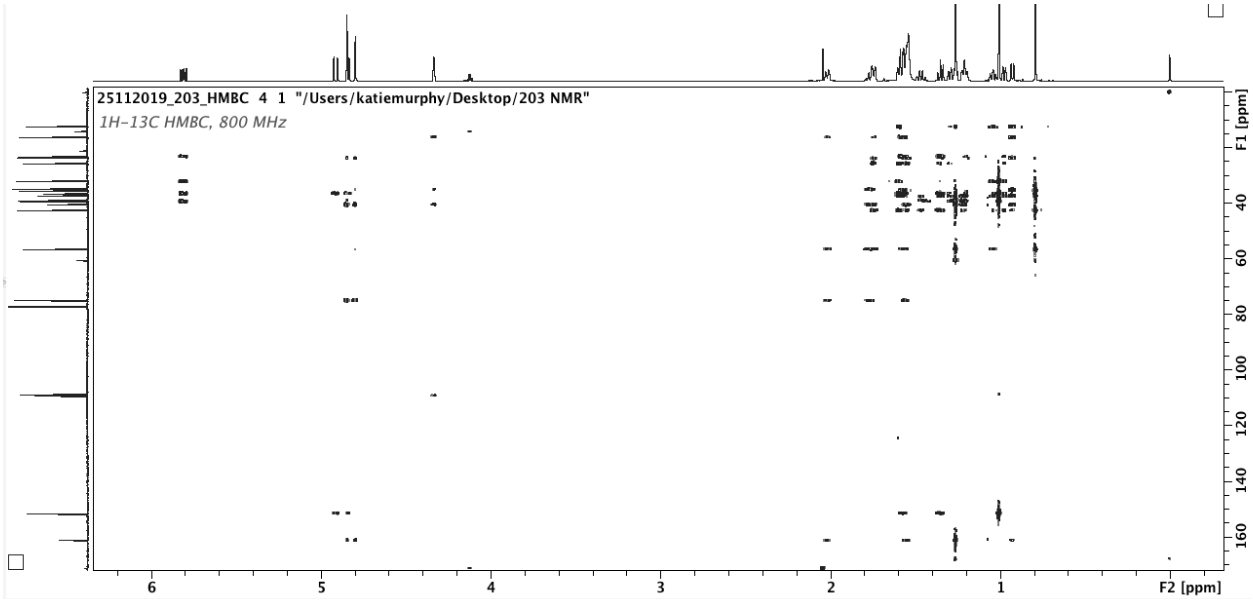

(J) $^{1}H$-$^{13}C$ H2BC spectrum of dolabradienol

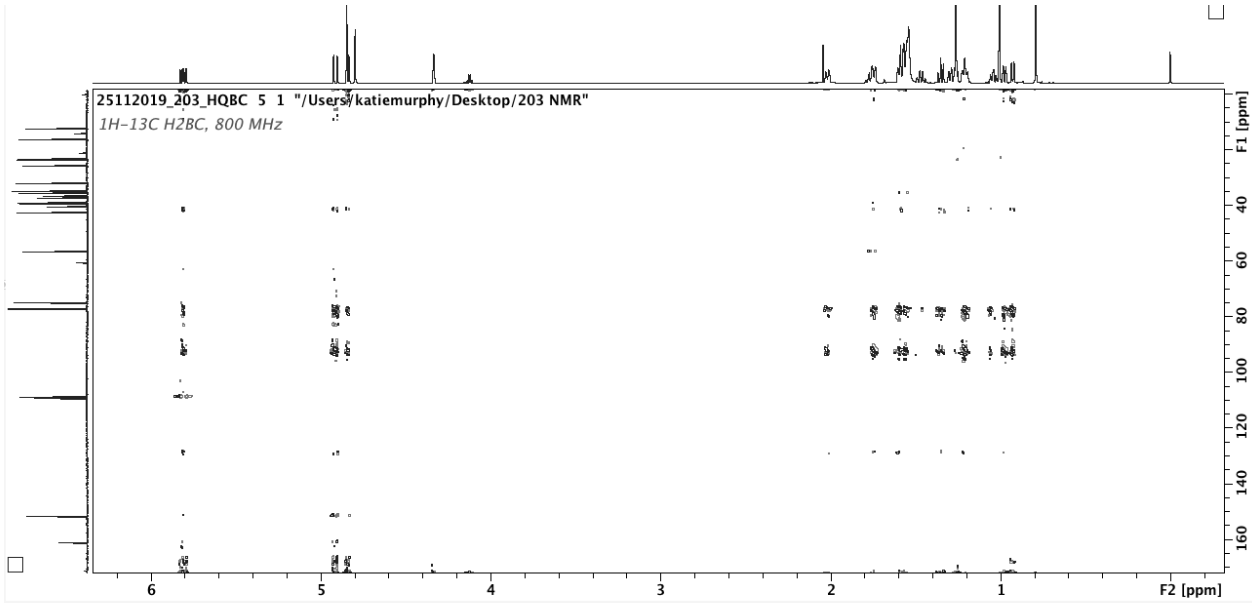


(K) $^{1}H$-$^{13}C$ HSQC spectrum of dolabradienol

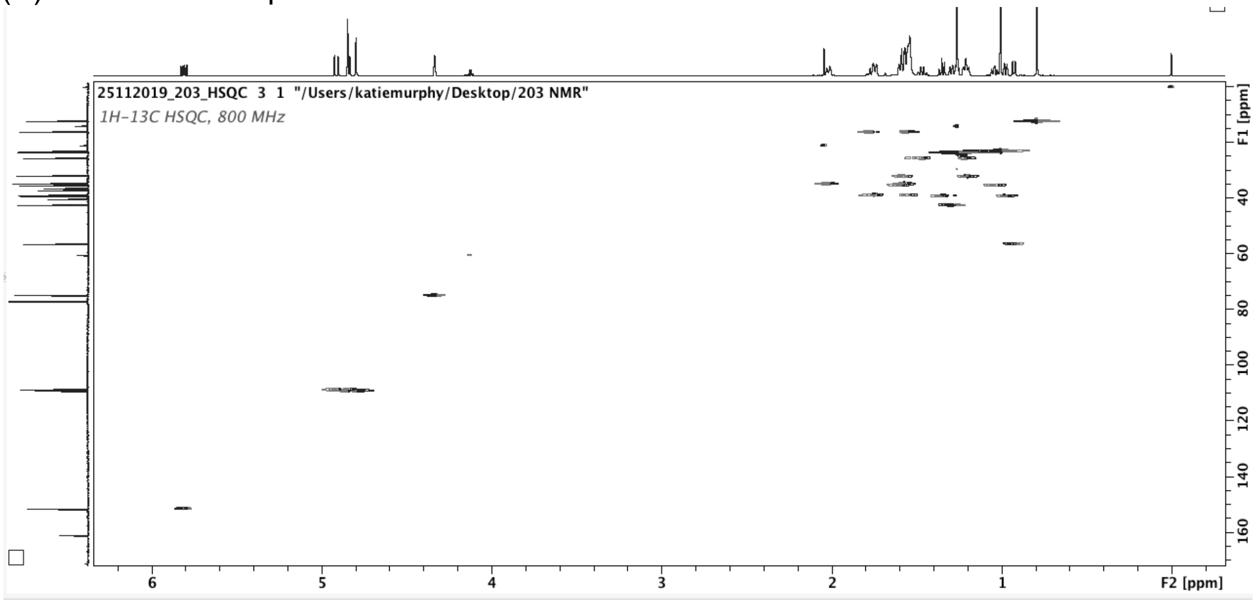
