## Supplemental Figure 5 for "A dolabralexin-deficient mutant provides insight into specialized diterpenoid metabolism in maize (*Zea mays*)"

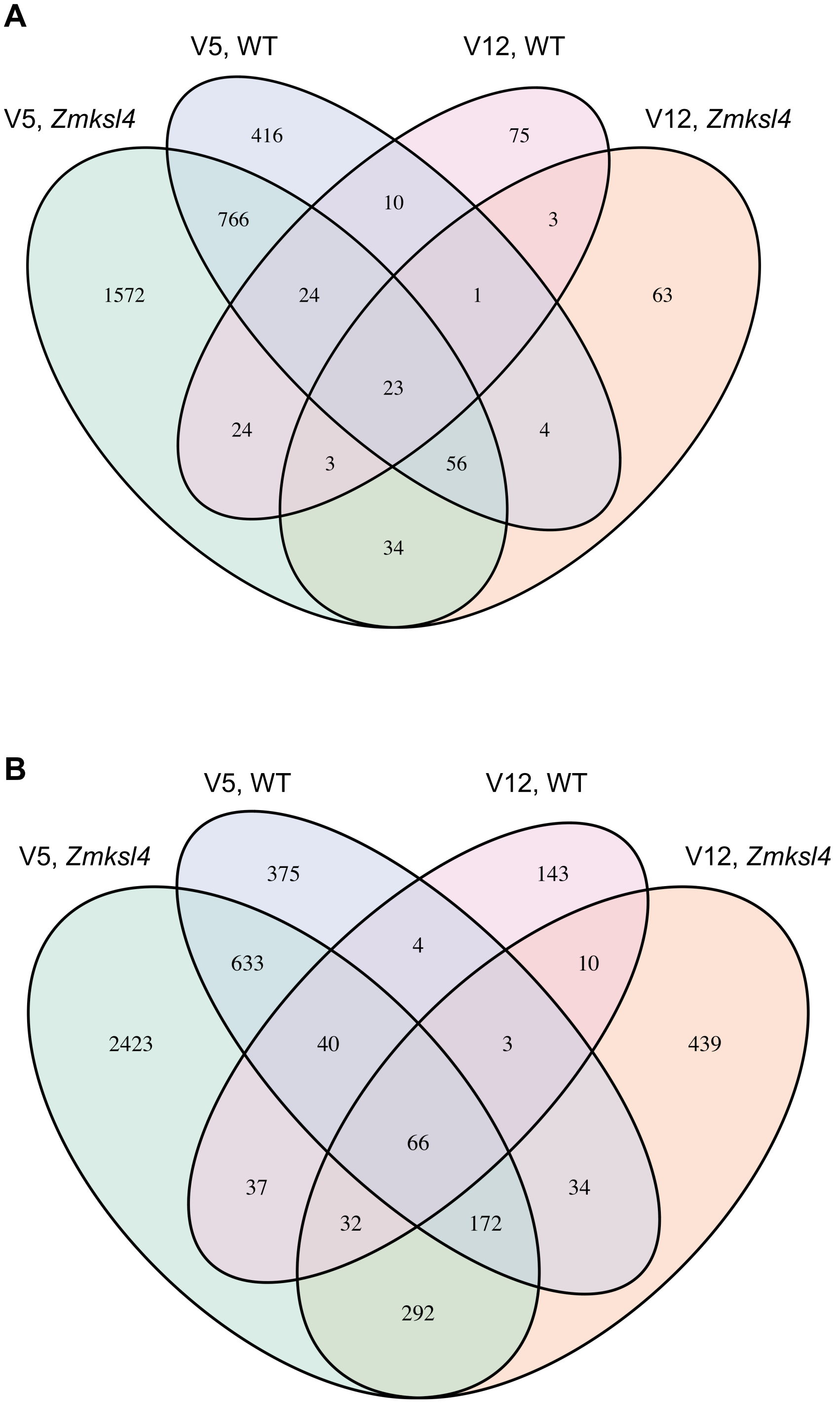


Supplemental Figure 5: Root metabolic features (LC-MS/MS) affected by water status in *Zmksl4* and WT. Few features are commonly affected in both genotypes. Features enriched in (A) water deficit samples and (B) well-watered samples due to water status. Significant features determined by a generalized linear model, P <= 0.1.
