## Supplemental Figure 6 for "A dolabralexin-deficient mutant provides insight into specialized diterpenoid metabolism in maize (*Zea mays*)"

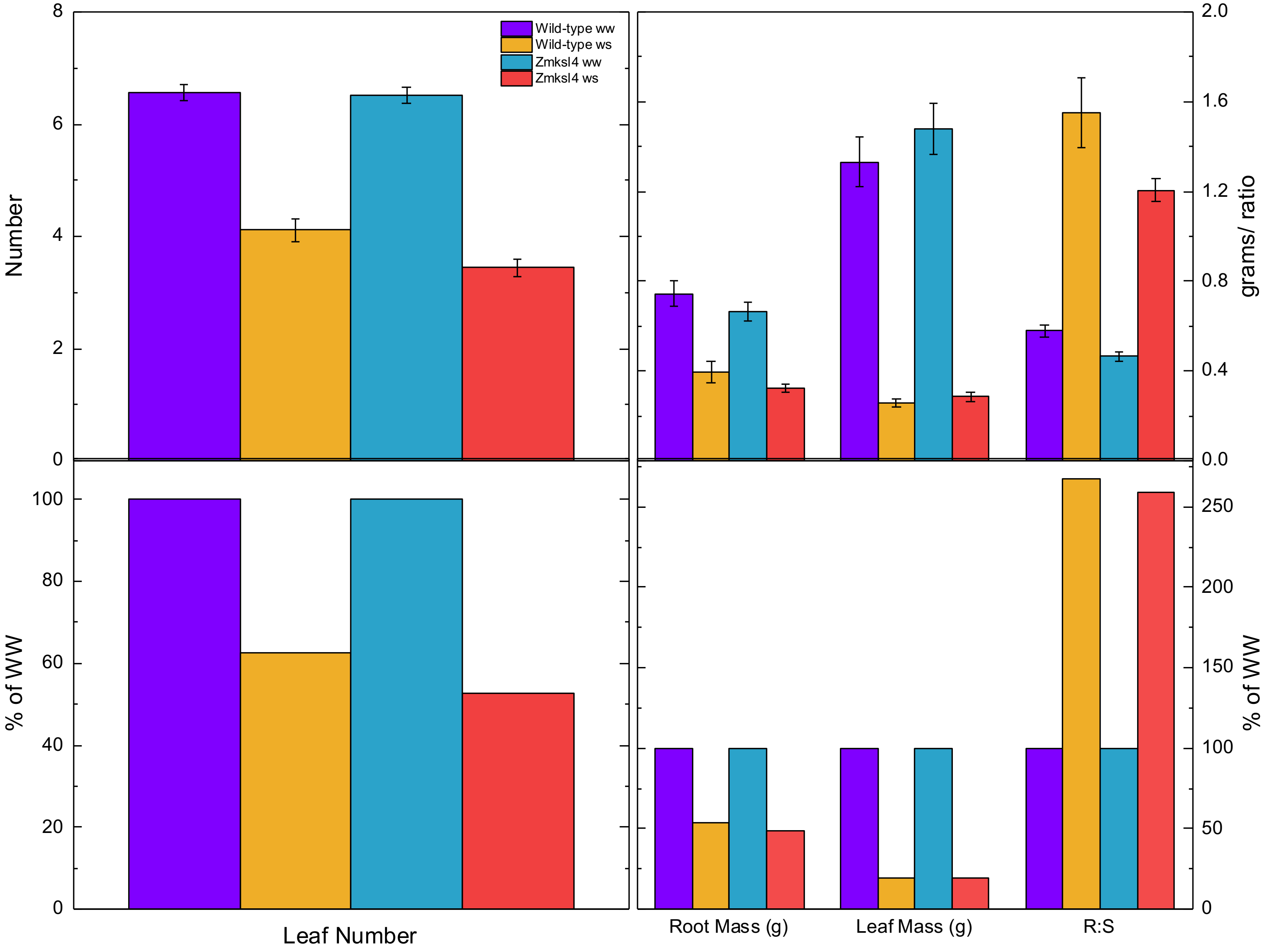


Supplemental Figure 6: Root and shoot weights for plants used in root architecture experiment. WS = water stressed, WW = well-watered, R:S = rot/shoot ratio.
