## Supplemental Figure 7 for "A dolabralexin-deficient mutant provides insight into specialized diterpenoid metabolism in maize (*Zea mays*)"

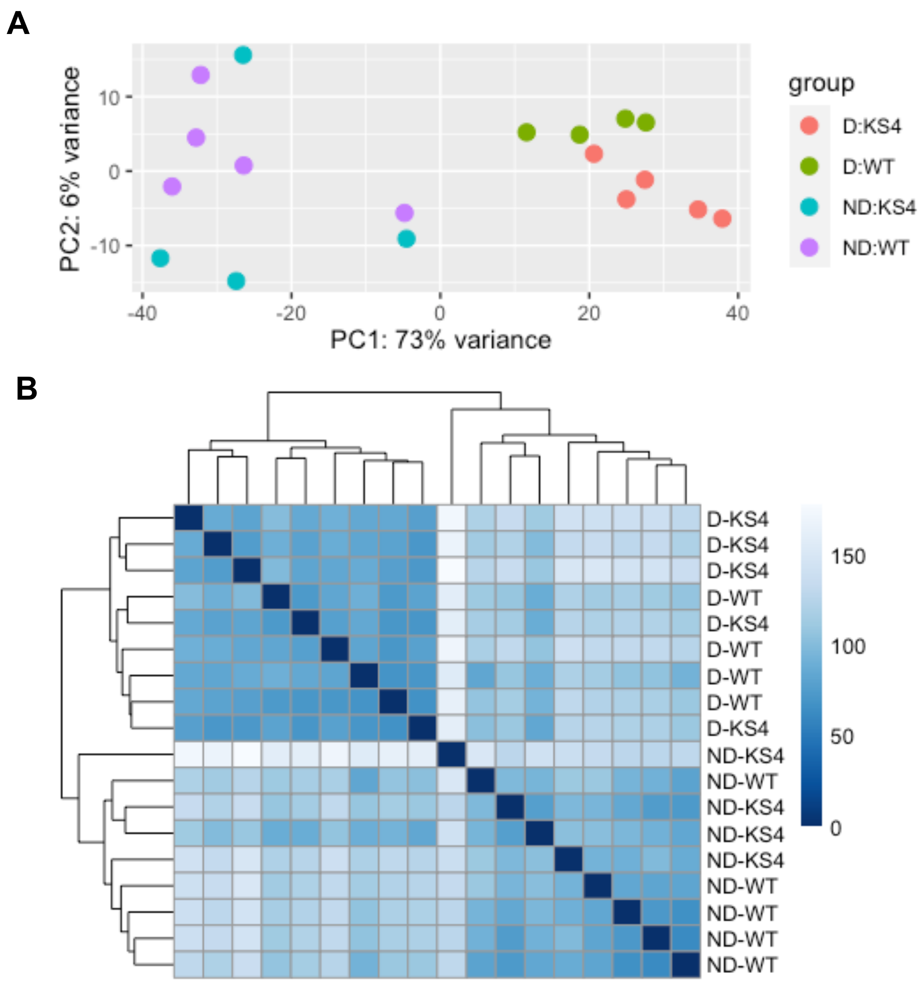


Supplemental Figure 7: Global transcriptome changes in WT and *Zmksl4*. RNA sequencing shows mutation has a minor effect on global transcriptome of *Zmksl4.* (A) Principal component analysis of all sample types. (B) Sample-to-sample distances, each row represents a biological replicate.
